# Forecasting viral antigenic evolution from phylogenetic trees

**DOI:** 10.64898/2026.08.27.741102

**Authors:** Ivan Specht, Soyoon Park, Aakarsh Vermani, Seyone Chithrananda, Claudia L. Driscoll, Garyk Brixi, Julia A. Palacios, Brian L. Hie

## Abstract

Forecasting the evolution of viral proteins can help inform pandemic preparedness and guide the development of interventions. Phylogenetic trees provide historical records of pathogen evolution, which are key for retrospective analysis, but using them to predict future genotypic changes remains challenging. We introduce antiGen, a machine learning model that learns from the evolutionary past of viruses to forecast the evolutionary future. antiGen achieves state-of-the-art performance in forecasting SARS-CoV-2 spike protein mutations according to a variety of metrics, including the prediction of mutations in the receptor binding domain, mutations never previously seen, and mutations with high prevalence. Most experimentally tested antiGen-forecasted spike variants retain pseudoviral infectivity *in vitro*, supporting their functional plausibility. Evaluations on surface proteins of influenza virus, respiratory syncytial virus, and dengue virus further demonstrate applicability across viruses with differing amounts of available sequencing data. Finally, we develop a model trained across 49 viral proteins from 27 viruses, which improves forecasting performance over models trained on individual protein phylogenies. Together, these results support learning from phylogenetic histories as an approach to model viral amino acid changes prospectively, with applications in epidemiology and therapeutic development.

## Introduction

Understanding viral evolution is essential for tracking outbreaks and informing public health response (Grubaugh et al., 2019). Although many genetic changes to viral pathogens are neutral or deleterious, some can alter transmissibility, disease severity, or immune evasion (Sanjuán, 2010; Geoghegan and Holmes, 2018). Forecasting future amino acid changes could help guide the development of vaccines and therapeutics that remain effective as viruses evolve (Łuksza and Lässig, 2014; Fraser et al., 2021). Experimental assays and computational models provide different types of information about viral protein evolution, including expected phenotypic changes and inferred differences in transmissibility. However, predicting the functional effects of an amino acid change, forecasting its occurrence, and anticipating its fitness in the population are distinct tasks. Here, we focus on forecasting future amino acid changes conditional on a starting sequence, including changes that have never before been observed. This task requires learning both the evolutionary constraints on protein sequences and the relative probabilities with which new mutations arise, which may depend on codon accessibility, transition-transversion rates, and the surrounding genetic context.

Phylogenetic trees provide a natural source of information for this task by encoding inferred relationships among sampled sequences. Given a phylogenetic tree, ancestral state reconstruction can then be used to infer the genotypic change along each branch, linking ancestral sequence backgrounds to subsequent amino acid substitutions (Fitch, 1971; Hartigan, 1973; Sankoff, 1975). Previous statistical and machine learning approaches have used phylogenetic information to model sequence changes of natural proteins; see our supplementary note on **Related work** for an overview. Here, we combine information from viral evolutionary histories with modern machine learning methods at the scale of global genomic surveillance to forecast future evolutionary trajectories.

We introduce antiGen, a machine learning model that predicts future amino acid changes by learning from large databases of past ones (**Figure 1**). We first developed a computational pipeline, drawing inspiration from the UShER algorithm (Turakhia et al., 2021), to construct phylogenies containing millions of sampled sequences. From these phylogenies, we extracted training and testing examples, each consisting of a genotype from the inferred evolutionary history paired with the set of single amino acid changes inferred to have altered that genotype. These examples train antiGen to score possible amino acid changes conditional on a starting sequence. We evaluated this formulation using a small Transformer trained from randomly initialized parameters.

**Figure 1.**
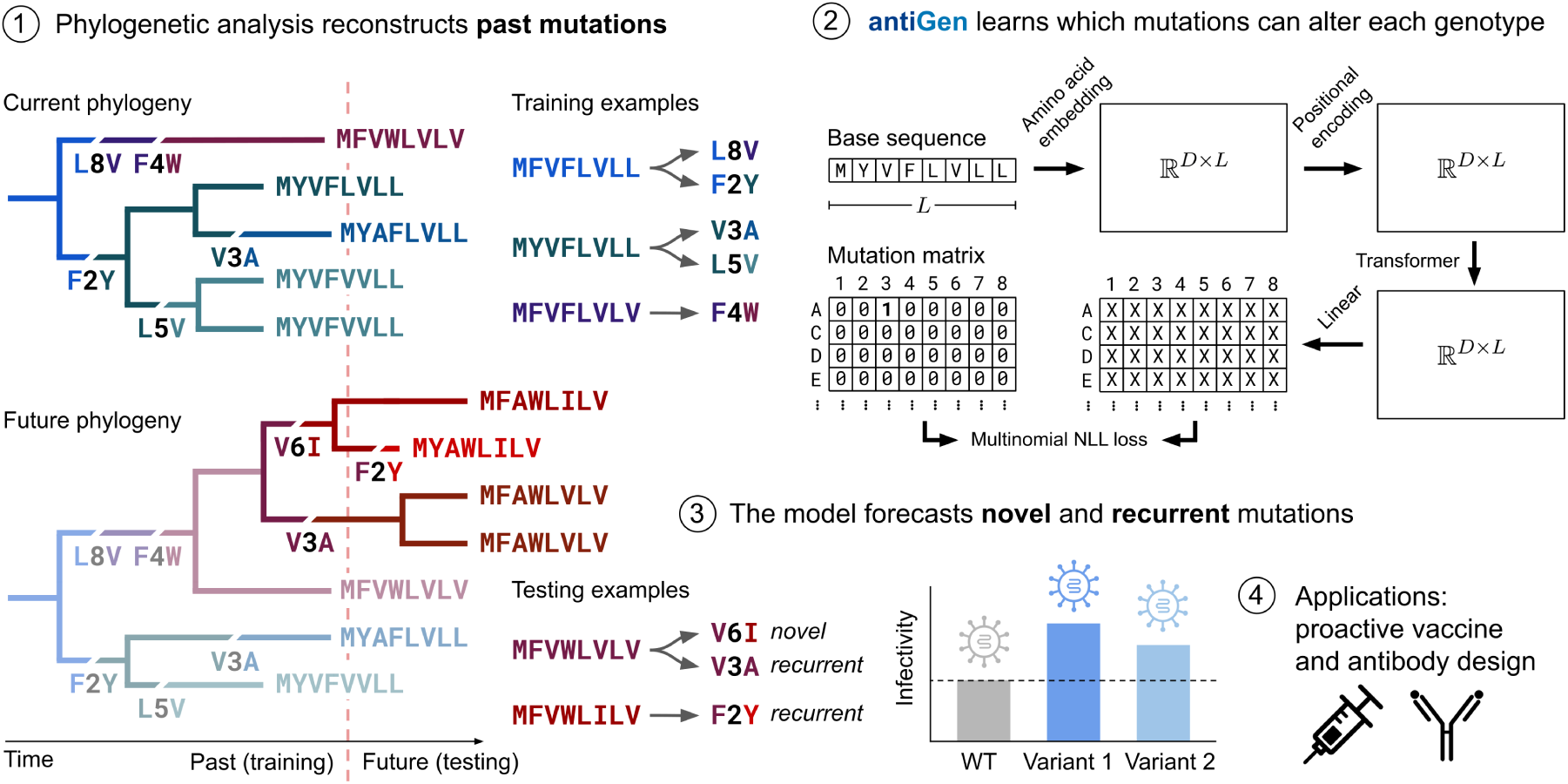
Concept, architecture, evaluation, and applications of antiGen. The goal of antiGen is to predict future mutations to viral proteins based on available genomic surveillance data at present. antiGen is a machine learning method trained on pandemic-scale phylogenetic trees annotated with inferred mutations along each branch. These mutation-annotated trees are partitioned into “examples,” each consisting of a past genotype together with the set of substitutions to have altered that genotype. antiGen then learns the probability of each possible mutation being the next to emerge, conditional on the starting sequence, using a likelihood-based training objective. We evaluate antiGen computationally and experimentally by assessing its ability to anticipate novel and recurrent mutations outside of its training data, and to propose mutations that remain compatible with pseudoviral infectivity *in vitro*. Mutation forecasts can be applied towards proactive vaccine and antibody design, potentially avoiding future outbreaks.

antiGen achieves state-of-the-art performance across comparisons with a wide range of previously reported approaches, including experimental deep mutational scans (DMSs) and machine learning models. We also introduce statistical models trained on the same data as antiGen that outperform existing methods and approximately match antiGen in certain scenarios, such as data-limited regimes. In temporal evaluations of SARS-CoV-2 spike evolution, antiGen outperformed comparator methods in forecasting amino acid changes in the receptor-binding domain across a range of train-test splits. antiGen also performed strongly in forecasting novel, high-prevalence mutations from only limited training data, which is an important public health task early in a pandemic. Combining antiGen with existing baselines further improved predictive performance for some temporal splits, particularly in the beginning of the pandemic. Most experimentally tested SARS-CoV-2 spike variants forecasted by antiGen, including those carrying previously unseen changes, retained pseudoviral infectivity, supporting their functional plausibility.

Beyond SARS-CoV-2, we applied antiGen to several other viral systems and extended it to train across multiple phylogenies simultaneously. Using the same model architecture as for SARS-CoV-2, antiGen achieved leading or competitive performance relative to existing approaches and our statistical models on influenza virus, respiratory syncytial virus (RSV), and dengue virus. We further improved performance by combining antiGen’s training data and objective function with the pretrained protein language model ESM-2 (Lin et al., 2023), enabling joint training of a hybrid model across 49 proteins from 27 viruses. Together, these findings support the use of phylogeny-informed models to forecast viral sequence evolution, with potential applications in vaccine and therapeutic research.

## Results

### Data generation and training

To train antiGen on evolutionary changes inferred from genomic surveillance data, we developed an efficient method for constructing large-scale trees. The method takes as input a small “guide tree” annotated with inferred ancestral sequences and internal-node times, such as those available from Nextstrain (Hadfield et al., 2018), together with a large protein multiple sequence alignment (MSA). It sequentially places each sequence from the MSA onto the tree at a location that minimizes the number of additional amino acid changes required (parsimonious placement), while also accounting for genotypic uncertainty at every node. This approach scales to datasets containing millions of sequences.

For each viral dataset, we constructed separate training and full-data trees to define the training and testing examples. The training tree used an archived guide tree available before a specifed train–test cutoff and included only sequences collected before that date. The full-data tree uses a contemporary guide tree and all available sequences. From each tree, we extracted examples consisting of an inferred ancestral protein sequence together with the amino acid substitutions that arose directly from that sequence. We used the examples from the training tree for model training and validation. From the full-data tree, we retained a testing example only if every sampled sequence (leaf of the phylogeny) descending from its ancestral node represented a sample collected after the given cutoff. We discarded all remaining examples from the full-data tree to prevent overlap with the training data (**Figure 1**; **Methods**).

Given a viral protein sequence, which we refer to as the “query,” antiGen predicts a probability distribution over the position of an amino acid change and the the identity of the replacement amino acid. We fit the model by minimizing the negative log likelihood of the observed amino acid changes conditional on their respective query sequences, normalized by the total number of changes (**Equation 1**). Minimizing this objective corresponds to maximum likelihood estimation of the model parameters (**Methods**).

antiGen consists of a small, randomly initialized Transformer (Vaswani et al., 2017). The model represents each amino acid as the sum of a learned token embedding and a learned absolute positional embedding, processes the sequence through two Transformer encoder layers, and applies a final linear projection to produce logits over the amino acid vocabulary at each position. Across the main four viruses we studied (SARS-CoV-2, influenza, RSV, and dengue), the model contains 465,044–564,628 trainable parameters, depending on protein length. We converted the output logits into a joint probability distribution by applying a single softmax function across the entire output matrix, then zeroing out wild-type probabilities (to avoid degenerate mutations in which an amino acid is replaced with itself), and re-normalizing the matrix. Unlike applying the softmax function separately at each position, this normalization allows the total probability assigned to each position to vary, reflecting differences in the predicted propensity for amino acid change across positions (**Methods**).

### Model evaluation

We primarily evaluated antiGen and comparator models using *next-mutation recall*: the fraction of mutations in the testing set that ranked among the top *q*-quantile of predicted mutations for their corresponding ancestral query sequences, with *q* = 10% by default. This metric can be calculated over all possible amino acid changes to a protein or restricted to a subset of interest, such as changes within a particular region, changes accessible through a single nucleotide substitution, and changes never before observed in training (*de novo* mutations). Next-mutation recall can be evaluated on a *per-mutation* basis, in which a mutation that recurs *k* times during the testing window contributes weight *k* to the metric, or a *per-cell* basis, in which each distinct mutation, defined by its genomic site and derived amino acid, contributes equal weight. These variations allow next-mutation recall to assess predictive performance across a range of forecasting tasks.

For SARS-CoV-2, we focused on mutations within the receptor-binding domain (RBD) due to its role in cell infection and immune escape. We calculated next-mutation recall both for all RBD mutations and for the subset of novel mutations carried by at least 100 sequences after the train-test cutoff, a prevalence threshold we considered sufficient to warrant public health interest or concern. For the other viruses studied, and for the model trained jointly across multiple viruses, we computed next-mutation recall over the entire protein of interest.

Following standard practice for evaluating protein language models (Notin et al., 2023), we also assessed the Spearman correlation between model predictions and DMS phenotypic measurements, where available for the query sequence. However, particularly for SARS-CoV-2, models could achieve high correlations by assigning the same scores to every possible amino acid change at a given position, varying only the per-position probability of mutation. We therefore also calculated the Spearman correlation after replacing the rank of each substitution at position *i* with the mean ranking across all amino acid states at that position.

We used these metrics to compare antiGen with existing sequence-based variant-effect prediction methods (Lin et al., 2023; Hopf et al., 2017; Notin et al., 2022; Fraser et al., 2021; Rao et al., 2021; Thadani et al., 2023; Ito et al., 2025; Obermeyer et al., 2022; Bloom and Neher, 2023), predictions derived from DMS measurements (Starr et al., 2022; Yu et al., 2026; Simonich et al., 2026), and baselines requiring neither parameter optimization nor new experimental measurements. These baselines included BLOSUM62 substitution scores (Henikoff and Henikoff, 1992), minimum codon distance (MCD) scores (also referred to simply as the “codon table”), a position frequency matrix (PFM) recording the number of training sequences containing each amino acid at each position (Wasserman and Sandelin, 2004), and a mutation count matrix (MCM) recording the number of times each (position, amino acid) pair occurred as a mutation across the training examples. We applied BLOSUM62 and MCD scores independently at each position. We also derived per-position conservation scores from the PFM by counting observations of amino acids that differed from the query at each site and assigning the resulting score equally to all possible substitutions at that site (referred to as “PFM conservation”).

To assess whether antiGen captures information beyond site-specific variability, general amino acid replacement preferences, and past allele frequency, we fitted several statistical models to the same data. These included an amino acid substitution matrix applied independently across sites, comprising a learned mixture of BLOSUM62 scores and a codon substitution model (“Codon-B62 mixture”); two models that treat the site of an amino acid change and the identity of the derived amino acid conditional on the ancestral one as independent (“Site-by-substitution”, abbreviated S×S, and “Site-by-Codon-B62 mixture”, abbreviated S×M); and an interpolation of each factorized distribution with the MCM (“S×S– MCM interpolation” and “S×M–MCM interpolation”). Complete definitions and implementation details for the evaluation metrics, baselines, and statistical models are provided in **Methods**.

### SARS-CoV-2 mutation forecasting

Given the impact of SARS-CoV-2 evolution on public health planning, we first evaluated antiGen’s ability to forecast amino acid changes in the SARS-CoV-2 spike protein using a train–test cutoff of January 1st, 2023. The training dataset comprised 7,813,585 sequences and 244,460 inferred spike amino acid changes. To construct testing examples, we built a full-data tree using 8,502,996 publicly available sequences that passed our filtering protocol as of March 22nd, 2026. This tree contained 273,311 total inferred changes, of which 57,319 were part of the testing examples. At the top-decile prediction threshold, antiGen achieved the highest next-mutation recall among the methods evaluated on the RBD, exceeding every comparator by at least 7.9 percentage points under per-cell pooling and 5.5 percentage points under per-mutation pooling (**Figure 2C, D**).

**Figure 2.**
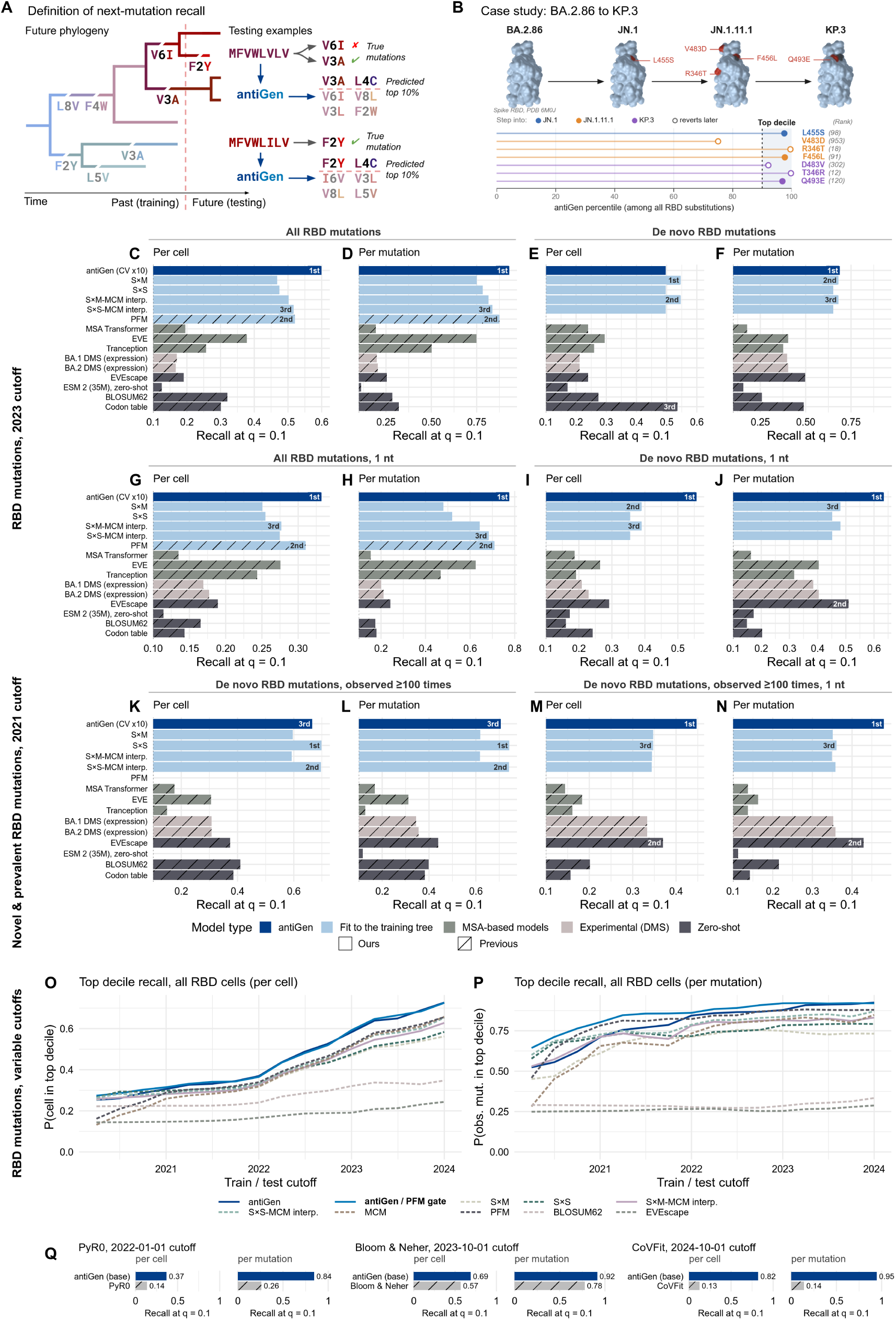
SARS-CoV-2 mutation forecasting. *Caption on the next page. Caption for* Figure 2. (**A**) Definition of next-mutation recall: for each query, we rank all predicted mutations from highest to lowest probability, and calculate the proportion of realized mutations to fall within the top decile of predictions relative to their respective queries. (**B**) antiGen’s rankings of the spike RBD mutations along the evolutionary trajectory from the BA.2.86 variant to the KP.3 variant based on training data through January 1st, 2023 (months prior to the emergence of BA.2.86). All but one appeared among the top decile of predictions. (**C, D**) Next-mutation recall evaluated at quantile *q* = 10% on the RBD based on a January 1st, 2023 train-test split, pooled (C) per cell versus (D) per mutation. (**E, F**) The evaluations in C and D, respectively, restricted to *de novo* amino acid changes, i.e., site-allele pairs never before seen in the training set. (**G, H**) The evaluations in C and D, respectively, restricted to amino acid changes that could be explained by a single nucleotide substitution. (**I, J**) The evaluations in C and D, respectively, with both aforementioned restrictions. (**K, L**) Next-mutation recall evaluated at quantile *q* = 10% on the RBD based on a January 1st, 2021 train-test split, restricted to *de novo* mutations at pandemic frequency ≥ 100 sequences after the cutoff, pooled (K) per cell versus (L) per mutation. (**M, N**) The evaluations in K and L, respectively, restricted to minimum codon distance of one nucleotide substitution, like in (G, H). (**O–P**) Next-mutation recall calculated among all RBD mutations at train-test cutoffs ranging from April 1st, 2020 to January 1st, 2024, pooled (O) per cell and (P) per mutation. **(Q)** Next-mutation recall evaluated at quantile *q* = 10% on the RBD at various train-test cutoffs matched to those of existing published methods.

Statistical models and baselines based on site-independent substitution rules or allele frequencies in the training data also performed well in this evaluation, motivating us to control for these factors. When we restricted the evaluation to *de novo* changes, which by definition cannot be predicted by the PFM, antiGen achieved the highest recall under per-mutation pooling, outperforming the best previous approach by 19% while approximately matching the best statistical model introduced here (**Figure 2F**). Under per-cell pooling, the best statistical models approximately matched the codon table baseline and outperformed the other methods (**Figure 2E**). When we instead restricted the evaluation to mutations with a minimum codon distance of one nucleotide, the codon table performed only slightly better than random, whereas antiGen achieved the highest recall, exceeding the PFM by 2.2 and 6.7 percentage points under per-cell and per-mutation pooling, respectively (**Figure 2G, H**). But when we controlled for both novelty and codon distance, antiGen outperformed existing approaches by a substantially wider margin of 20% under per-cell pooling and 13% under per-mutation pooling. Together, these results support antiGen’s ability to learn patterns of SARS-CoV-2 evolution beyond codon accessibility, per-site conservation, and mutation recurrence.

As a retrospective illustration, we examined antiGen’s rankings of amino acid changes along the evolutionary trajectory from the Omicron sublineage BA.2.86 to KP.3, which was highly prevalent in mid-2024. Using the same January 1st, 2023 train-test split, months before BA.2.86 emerged, antiGen ranked six of the seven changes along this trajectory within its top decile of predictions, including all changes that were not subsequently reversed (**Figure 2B**).

To assess antiGen’s ability to forecast amino acid changes that reach substantial prevalence, we next evaluated recall on RBD changes that never appeared in training and were observed in at least 100 sequences throughout the testing window. This evaluation was designed to match that of EVEscape (Thadani et al., 2023) while allowing models to condition their predictions on a provided query when applicable. Given the public health importance of this task early in a pandemic, when a larger proportion of mutations are *de novo*, we trained antiGen using data through January 1st, 2021, comprising 351,410 sequences and 14,233 inferred spike amino acid changes. After controlling for codon distance, antiGen again achieved state-of-the-art performance, exceeding EVEscape, the second best model, by 7.7 and 5.1 percentage points under per-cell and per-mutation pooling, respectively (**Figure 2M, N**). Without this control, our site-by-substitution statistical model performed best, exceeding EVEscape by 33% and 30%, respectively. Because EVEscape was trained exclusively on prepandemic data, these results suggest that one year of pandemic data would have been sufficient to train antiGen or one of our statistical models to leading performance in predicting novel, prevalent amino acid changes.

We expanded these comparisons to include additional DMS phenotypes, statistical models, and zeroshot baselines (**Figure S1**); antiGen’s ranking relative to other models remained unchanged across the evaluations presented in **Figure 2C–J**. We also compared antiGen with PyR0 (Obermeyer et al., 2022), the model of Bloom and Neher (2023), and CoVFit (Ito et al., 2025) on mutation forecasting using their respective matched cutoff dates. antiGen achieved higher next-mutation recall in every comparison (**Figure 2Q**). Finally, we repeated the evaluations in **Figure 2C–J** across the entire spike protein. Model rankings remained similar, as did the effects of controlling for codon accessibility and mutation novelty (**Figure S2**).

Finally, we assessed agreement between model predictions and experimentally determined phenotypes by calculating Spearman correlations with DMS measurements of BA.2 RBD expression and ACE2 binding (**Figure S3**). antiGen scores conditioned on the BA.2 query sequence correlated positively with both readouts. For antiGen and several statistical models, replacing each substitution’s rank with the mean rank across substitutions at the same position increased these correlations. This finding indicates that position-level constraints can contribute substantially to agreement with these DMS measurements; a high correlation therefore need not imply an ability to distinguish *which* amino acid change will occur at a given position. Corresponding BA.1 DMS measurements correlated more strongly with the BA.2 measurements than did any evaluated computational predictor, although both experimental datasets were generated by the same group using similar systems. This result complements the forecasting evaluations by demonstrating agreement between antiGen’s predictions and experimentally measured effects on RBD expression and ACE2 binding.

### Improving performance over time with gating

In our SARS-CoV-2 analysis, antiGen achieved leading performance in forecasting RBD mutations based on data through January 1st, 2023, outperforming other approaches in most evaluations. Examining a range of different train-test splits from 2020 through 2024, however, we observed that antiGen’s performance relative to other methods depended on the availabililty of sufficient training data. antiGen first surpassed all evaluated baselines at the January 2022 cutoff under per-mutation pooling (**Figure 2P**) and the January 2021 cutoff under per-cell pooling (**Figure 2O**), maintaining its lead through the final cutoff on January 1st, 2024. The PFM remained particularly competitive during 2021 under permutation pooling, consistent with the contribution of recurrent amino acid changes to this evaluation. These findings motivated us to combine antiGen’s predictions with empirical amino acid frequencies to improve forecasting across different levels of training-data availability.

We therefore developed a *gated* predictor that combines antiGen and PFM scores using learned, position-specific weights (**Methods**). Under per-mutation pooling, the gated predictor outperformed both components at nearly every evaluated cutoff (**Figure 2M**). Under per-cell pooling, it outperformed both components at the earliest cutoffs and approximately matched antiGen thereafter (**Figure 2L**). Across both pooling schemes, the gated predictor outperformed every other evaluated methods at each cutoff from October 1st, 2020 onward, with occasional exceptions for standalone antiGen. Its clearest gains over standalone antiGen occurred at earlier cutoffs, when less training data was available. This improvement also extended to forecasting mutations observed at least ≥ 100 times during the testing window using the January 1st, 2021 cutoff, where the antiGen-PFM gate surpassed both antiGen alone and the statistical models (**Figure S4A, B**).

We next examined how training-data availability varied across the SARS-CoV-2 cutoffs and related to forecasting performance (**Figure S5**). Because antiGen learns from inferred evolutionary changes, we reported both the number of inferred amino acid change events and the number of sampled sequences at each cutoff. At the April 1st, 2020 cutoff, the training tree contained 28,622 sequences and 1,129 inferred changes. Although standalone antiGen outperformed EVEscape and BLOSUM62 at this and all subsequent evaluated cutoffs, the site-by-substitution model (S×S) and its interpolation with the mutation count matrix (MCM) outperformed both antiGen and the antiGen–PFM gate under both pooling schemes. These two statistical models also outperformed both predictors at the July 1st, 2020 cutoff, when the training tree contained 93,535 sequences and 3,312 inferred change events. At the October 1st, 2020 cutoff, where the antiGen–PFM gate began to outperform the statistical models, the training tree contained 6,125 inferred amino acid change events across spike, including 664 in the RBD.

These findings provide an empirical reference for the amount of surveillance data available when the antiGen–PFM gate outperformed our statistical models in this SARS-CoV-2 analysis; they do not establish a minimum sequencing or mutation-count requirement. Data requirements may differ across viruses and surveillance settings. The preferred model configuration also depended on the prediction task. For instance, using the January 1st, 2023 cutoff, standalone antiGen outperformed the gate in forecasting *de novo* mutations (**Figure S1C, D, G, H**), which by definition cannot be anticipated by the PFM.

### Experimental validation of predicted SARS-CoV-2 variants

We next assessed whether amino acid changes predicted by antiGen, including changes absent from the training data, were compatible with spike-mediated pseudovirus entry in the XFG sequence background. We developed an experimental workflow to test whether spike variants carrying these changes retained pseudovirus infectivity (**Figure 3A**; **Methods**) (Dadonaite et al., 2023). We queried antiGen with the sequence of XFG, a major circulating SARS-CoV-2 variant at the time of this study (World Health Organization, 2025), and tested predicted mutations within the spike RBD (**Figure 3B**).

**Figure 3.**
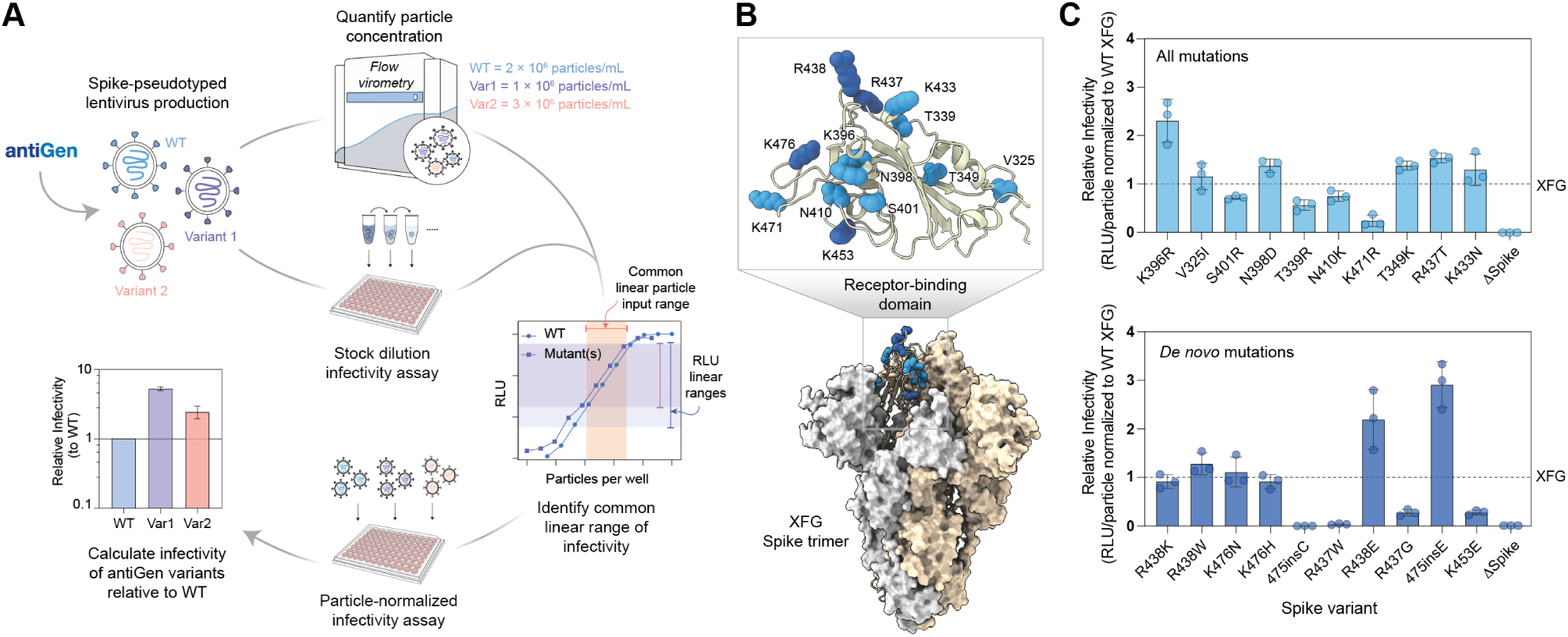
Infectivity of antiGen-predicted SARS-CoV-2 XFG spike variants. **(A)** Experimental workflow for testing antiGen-predicted spike variants. XFG wild-type (WT) and mutant spike-pseudotyped lentiviral particles were produced, quantified by flow virometry, and assayed for infectivity using a luciferase reporter assay following particle normalization. An initial stock-dilution infectivity assay was used to define a shared linear response range across all variants, after which variants were compared at matched particle inputs. **(B)** Structural mapping of antiGen-predicted mutations on an AlphaFold 3-predicted structure of the receptor-binding domain (RBD; residues 312–525) (top) of the SARS-CoV-2 spike trimer (bottom; PDB 8X4H) (Abramson et al., 2024). The ten highest-ranked predicted mutations across all candidate mutations are shown as light blue spheres; the ten highest-ranked *de novo* mutations are shown as dark blue spheres. **(C)** Particle-normalized relative infectivity of XFG spike variants carrying antiGen-predicted RBD mutations, categorized by mutation type as in (**B**). Infectivity was calculated as luciferase reporter signal (relative luminescence units; RLU) normalized to physical particle input determined by side scatter (SSC)-based flow virometry and expressed relative to XFG WT. Relative infectivity of a ΔSpike pseudovirus control, generated by omitting the spike-expression plasmid, is included as a spike-negative pseudovirus reference for background signal. Bars represent the mean of independent experiments; points indicate independent measurements; and error bars denote standard deviation. The dashed line denotes XFG WT infectivity.

Because infectivity measurements obtained using equal volumes of pseudovirus supernatant can be confounded by differences in particle abundance (Youssef et al., 2025), we quantified each pseudovirus stock by flow virometry (Arakelyan et al., 2013; Maltseva and Langlois, 2022). We then compared variants at matched physical particle inputs within a linear operating window shared across the panel of variant pseudoviruses (**Figures S6**, **S7**, **S8**, **S9**, **S10**; **Methods**). Here, “particle” denotes a pseudovirussized nanoparticle enumerated by side scatter (SSC)-based flow virometry. We use “infectivity” to denote the luciferase reporter signal (relative luminescence units; RLU) normalized to such physical particle input, which serves as a readout of pseudovirus entry (**Figure 3A**).

Using this workflow, we tested the the ten highest-ranked antiGen predictions across all candidate RBD mutations and the ten highest-ranked *de novo* mutations, defined as mutations not observed in the training data spanning the first five years of the SARS-CoV-2 pandemic (**Figure 3B**). Most tested XFG spike variants retained measurable pseudovirus infectivity, with only two *de novo* variants exhibiting severe functional defects (**Figure 3C**). Individual mutations produced a broad spectrum of phenotypes, from severe loss of infectivity to levels exceeding those of the XFG wild type. Six of the ten top-ranked overall predictions and four of the ten top-ranked *de novo* predictions showed pseudovirus infectivity comparable to or greater than that of the XFG wild type. The *de novo* predictions showed greater variability, including one variant with a greater than two-fold increase in infectivity and two with no detectable infectivity.

These results show that most tested antiGen-predicted XFG spike variants retained pseudovirus infectivity, including those carrying amino acid changes absent from the training data. While our *in vitro* system does not capture all determinants of epidemiological fitness, including live-virus replication capacity and pathogenicity or antigenic effects that contribute to antibody escape (Carabelli et al., 2023), these findings provide experimental support for the functional plausibility of the tested predictions in the XFG sequence background.

### Forecasting endemic viruses and training across trees

Following our evaluation and experimental characterization of SARS-CoV-2 predictions, we applied antiGen to three endemic viruses of public health importance: seasonal influenza A(H3N2), RSV subgroup A, and dengue virus serotype 2. For each virus, we set the train–test cutoff to the median sample collection date, rounded to the nearest quarter-year: October 1st, 2022 for H3N2; October 1st, 2023 for RSV; and October 1st, 2019 for dengue. We evaluated forecasting performance using next-mutation recall under per-mutation pooling across all amino acid changes in each virus’ surface glycoproteins: hemagglutinin (HA) for H3N2, fusion protein (F) for RSV, and envelope protein (E) for dengue. antiGen consistently matched or outperformed the strongest evaluated baselines and statistical models across all three datasets, both for all changes and for *de novo* changes alone (**Figure 4A–F**). We also scored HA changes along the evolutionary trajectory from the J to K H3N2 variants, which lie on the phylogenetic trunk ancestral to present-day sampled sequences. antiGen ranked every change along this trajectory within its top decile of predictions (**Figure 4G**). However, its gains over our strongest statistical baseline—the interpolation of site-by-mixture with the mutation counts—were modest, motivating us to investigate whether joint training across viruses could further improve forecasting performance.

**Figure 4.**
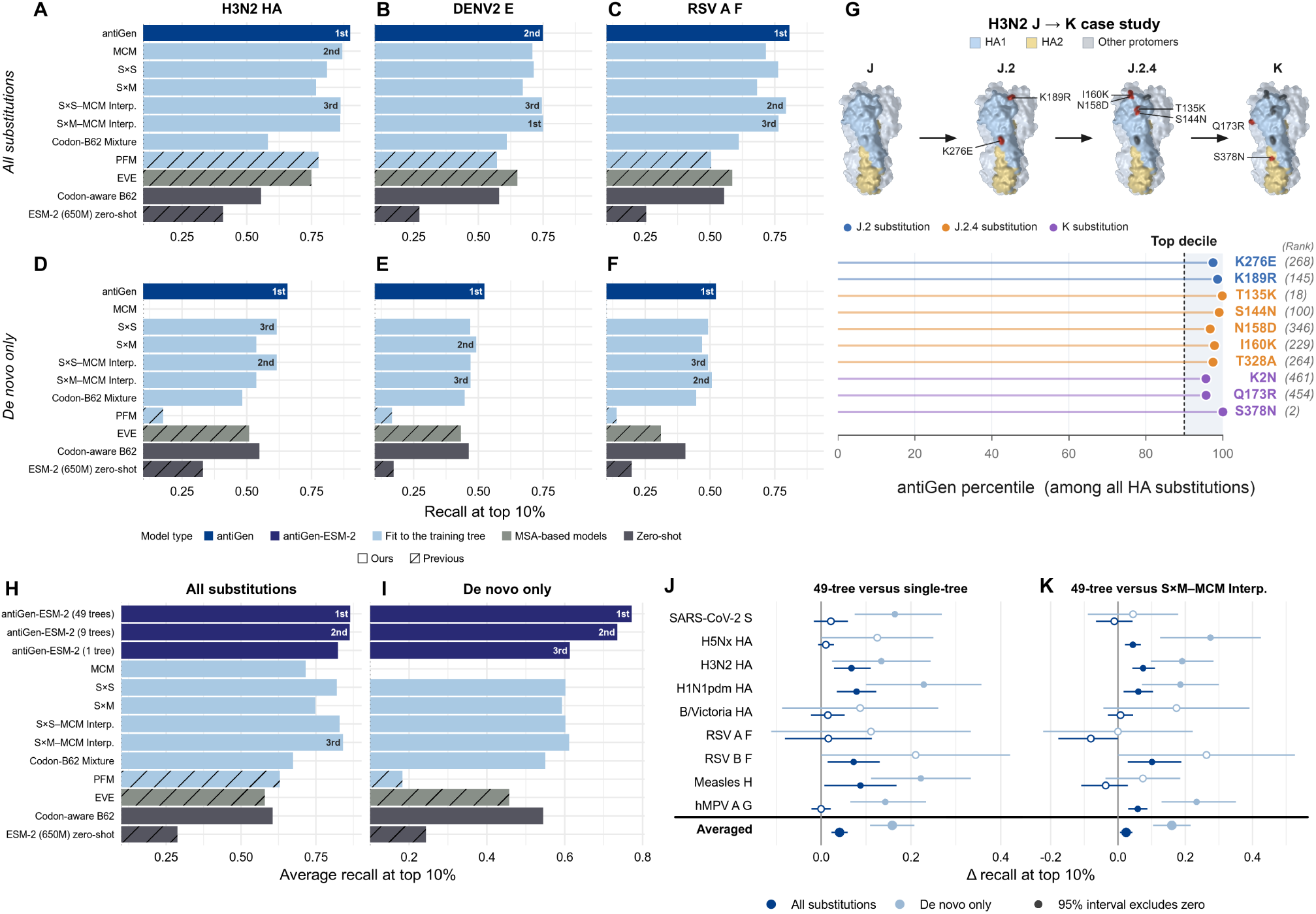
Mutation forecasting for endemic viruses and multi-tree training. (A–F) Next-mutation recall evaluated at quantile *q* = 10% on H3N2 hemagglutinin, dengue envelope protein, and RSV fusion protein. **(G)** antiGen’s rankings of H3N2 HA mutations along the evolutionary trajectory from the J variant to the K variant based on training data through October 1st, 2022 (prior to the emergence of J.2). **(H–I)** Next-mutation recall averaged over a panel of nine viruses such that each virus gets equal weight. **(J–K)** Difference in next-mutation recall on each of the nine viral test sets between the 49 tree antiGenESM-2 and two comparators: single tree antiGen-ESM-2 hybrid and S×M-MCM Interpolation. The difference in recall is reported per virus and as an equal-weight average across viruses. For each virus, whiskers are 95% bootstrap intervals from 2,000 resamples (with replacement) of test events. For the averaged row, each per-virus interval is converted into a standard error, *se_v_* = (high*_v_* −low*_v_*)/2 · 1.96, and these are combined as the standard error of the mean, 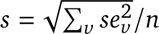, giving the interval Δrecall ± 1.96*s*.

To investigate whether a single model could forecast amino acid changes across multiple viruses, we fine-tuned ESM-2 (Lin et al., 2023) using antiGen’s training data and objective function (**Methods**). We selected ESM-2 because it accommodates variable-length protein sequences, whereas our original antiGen implementation uses absolute positional encodings configured for a single protein. We also hypothesized that ESM-2’s architecture and pretraining on diverse proteins would support learning across viral datasets, motivated in part by its zero-shot performance significantly exceeding uniformly random noise (**Figure 4A, B**). We used two nested training sets: set A contained 9 proteins from distinct viruses, and set B contained 49 proteins from 27 viruses, including all proteins in set A. We applied a common train–test cutoff of January 1st, 2025 to all phylogenies and restricted evaluation to set A’s test data because post-cutoff data were limited for the additional proteins in set B. We omitted MSA attachment for all trees to reduce imbalance in training-data representation across viruses, whose available sequence counts differed substantially—for example, more than nine million for SARS-CoV-2 versus fewer than 16,000 for human metapneumovirus (Shu and McCauley, 2017; Dalla Vecchia, 2024)). Omitting sequence attachment also expanded the set of changes classified as *de novo*, because this designation reflected absence from the resulting training data rater than from all available pre-cutoff sequences.

Fine-tuning the antiGen–ESM-2 hybrid across multiple viral phylogenies yielded higher average forecasting performance than all other evaluated methods, including corresponding models fine-tuned separately on each target protein, with particularly pronounced gains for *de novo* changes (**Figure 4**). Expanding training from nine to 49 viral phylogenies further improved next-mutation recall for both overall and *de novo* changes, suggesting that the additional 40 phylogenies contributed useful training information despite representing proteins outside the evaluation set. In per-virus comparisons, the 49-protein model performed comparably to or better than its single-protein counterpart and the S×M–MCM interpolation baseline across the evaluated viruses (**Figure 4C–D**). Together, these results support the value of training across viral phylogenies and suggest that information learned from additional proteins can improve forecasting for target proteins, particularly for changes absent from their training data.

## Discussion

antiGen predicts future amino acid changes to viral proteins by learning from inferred evolutionary histories. In temporal evaluations of SARS-CoV-2 spike evolution using a January 2023 train-test cutoff, antiGen outperformed a broad range of comparator methods on top-decile recall for RBD changes. Its advantage was particularly pronounced after controlling for codon accessibility and mutation recurrence, factors that the statistical models can exploit to perform well in forecasting amino acid changes at large. Although antiGen required sufficient training data to surpass the PFM on some tasks, including forecasting all RBD mutations regardless of novelty, combining the two through a learned gate generally improved performance over either component. These findings demonstrate antiGen’s value relative to the evaluated comparators for forecasting how a viral protein sequence will mutate next.

antiGen’s training data and objective function can also be paired with a pretrained protein language model to learn evolutionary patterns across viruses. On its own, antiGen matched or modestly exceeded the strongest statistical model, an interpolation of site-by-mixture with a mutation count matrix, in forecasting changes in influenza, RSV, and dengue proteins. Our antiGen-ESM-2 hybrid, co-trained across 49 viral protein phylogenies and evaluated on nine, substantially outperformed both standalone antiGen and our statistical models, particularly for *de novo* changes. More broadly, these results suggest that parent-child relationships extracted from diverse phylogenetic trees can help adapt protein language models to the task of next-mutation forecasting for viruses.

Experimental characterization provided complementary evidence for the functional plausibility of selected predictions. Most tested antiGen-predicted XFG spike variants retained pseudoviral infectivity, including variants carrying amino acid changes absent from the training data. These findings support compatibility with spike-mediated entry in the tested sequence background, but do not directly assess other determinants of epidemiological fitness, including viral replication, pathogenicity, or antibody escape (Carabelli et al., 2023).

antiGen builds on established approaches that use phylogenetic relationships to model evolutionary trajectories and sequence changes (**Related work**). Its contribution lies in combining a scalable pipeline for generating training examples from genomic surveillance data, a conditional prediction objective for fitting a modern machine learning architecture, and temporal evaluation on held-out future amino acid changes. Pairing inferred ancestral genotypes with subsequent changes aligns model training with the forecasting task. Nonetheless, antiGen assumes that its training sequences are minimally diverged from the sequences on which it is queried. This assumption may limit its applicability to novel pathogens with scarce surveillance data.

We do not expect antiGen’s ability to predict single-substitution variants to pose an outsized biosecurity risk. Single amino acid variants preserve high sequence similarity to their parental lineage and are therefore most likely to remain recognizable by homology-based detection methods, such as BLAST (Altschul et al., 1990). Recent advances in nucleic acid synthesis screening also incorporate predicted functional variants to improve detection of modified sequences (Gretton et al., 2025). Nonetheless, sequence detectability alone does not establish the absence of risk, and we acknowledge that biosafety and biosecurity defenses can be imperfect. We recognize the importance of nucleic acid synthesis screening and biological research oversight, as emphasized in recent United States government frameworks and executive actions (National Science and Technology Council, 2024; Exec. Order No. 14,292, 2025). We support continued efforts to strengthen these safeguards and develop responsible biological artificial intelligence (Responsible AI for Protein Design Community, 2023) while enabling important public health research.

From a practical perspective, antiGen could inform interventions against viral diseases by helping anticipate mutations that may escape antibody therapeutics or guide vaccine strain selection to account for anticipated changes. Emerging methods for *de novo* antibody design (Mille-Fragoso et al., 2026) could also be combined with antiGen predictions to proactively develop therapeutic antibodies targeting both historical and future variants of a viral protein.

antiGen’s improved performance with additional training data underscores the importance of making genomic surveillance data widely available to the broader scientific community to support public health modeling and decision-making. Considering its utility for charting the evolutionary trajectory of viral pathogens, antiGen could better equip epidemiological authorities to preempt future variants that pose a serious risk to human health.

### Related work

#### Experimental measurements of amino acid change effects

Deep mutational scanning (DMS) measures the effects of large numbers of amino acid substitutions on specified phenotypes, such as cell entry and antibody binding, under defined experimental conditions (Starr et al., 2022; Taylor and Starr, 2026). These measurements provide valuable functional evidence and are widely used to benchmark computational predictors of variant effects (Notin et al., 2023). However, an experimentally measured functional effect does not directly specify the probability that a change will arise and spread in circulating populations. DMS measurements also do not, by themseleves, account for mutational accessibility, including the number of nucleotide changes required to produce a particular amino acid change. Experimental measurements therefore characterize valuable components of viral function without fully capturing the processes shaping observed evolution (Thadani et al., 2023).

#### Forecasting lineage dynamics and estimating fitness

Established influenza forecasting methods predict future clade or genotype frequencies using information about currently circulating strains (Łuksza and Lässig, 2014; Huddleston et al., 2020). Genealogical tree structure has also been used to predict the success of influenza strains (Neher et al., 2014). For SARS-CoV-2, Obermeyer et al. (2022) use a Bayesian hierarchical model to estimate the contributions of observed mutations to lineage growth. Bloom and Neher (2023) estimate mutation effects on fitness by comparing observed to expected mutation counts on phylogenetic trees. Although this method can score previously unobserved mutations, it assigns them deleterious effects or leaves their fitness effects unresolved by construction. Overall, these approaches primarily forecast changes in the frequencies of currently circulating viral lineages, rather than anticipating new ones—the primary focus of antiGen.

#### Sequence-based models of variant effects

Protein language models learn patterns in amino acid sequences from large training corpora and can score candidate changes relative to a given sequence background (Lin et al., 2023; Hie et al., 2021). Such scores can support variant-effect prediction, but they are not inherently estimates of the probability that a change will subsequently appear in genomic surveillance data. Task-specific adaptation can support more targeted predictions: CoVFit, for example, adapts the protein language model ESM-2 (Lin et al., 2023) to predict the relative epidemiological fitness of SARS-CoV-2 variants (Ito et al., 2025).

Other approaches learn evolutionary constraints from multiple sequence alignments of related proteins. EVE uses a deep generative model trained on these alignments to predict variant effects (Fraser et al., 2021), while MSA Transformer learns representations from aligned sequences (Rao et al., 2021). EVEscape combines evolutionary information with structural and biophysical features to predict viral antibody escape (Thadani et al., 2023). These methods capture evolutionary constraints and functional information relevant to viral evolution, though have been critiqued on the grounds that unequal sequence sampling and training-data composition can bias their predictions (Ding and Steinhardt, 2024; Gordon et al., 2025). antiGen complements these approaches by learning directly from inferred evolutionary changes instead of sequence alignments themselves, making it well suited for the particular task of next-substitution prediction.

#### Learning evolutionary changes from phylogenetic relationships

Phylogeny-based approaches have a substantial history in the modeling of evolutionary trajectories. Inferred changes can be represented in a “mutation tree” (Jahn et al., 2016), which retains relationships among ancestral and descendant genotypes through individual amino acid changes while omitting other details of the phylogeny. Such representations have been used in probabilistic frameworks for modeling evolutionary trajectories (Diaz-Uriarte and Johnston, 2025). TreeMHN, for example, learns dependencies among mutational events and uses them to rank subsequent ones (Luo et al., 2023). These frameworks establish precedents for conditional prediction from evolutionary histories but are based on a fixed set of irreversible events and hence do not directly accommodate the full range of recurrent changes and reversions encountered in viral proteins. Moreover, the largest published applications cover only tens of mutations, far short of pandemic-scale data.

Levinstein Hallak and Rosset (2022) drop the assumption of a fixed mutation set and regress substitution rates on sequence features, including position and neighboring context, to score candidate amino acid changes. However, each predicted rate depends only on a substitution’s immediate nucleotide context, so the model assigns the same score to a given change regardless of the genetic background (beyond neighboring positions). Moreover, because the model is defined on single-nucleotide substitutions, it cannot score amino acid changes that require two or three nucleotide changes within a codon. MutaGAN (Berman et al., 2023) trains a sequence-to-sequence generative adversarial network (GAN) on parent–child sequence pairs from an influenza H3N2 hemagglutinin phylogeny, providing a precedent for modeling viral sequence changes from inferred evolutionary relationships with deep learning. Its utility for predicting specific mutated residues is limited, as the authors report no overlap between the positions most frequently mutated in generated sequences and the most variable positions in their validation set.

Related approaches have also been developed outside of viral forecasting. In antibody affinity maturation, Matsen et al. (2026) use a Transformer architecture trained on parent-child sequence pairs from B-cell phylogenies to distinguish mutation processes from selection and predict functional effects. Lu et al. (2026) fit a neural network-parameterized continuous-time Markov chain to such pairs, modeling antibody evolution as a continuous process. For protein evolution more broadly, Koehl et al. (2026) learn evolutionary rules from phylogenetic cherries—terminal sequence pairs sharing an immediate common ancestor—and generate evolutionary trajectories.

## Methods

### Phylogenetic tree construction and training data generation

As inputs, antiGen accepts (a) a “guide” protein phylogeny with inferred ancestral states assigned to every node and (b) a protein MSA with a collection date for each sample. For all viruses studied in this paper, we obtained the guide trees from Nextstrain (Hadfield et al., 2018) (dataset IDs provided in **Tables S1, S2, S3**). The Auspice format JSON files for phylogenetic trees available on Nextstrain contain the inferred evolutionary changes along each branch, from which the sequence at each leaf and internal node may be reconstructed. We note that these trees are not strictly bifurcating. The MSA for SARS-CoV-2 was compiled from all available public genomes and downloaded from LAPIS (Chen et al., 2023) on March 22nd, 2026. MSAs for the other three pathogens were obtained by downloading all complete sequences not subject to data embargo from GISAID (Shu and McCauley, 2017; Wallau et al., 2023) collected prior to January 1st, 2026, and aligning and translating them with Nextclade (Aksamentov et al., 2021).

We applied two data cleaning steps: filtering out mis-dated sequences and distinguishing genuine gap tokens in sequences from ambiguous amino acids. To remove mis-dated sequences, for each calendar month spanning the range of collection dates, we computed each sequence’s hamming distance to the tree root, then calculated that month’s 99.99th and 0.1st percentiles. If a month had fewer than 2,000 total sequences, its start and end times were symetrically widened until the number of sequences within the window exceeded 2,000. We then made the upper (99.99th-percentile) bound on Hamming distances non-decreasing in time: working backward from the most recent month, we replaced each month’s bound with the minimum of its own value and the following month’s bound, the rationale being that the most divergent lineage in circulation can only accumulate mutations, so a month whose bound exceeds a later month’s most likely contains mis-dated sequences. A sequence was kept only if its hamming distance from the root lay within the band of some month within three months of its stated collection date (plus 3 substitutions of slack on the upper edge). To handle sequencing ambiguity, end gaps, as well as internal gaps exceeding 10 codons, were treated as missing data and replaced with `X` tokens.

To produce pandemic-scale phylogenetic trees efficiently, we took the concept behind the UShER algoithm (Turakhia et al., 2021) and extended it in several key ways. First, we adopted the amino acid alphabet instead of the nucleotide alphabet. We allow for two options: a standard 20-character amino acid alphabet and a 22-character alphabet consisting of the 20 standard amino acids plus a gap token (`-`) and a stop token (`*`). In both cases, all characters outside of the alphabet were treated as equivalent to missing data (`X`). Second, rather than committing a fixed amino acid state at each site for a newly-created internal node, we instead maintain a set of possible parsimonious states that can be updated over the course of the algorithm. This design decision leads to better parsimony scores while incurring minimal extra computational cost (**Figure S11**).

The implementation is presented as **Algorithm S1**. We begin by applying Hartigan’s parsimony (Hartigan, 1973) to construct the set of parsimonious alleles at each site at each internal node of the Nextstrain backbone tree, based on the sequences observed at the leaves. We then iteratively attach sequences in chronological order. We consider two cases when performing an attachment: first, suppose there exists a current node on the tree such that at every unambiguous site on the sequence, the allele set of the node has cardinality 1 and matches the allele on the sequence. In this case, we skip attachment of the sequence, as it could be attached at the node without incurring any extra mutations and without changing the sets of parsimonious allele states on the tree. If there exists no such node, we compute the number of additional mutations necessary to attach the new sequence to each possible current branch of the tree. We attach on a branch selected uniformly at random among those that minimize the number of additional mutations. We then apply Hartigan’s parsimony again to update the set of parsimonious alleles at each site for each node; in most cases, this only requires a local update around the attachment point, hence avoiding high computational cost. We note that this local update and other optimizations are not reflected in the pseudocode, which is meant to give an intuitive understanding of the attachment procedure.

Our method for producing examples from these augmented phylogenetic trees is provided as **Algorithm S2**. First, a random ancestral state reconstruction is selected from the set of parsimonious ones, following Felsenstein (2004), ch. 6. Then, branches with zero mutations are collapsed, while branches with more than one mutation are exploded into multiple branches each with one mutation. The mutation order among said newly created branches is chosen uniformly at random. The resulting tree has exactly one mutation along each branch, meaning that we can construct examples as pairs of the following: the ancestral state at an internal node, together with the set of mutations across the direct children of that node. Despite the randomness introduced at this stage as well as in the attachment algorithm, we saw that antiGen’s next-mutation recall scores as computed in **Figure 2C–J** remained consistent across different random seeds (**Figure S12**). For a worked example of the full pipeline, see **Figure S13**.

The training data generation pipeline was written in C++17 with the OpenMP library for CPU parallelism (Dagum and Menon, 1998).

### Model setup and loss function

Let *L* denote the length of the amino acid sequence of the protein of interest and let *A* denote the number of tokens in the amino acid alphabet (*A* = 20 by default, or optionally *A* = 22 if gap tokens and stop codons are treated the same as standard amino acids). Let [*A*] := {1, …, *A*} represent the set of tokens and let [*L*] := {1, …, *L*} denote the set of positions on the genome. Adopting statistical notation, we define antiGen to be a function **F***_θ_* : [*A*] *^L^* → [0, 1] *^L^*^×*A*^ indexed by unknown, learnable, high-dimensional real parameters *θ*. The function **F***_θ_* takes in an amino acid sequence, represented as a vector of *L* tokens, and returns an *L* × *A* matrix of probabilities specifying the joint distribution over the position at which an amino acid substitution occurs and the replacement token for the amino acid at said position. We write the probability under the model of replacing the token at position *i* ∈ [*L*] with token *j* ∈ [*A*] in a sequence *q* ∈ {1, …, *A*}*^L^* as **F***_θ_*(*q*)*_i j_*. By convention, we assume that entries of the matrix **F***_θ_*(*q*) that do not represent actual mutations (i.e., (*i*, *j*) for which *j* = *q_i_*, corresponding to degenerate substitutions where the mutant amino acid is the same as the original amino acid) are set to zero.

We define the loss to be the negative log likelihood (NLL) of the mutations in the training data given *θ*, treating each mutation extracted from the phylogeny as an independent observation. Formally, let E denote the set of training examples, where an example *e* ∈ E consists of a query sequence *q*^(^*^e^*^)^ ∈ {1, …, *A*}*^L^* and a matrix 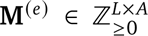 whose (*i*, *j*)th entry 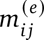 is the number of times a mutation at position *i* to amino acid *j* was observed in training example *e*. As shorthand, for *e* ∈ E, let **F***_θ_*(*e*) ∈ [0, 1] *^L^*^×*A*^ denote the evaluation of **F***_θ_* on the sequence *q*^(^*^e^*^)^ . The loss is then given by

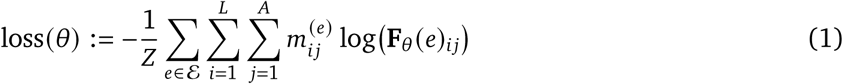

where *Z* is the total number of mutations across all training data, given by

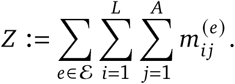

We minimized this loss function via stochastic gradient descent in order to yield an approximate maximum likelihood estimate *θ*^^^ of *θ*.

### Evaluation metrics

Here, we provide mathematical details of the evaluation metrics used to assess antiGen and other models. We continue with the notation established above in **Model setup and loss function**. We define next-mutation recall to be the proportion of mutations across all testing examples that lie in the top *q*-quantile of predicted mutations, for 0 ≤ *q* ≤ 1. To state this quantity mathematically, we first need to handle what membership in the top quantile means in the presence of tied rankings. Let E^′^ denote the set of testing examples, generated per **Phylogenetic tree construction and training data generation**, using the same format as E. For an example *e* ∈ E^′^, a position *i*, and a new amino acid *j*, let 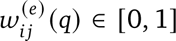 denote the *weight* with which cell (*i*, *j*) belongs to the top *q*-quantile of **F***_θ_*_^_(*e*), the predicted mutations relative to query *e*. Let *n* = *L*( *A*− 1) be the number of entries of **F***_θ_*_^_(*e*), let *ζ*_1−_*_q_* (**F***_θ_*_^_(*e*)) be its ⌈*qn*⌉-th largest entry (the (1 − *q*)-quantile boundary), and let *a* and *t* be the numbers of entries of **F***_θ_*_^_(*e*) strictly greater than and equal to that boundary, respectively. Then

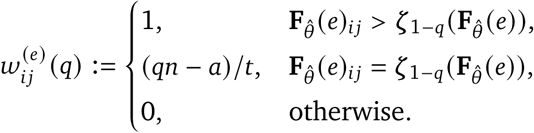

Predictions strictly above the boundary count fully, and those exactly at it share the remainder equally, so that the weights sum to exactly *qn* whether or not there are ties; equivalently, 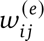 (*q*) is the probability that cell (*i*, *j*) is among the top *qn* predictions when ties are broken uniformly at random and the boundary position is included with probability equal to the fractional part of *qn*. With this definition in place, recall pooled by mutation is given by

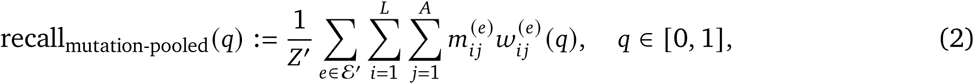

where *Z*^′^ is the total number of mutations across all testing examples, i.e.

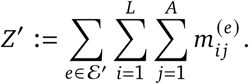

Pooled by cell, the recall equation becomes

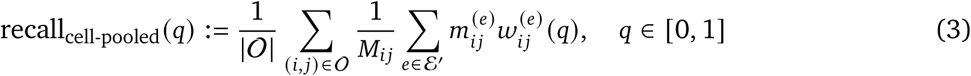

with

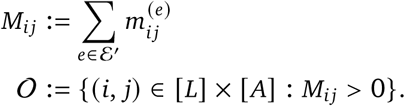

Now, suppose we wish to restrict the recall evaluation to some subset of mutations of interest. Let *S*(*e*) denote the mutations of interest relative to a query *e*. Some examples of a choice of *S*(*e*) used in the paper include:

- RBD mutations for SARS-CoV-2: let *R* ⊆ [*L*] denote the set of sites that consitute the RBD. Then set

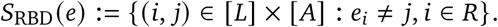

- *De novo* mutations: let PFM denote the position frequency matrix of the training sequences (see **Statistical Models**). Then

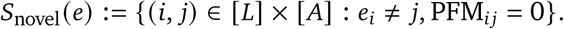
- Mutations that could be explained by a single nucleotide substitution: let *D* denote minimum codon distance. Then

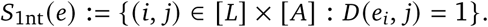

For any such restriction, we may revise the recall definitions as follows. Write *F_i j_* := **F***_θ̂_*(*e*)*_ij_*, and let

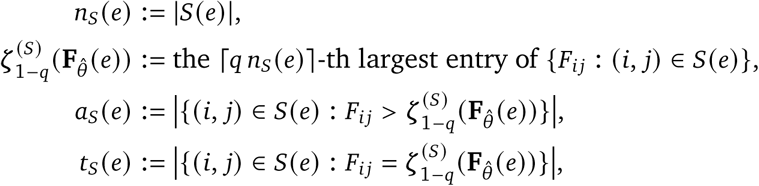

and, for (*i*, *j*) ∈ *S*(*e*),

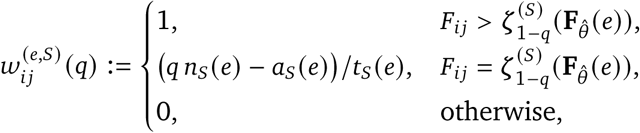

so that 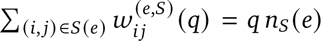. Only mutations of interest count as targets: pooling by mutation, we have

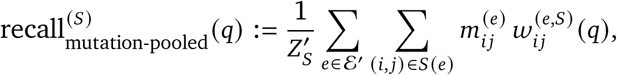

With

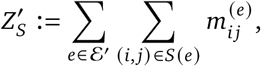

and, pooling by cell

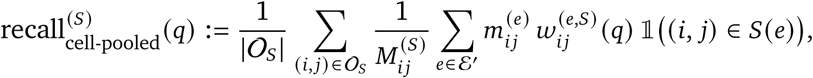

with

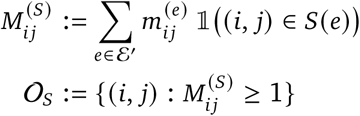

That is, *S*(*e*) restricts both the candidates a query’s predictions are ranked among and the mutations that count as targets. Restrictions compose by intersection; in **Figure 2**, we report *S*_RBD_ (panels C, D), *S*_RBD_ ∩ *S*_novel_ (panels E, F), *S*_RBD_ ∩ *S*_1nt_ (panels G, H) and *S*_RBD_ ∩ *S*_novel_ ∩ *S*_1nt_ (panels I, J). Taking *S*(*e*) to be every non-wild-type cell recovers equations (2) and (3).

All of the restrictions imposed by *S*_RBD_, *S*_novel_ and *S*_1nt_ can be determined based on the training data alone. Some restrictions, such as a mutation being observed in at least 100 sequences after the train-test cutoff date, require knowledge of the testing data, in which case it would constitute leakage to filter the pool of predictions based on such a criterion. In this case, we define another filter *T* that affects only the set of targets, not the set of predictions. For instance, the prevalence-in-testing-window filter would be given by

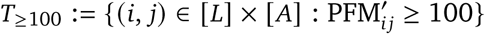

with PFM^′^ the position frequency matrix of all of the sequences collected after the train-test cutoff. Given a restriction *S* on the pool (as before), we define

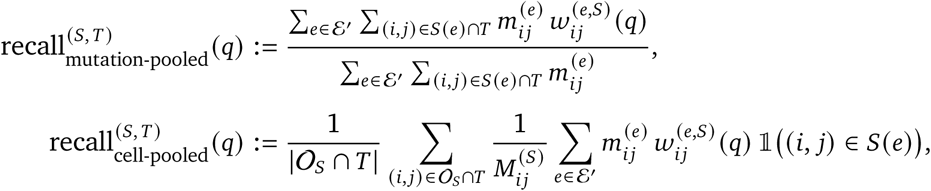

with 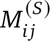 and O*_S_* as above. This metric with *T* = *T*_≥100_ is reported in **Figure 2**, using *S* = *S*_RBD_ ∩ *S*_novel_ (panels **K, L**) and *S* = *S*_RBD_ ∩ *S*_novel_ ∩ *S*_1nt_ (panels **M, N**).

#### Training and fine-tuning

Here, we supply the technical details for training antiGen from scratch and fine-tuning existing models on the antiGen objective (or simply evaluating them as-is). We provide a brief description of each model as well.

- **antiGen** (Transformer trained from randomly initialized parameters). A small encoder-only Transformer (amino-acid token embeddings plus learned absolute positional embeddings; *d*_model_=128, 2 post-norm layers, 4 attention heads, feed-forward dimension 512 with ReLU, dropout 0.1) with a linear head onto the 20-(default) or 22-(if gaps and stops are included) token alphabet at every position. Because it has no pre-training stage, every parameter was initialized at random and trained end-to-end with the loss function of Equation 1 at learning rate 10^−4^ and batch size 64.
- **antiGen-PFM gate.** A learned per-position mixture of two predictors: the antiGen Transformer above (same architecture, trained from random initialization) and the position frequency matrix of the training sequences, scored as log(1 + count). Within each query, both score maps are standardized (z-scored over all positions and amino acids), then combined at each position *i* with weight *g_i_* = *σ*(*γ_i_*) on the Transformer and 1 − *g_i_* on the PFM. Here *γ_i_* is a learned per-position parameter initialized at 0 (equal weighting). The Transformer and the gate are trained jointly with the loss of Equation 1 at learning rate 10^−4^ and batch size 64; the PFM term has no parameters.
- **antiGen-ESM-2 finetune.** ESM-2 is a masked protein language model pre-trained on UniRef50 (Lin et al., 2023). We used the 35M-parameter (`facebook/esm2_t12_35M_UR50D`) and 650M-parameter (`facebook/esm2_t33_650M_UR50D`) checkpoints. We projected the per-position hidden states onto the amino acids using ESM-2’s pre-trained masked-language-model head, with its output logits restricted and reordered to our amino acid alphabet, and then continued training using antiGen’s objective (**Equation 1**). We evaluated three configurations:

**–** *LM head finetune* (learning rate 10^−3^): backbone gradients were disabled; only the head was updated.
**–** *Partial finetune* (learning rate 10^−4^): the last 4 Transformer blocks and the head were unfrozen; earlier layers remained fixed.
**–** *Full finetune* (learning rate 10^−5^): every parameter was trainable.

Batch sizes were 32 (LM head and partial) and 16 (full) for the 35M model, and 4 for every configuration of the 650M model except for those trained for the nine-virus panel, which used a batch size of 8. In **Figure 4** we report only the 650M partial finetune configuration which generally exhbited leading performance among the group.

By default, antiGen is trained 10 separate times, each on its own random 90/10 split of the training examples into training and validation sets, with its own initialization seed. Each model was trained for 256 epochs with a checkpoint saved after every epoch, and the checkpoint with the lowest negative log-likelihood on its validation set was kept (**Figure S14**). At test time, each model’s scores are normalized to log-probabilities over all candidate substitutions, and the ten are averaged (equivalent to a product of experts up to a normalization constant). Epoch selection and cross validation were applied the same way for the antiGen-ESM-2 finetune. We additionally report a single antiGen model trained on all training examples for a fixed 32 epochs, with no validation split; it performed slightly worse than the cross-validated ensemble in 7 of the 8 panels of **Figure S1**. The antiGen-PFM gate was likewise each trained once on all training examples for a fixed 32 epochs.

All trainable models above were optimized with Adam (Kingma and Ba, 2015) (default PyTorch hyperparameters: *β* = (0.9, 0.999), *ε* = 10^−8^, no weight decay), with gradient norms clipped to 1.0, no learning-rate schedule, and full (32-bit) precision. Data loaders used 4 worker processes, pinned memory, and batches shuffled every epoch. The training and evaluation pipeline was written in Python (version 3.12.2) with the following libraries: numpy (Harris et al., 2020), scipy (Virtanen et al., 2020), pandas (McKinney, 2010), matplotlib (Hunter, 2007), biopython (Cock et al., 2009), PyTorch (Paszke et al., 2019), and Hugging Face Transformers (Wolf et al., 2020).

#### Statistical Models

We designed several statistical models to better contextualize the performance of the antiGen framework.

- **Mutation count matrix (MCM).** we first generate a matrix MCM whose (*i*, *j*)th entry is given by

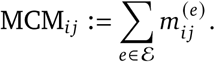 The matrix of predictions for an example *e* is given by

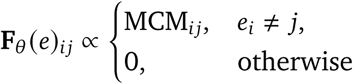 with the constant of proportionality determined such that the sum of entries of **F***_θ_*(*e*) equals 1.
- **Position frequency matrix (PFM).** We first generate a matrix PFM of dimension *L*×*A* whose (*i*, *j*)th entry PFM*_i j_* is the number of times site *i* exhibits allele *j* among the sequences used to construct the training tree, subject to the same filtering procedure as in **Phylogenetic tree construction and training data generation**. This matrix is converted to **F***_θ_*(*e*)*_i j_* via the same procedure as the MCM.
- **PFM conservation**. Building on the notation from the previous predictor, the matrix of next-mutation probabilities relative to a query *e* is given by

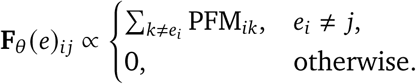 All non-query alleles at a site *i* receive the same score. Again, the normalization is such that the sum of all entries of **F***_θ_*(*e*) is 1.
- **Site** × **Substitution (S**×**S).** Consists of a vector *v* ∈ ℝ*^L^* that denotes how many mutations were observed at each site in the training data and a substitution matrix *S* ∈ ℝ*^A^*^×*A*^ where *S_i_*, *_j_* = *p*(mutation to *j* | wt = *i*) and is determined by counting how many times amino acid *i* mutated to amino acid *j* in the training data and row-normalizing. A mutation (ℓ, *j*) is then scored as *v*_ℓ_ · *S_i_*, *_j_* where *i* is the wildtype amino acid at site ℓ.
- **Codon-B62 Mixture.** A matrix *S̃* ∈ ℝ*^A^*^×*A*^ defined by

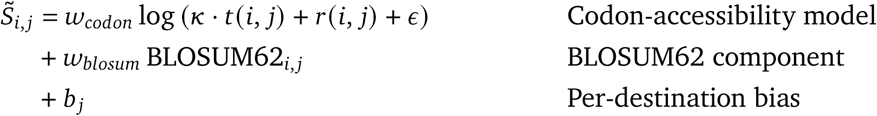 where *w_codon_*, *κ*, *w_blosum_*, and *b* ∈ ℝ*^A^* are parameters learned by gradient descent (Adam optimizer, learning rate 0.05, 300 steps, batch size 16). *t*(*i*, *j*) is the number of single-nucleotide transitions (*A* ↔ *G*, *C* ↔ *T*) from amino acid *i* that result in a codon for amino acid *j*, averaged over all possible synonymous codons for *i*. *r*(*i*, *j*) is the same statistic for transversions (*A* ↔ *C*, *A* ↔ *T*, *C* ↔ *G*, *G* ↔ *T*). We set *ε* = 0.01 to avoid log(0) values for amino acid pairs that always differ by at least 2 nucleotides. Analogous to the substitution matrix in the S×S model, *S̃* is row-normalized (using Softmax instead of linear normalization) and mutations are scored as *v*_ℓ_ · softmax*_j_* (*S̃*_*i*,_.)
- **Site** × **Mixture (S**×**M):** Consists of the same vector *v* as S×S and but replaces the substitution matrix with the Codon-B62 Mixture matrix described above.
- **S**×**S–MCM Interpolation.** A linear interpolation of the S×S and MCM (see **Zero-shot baselines**) models where the *S*×S component is scaled by a factor *η*. Intuitively, the S×S model can be thought of as providing pseudocounts to support the MCM model. This combined model also has a well-justified Bayesian interpretation as a Dirichlet-multinomial model: the normalized MCM is the MLE under a multinomial model of mutations, and the normalized S×S distribution can be thought of as the mean of a Dirichlet distribution over the (*L* × *A*) simplex with concentration *η*. Since a Dirichlet distribution is the conjugate prior for a multinomial distribution, the posterior mean is just a linear interpolation of the two models and *η* describes the strength of our belief in the Dirichlet prior. In practice, we normalize S×S to form a probability distribution over the sequence and set *η* equal to the total number of mutations in the training data, placing equal confidence in the likelihood and the prior and keeping the mixing weight consistent across viruses.
- **S**×**M–MCM Interpolation.** Same interpolation as described in the previous bullet but using S×M instead of S×S.

#### Existing computational methods

In addition to antiGen and statistical models, we also evaluated a survey of current methods without any modification specific to our prediction task on the SARS-CoV-2 data with January 1st, 2023 train-test split. These included ESM-2 without any fine-tuning, as well as the following:

- **EVE.** A variational autoencoder with a Bayesian decoder, trained on a multiple sequence alignment (Fraser et al., 2021). We used the published architecture (latent dimension 50; encoder 2000– 1000–300; decoder 300–1000–2000 with Bayesian weights) but trained it from scratch for each cutoff, on an alignment of SARS-CoV-2 spike sequences from that cutoff’s training window. These were the same sequences that make up antiGen’s training data. At the 2021-01-01 cutoff the alignment held all 351,410 sequences. At 2023-01-01 we drew 351,410 of its 7,813,585 sequences uniformly at random so that both cutoffs had the same training-set size; sequence weighting scales quadratically with alignment depth. Following the EVE conventions, we removed columns where the reference (Wuhan/IPBCAMS-WH-01/2019) has a gap, dropped sequences with more than 50% gaps, modeled only columns with fewer than 30% gaps, and reweighted sequences with *θ* = 0.01. At 2023-01-01 this excluded sites 69, 70, 156 and 157, all outside the RBD; mutations there were left unscored. We trained for 400,000 steps (batch size 256, learning rate 10^−4^) and then froze the model. Each query was encoded to its posterior mean, and a substitution to residue *a* at site *i* was scored by the decoder’s log-probability of *a* at *i*.
- **Tranception.** An autoregressive transformer whose attention heads include multi-scale depthwise convolutions and grouped ALiBi position encodings (Notin et al., 2022). As with EVE, we did not use the published UniRef-pretrained weights or inference-time retrieval. We trained the architecture from scratch (6 layers, embedding dimension 256, 8 heads, dropout 0.1) on the same per-cutoff alignments, filters and sequence weights, for 150,000 steps (batch size 32, peak learning rate 3 × 10^−4^), and then froze it. A substitution to residue *a* at site *i* of a query was scored by the mean of two conditional log-probabilities of *a* at *i*: one from the model read left to right, given the query’s preceding residues, and one read right to left, given its following residues.
- **CoVFit.** An ESM-2 model fine-tuned to predict SARS-CoV-2 spike variant fitness (Ito et al., 2025). We used the published `covfit_cli` model (fold 0), and evaluated it only in the fixed-split comparison at the 2024-10-01 cutoff. Unlike a static substitution map, CoVFit scores whole spike sequences, so we queried it on each test genotype’s own background: for each of the 921 distinct test-set genotypes with at least one subsequent mutation in the RBD, we scored the genotype itself and every RBD single-amino-acid mutant of it (201 positions × 19 alternative residues = 3,819 mutants). Each substitution’s score was the change in predicted fitness, i.e., fitness(mutant) − fitness(query), using the CLI’s mean fitness across its 17 countries. Sequences were passed in the 20-amino-acid alphabet with no deletion characters. Because the RBD (positions 331–531) lies within CoVFit’s 1,022-residue input window, every evaluated mutation was within its context.
- **PyR0**. A hierarchical Bayesian multinomial logistic regression that estimates the effect of each amino-acid substitution on lineage growth rate from PANGO lineage frequencies over time (Obermeyer et al., 2022). We used the per-mutation rankings released in the authors’ repository (https://github.com/broadinstitute/pyro-cov, `paper/mutations.tsv`, commit `f11fc53`, 2 February 2022), which were fit to the GenBank data available on that date. Each spike substitution was scored by its rank, with the most significant scoring highest, and unlisted substitutions scored zero. PyR0 is fit once and cannot be refit at an arbitrary date, so we evaluated it only at the 1 January 2022 cutoff. Its data extend somewhat past that cutoff, which favors PyR0.
- **Bloom and Neher (2023).** Per-site, per-amino-acid fitness effects for every SARS-CoV-2 protein, estimated by comparing the observed to the expected number of independent occurrences of each mutation along a phylogeny of public sequences (Bloom and Neher, 2023). We used their dated release built from sequences through 2023-10-01 and evaluated it only in the fixed-split comparison at that same cutoff. Each substitution was scored by its estimated fitness effect, the same for every query; substitutions whose expected count fell below 10, the authors’ threshold for reporting an estimate, were left unscored and ranked last.
- **MSA Transformer.** A protein language model that reads a multiple sequence alignment rather than a single sequence (Rao et al., 2021) (`esm_msa1b_t12_100M_UR50S`). We built an MSA of 256 SARS-CoV-2 spike sequences collected before 2020-12-23, with an early Wuhan isolate as the query, and scored every substitution by masked-marginal probability: each position of the query was masked in turn and the model’s distribution over amino acids at that position was recorded. The resulting map was used as a fixed predictor, the same for every query and every cutoff.
- **EVEscape.** EVEscape combines an EVE fitness term, trained on pre-pandemic coronavirus sequences, with structural accessibility and charge/hydrophobicity dissimilarity terms Thadani et al. (2023). We used the published receptor-binding domain (RBD) scores (https://github.com/OATML-Markslab/EVEscape, `results/summaries_with_scores/spike_rbd_evescape.csv`) for sites 331–531 without modification. Mutations outside the RBD are not scored.

#### Deep mutational scans

Deep mutational scans were treated as static predictors. For a query *e*, define DMS(*e*) ∈ (ℝ ∪ {−∞})*^L^*^×*A*^ by

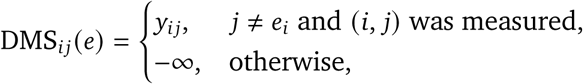

where *y_i j_* is the scan’s measurement for amino acid *j* at site *i*. We set **F***_θ_*_^_(*e*) = softmax(DMS(*e*)). Because softmax preserves order and our metrics depend only on ranks, this softmax step is not strictly necessary in practice. The deep mutational scans evaluated in this study were BA.1 binding and escape and BA.2 binding and escape (Starr et al., 2022). We intentionally omitted the BA.2 LY-CoV1404 escape DMS from the same paper due to 47% of RBD cells being unmeasured.

#### Zero-shot baselines

Finally, we evaluated a number of simple baselines that require no parameter fitting and are hence “zero-shot.” These include:

- **BLOSUM62**. a static *L*×*A* matrix of BLOSUM62 scores of each possible mutant allele relative to that of the query at each site. Specifically, the (*i*, *j*)th entry of this matrix for a query *e* is BLOSUM62*_ei_*_, *j*_ if *e_i_* ≠ *j*, otherwise −∞. The corresponding **F***_θ_*(*e*) would be obtained by taking the softmax over this matrix (though for the same reason as in the DMS predictor, this is not necessary in practice).
- **Codon-aware BLOSUM62.** Modified version of BLOSUM62 that accounts for nucleotide-level biases. Specifically, we take BLOSUM62, which is published as half-bit log odds *D* ∈ ℝ*^A^*^×*A*^ and Henikoff background frequencies *p* ∈ ℝ*^A^*, and convert it to an exchangeability matrix *Q* ∈ ℝ*^A^*^×*A*^ where

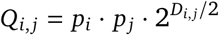

is proportion to the probability of amino acid *i* getting replaced by *j*, and *Q_i_*_,*i*_ = 0. Then we construct a codon accessibility matrix *C* ∈ ℝ*^A^*^×*A*^ by counting the number of single-nucleotide substitutions from amino acid *i* that result in a codon for *j*, averaged over all possible synonymous codons for *i*. That count weights transitions (*A* ↔ *G*, *C* ↔ *T*) by *κ* = 4. Then each row is normalized into a probability distribution and mixed with a uniform distribution over all *j* ≠ *i* with weight *ε* = 0.05 to ensure that two-nucleotide substitutions are still possible, albeit unlikely. The final model is given by row-normalizing *Q* ⊙ *C*, the element-wise product of the two matrices. Intuitively, this method combines the biochemical priors BLOSUM62 encodes about residue exchangeability with the nucleotide-level biases that dominate mutational patterns over the short timescales of viral evolution.

#### Multi-Tree Training and Evaluation

For multi-tree training, we finetune 3 versions of ESM-2 650M parameters using the antiGen objective and *partial* strategy defined in **Training and fine-tuning** and the same cross validation recipe as antiGen, except we set the maximum number of epochs to 30. The single-tree version is trained individually on each of the nine viral training sets. For the nine-tree and 49-tree versions, we randomly shuffle training examples from across viruses and select the best checkpoint based on validation negative log likelihood macro-averaged over the nine viruses being evaluated. Note that we evaluate the multi-tree models using the same checkpoint on all nine viruses rather than choosing a different checkpoint for each one.

Unlike the SARS-CoV-2 and endemic-virus evaluations, where a test example requires every leaf descending from the query (parent) node to be sampled on or after the train–test cutoff, here we retain a test example when every leaf descending from its *child* node was sampled on or after the cutoff. A query may therefore have descendant leaves sampled before the cutoff, provided none of them descends from the child that defines the test example. We make this relaxation to increase statistical power for viruses with sparse sampling while still avoiding data leakage, since the parent-child examples evaluated could not have existed in the training data alone.

#### Versioning

The original version of antiGen (0.1.0) was released on July 27th, 2026. For the current version (0.2.1), released November 13th, 2026, we primarily updated the training data generation pipeline, training regime, and evaluations, leaving the model architecture unchanged. In particular, the tree augmentation algorithm now maintains sets of possible parsimonious alleles at every node to improve parsimony scores (per **Phylogenetic tree construction and training data generation**), no longer requires GPU support, and allows for a choice between 20and 22-token alphabets (previously only 22 was supported). Moreover, the repository no longer splits a single augmented tree into training and testing examples, and rather enforces two separate runs of the augmentation algorithm—one using a historical tree dated before the desired train-test split date and the other using a present tree. In terms of model training, validation loss can now be used to select the final training epoch and cross-validation is supported (previously, all models were trained for a fixed 32 epochs). Model evaluation code has been adapted to the new testing data format and new train-test split dates, and various minor bug fixes have been applied to the quantile calculations.

The current manuscript reflects antiGen version 0.2.1 with the exception of the *in vitro* pseudoviral infectivity assay, which is based on predictions from version 0.1.0.

#### Cell culture

HEK293T/ACE2/TMPRSS2 cells were obtained from Peter S. Kim’s laboratory at Stanford University. Both HEK293T and HEK293T/ACE2/TMPRSS2 cells were grown in a 5% CO_2_ humidified incubator at 37^◦^C in complete D10 medium consisting of Dulbecco’s Modified Eagle Medium (DMEM; Corning) supplemented with 10% fetal bovine serum (ThermoFisher), 2 mM L-glutamine, and 1% penicillin/streptomycin (Gibco).

#### Lentiviral plasmids

An HDM-SARS2-Spike expression plasmid encoding the SARS-CoV-2 Wuhan-Hu-1 spike protein with a 21-amino-acid C-terminal truncation (Δ21) in the HDM expression backbone (“HDM-SARS2-Spikedelta21”; Addgene plasmid #155130) and the lentiviral packaging plasmids pHAGE-Luc2-IRES-ZsGreen (NR-52516), HDM-Hgpm2 (NR-52517), pRC-CMV-Rev1b (NR-52519), and HDM-Tat1b (NR-52518) were obtained from BEI Resources (NIAID, NIH; SARS-Related Coronavirus 2, Wuhan-Hu-1 Spike-Pseudotyped Lentiviral Kit V2, NR-53816). The Δ21 truncation enhances pseudovirus production and was applied uniformly across all spike variants (Crawford et al., 2020; Weidenbacher et al., 2022). Sequences of all plasmids used for pseudovirus production are provided in **Data S1**.

#### Spike plasmid design and construction

The sequence of the Wuhan-Hu-1 spike protein within the HDM-SARS2-Spike-delta21 parental construct was replaced with the corresponding XFG spike sequence (GenBank accession YCJ32983.1; synthesized by Twist Bioscience) by PCR-based cloning. All antiGen-predicted mutations in XFG spike reported throughout this study are numbered relative to this wild-type XFG spike reference sequence, annotated accordingly on the plasmid map “HDM_SARS2_Spike_del21_XFG_antiGen” in **Data S1**. Individual amino acid substitutions were introduced into the XFG spike construct by site-directed mutagenesis using mutation-specific primer pairs, as described in **Data S1**. All primers were synthesized by Integrated DNA Technologies.

All cloning reactions were performed using NEBNext High-Fidelity Polymerase (New England Biolabs, Cat. No. M0544) for PCR amplification, followed by In-Fusion assembly (Takara Bio; #638948) or Gibson assembly (New England Biolabs; #E2621S) of fragments according to the manufacturers’ protocols. Recombinant plasmids were transformed into Stellar Competent Cells (Takara Bio; #636766) for propagation and plasmid production. Plasmid DNA was purified using either the QIAprep Spin Miniprep Kit (Qiagen) or the ZymoPURE II Plasmid Midiprep Kit (Zymo Research), according to the manufacturers’ instructions. The sequences of all constructs were confirmed by whole-plasmid Nanopore sequencing.

#### Antibody production

Purified anti-SARS-CoV-2 RBD antibody BD55-1205 was obtained from GenScript. Purified anti-SARS-CoV-2 spike antibody CV10 was provided by the laboratory of Peter S. Kim, Stanford University. The heavyand light-chain variable region amino acid sequences of all antibodies used in this study are provided in **Data S2**.

#### Production of SARS-CoV-2 pseudovirus

SARS-CoV-2 pseudoviruses were generated using a five-plasmid lentiviral system in HEK293T cells. Cells were seeded at 6 × 10^6^ cells per 10-cm dish one day prior to transfection.

Plasmids were combined in D10 medium at the following amounts: 10 *μ*g pHAGE-Luc2-IRES-ZsGreen, 3.4 *μ*g spike expression plasmid, 2.2 *μ*g HDM-Hgpm2, 2.2 *μ*g HDM-Tat1b, and 2.2 *μ*g pRC-CMV-Rev1b. BioT transfection reagent (30 *μ*L; BioLand #B01-01) was added, and complexes were formed for 10 min at room temperature before addition to cells (Powell et al., 2026).

After 24 h, media was replaced with fresh D10 medium. Viral supernatants were harvested at 72 h post-transfection, centrifuged at 300 ×*g* for 5 min, and filtered through a 0.45 *μ*m PES filter. Viral stocks were aliquoted and stored at −80^◦^C.

For concentration, filtered supernatants were processed using Lenti-X Concentrator (Takara Bio #631231) at a 3:1 ratio (1.8 mL supernatant + 0.6 mL reagent), incubated overnight at 4^◦^C, and centrifuged at 1,500 ×*g* for 45 min. Pellets were resuspended in DMEM at one-tenth of the original volume, yielding approximately 10× concentrated stocks.

#### Flow virometry

##### Flow virometry gating strategy

Pseudovirus-containing supernatants were diluted in 0.02 *μ*m-filtered Dulbecco’s Phosphate-Buffered Saline (DPBS; Gibco) prior to acquisition. Flow virometry was performed using a CytoFLEX Nano flow cytometer (Beckman Coulter) at a flow rate of 1 *μ*L/min for 30 s. Sample dilutions were adjusted as necessary to maintain the manufacturer’s recommended event rate. For all flow virometry analyses, events below the SSC-H threshold were excluded as instrument noise, after which singlet events were identified using SSC-W versus SSC-H gating (**Figure S7A**; Jungbauer-Groznica et al. (2026)). To establish a size-based pseudovirus gate, nanoViS low nanoscale sizing standards (Beckman Coulter #D03231) were first used to define bead-equivalent SSC boundaries (**Figure S7B**). Based on reported lentiviral particle diameters (80–100 nm; Transfiguracion et al. (2020)) and spike glycoprotein protrusion (about 20 nm; Tai et al. (2021)), the expected pseudovirus population was estimated to fall within an 80–140 nm size-equivalent range (**Figure S7B**).

To distinguish pseudovirus-associated events from background particles, three control preparations were analyzed: (i) A mock-transfected conditioned medium control consisting of culture supernatant collected from producer HEK293T cells without plasmid transfection was used to characterize extracellular vesicles and other cell-derived background particles; (ii) A ΔGag/Pol control, generated in the absence of the Gag/Pol packaging plasmids, served as a negative control for pseudovirus particle assembly; and (iii) A ΔSpike pseudovirus control, generated by omitting the spike-expression plasmid, served as a spike-negative pseudovirus reference.

For crude pseudovirus preparations, SSC distributions showed substantial overlap between wild-type (WT) and mock-transfected samples. However, following 10× concentration, the WT preparations exhibited a readily distinguishable particle population within the expected SSC range relative to the mock-transfected control, enabling confident definition of the SSC gate (**Figure S7C**). SSC-based particle gates were therefore determined using 10× concentrated pseudovirus preparations.

The final pseudovirus particle gate was defined on singlet events as SSC-H = 4 × 10^4^–4 × 10^5^, corresponding to the bead-calibrated size region enriched for WT pseudovirus events while excluding the majority of low-scatter events observed in mock-transfected conditioned medium. This SSC gate was used for all subsequent flow virometry analyses. The flow virometry results were analyzed using FlowJo v10.10.1 and v11.1.1 Software (BD Life Sciences).

##### Particle quantification and linearity assessment

Particle abundance was quantified using CytoFLEX Nano as the number of SSC-gated events acquired at a fixed flow rate of 1 *μ*L/min over a 30 s acquisition window, and converted to concentration based on the known acquisition volume, sample dilution factors, and sample concentration steps applied prior to acquisition.

To evaluate the quantitative operating range of CytoFLEX Nano particle enumeration, WT pseudovirus preparations were analyzed across a 1×–10× concentration series. Measured particle counts were compared with the expected proportional scaling, and linearity was assessed using linear regression. SSC-gated particle counts exhibited linear scaling across the tested concentration range (**Figure S7D,E**). Accordingly, 10× concentrated pseudovirus preparations were diluted 4-fold prior to acquisition, which maintained the event rate within the manufacturer’s recommended operating range while preserving measurements within the validated linear range.

##### Fluorescence-based validation of SSC-gated pseudovirus particles

To characterize spike-bearing particles within the SSC-defined nanoparticle population, 10× concentrated pseudovirus preparations were stained with monoclonal antibodies against the SARS-CoV-2 spike protein (Jungbauer-Groznica et al., 2026; Burnie et al., 2026). Because all antiGen-predicted mutations were located within the receptor-binding domain (RBD), the conserved S2-specific monoclonal antibody CV10 (human IgG1; Weidenbacher et al. (2022)) was used to detect spike-bearing particles while minimizing the potential effects of RBD mutations on antibody recognition. In parallel, the anti-RBD monoclonal antibody BD55-1205 (human IgG1; Jian et al. (2025)), a broadly neutralizing antibody recognizing a receptor-binding epitope distinct from all antiGen-predicted mutation sites analyzed in this study, was used to detect the display of intact RBD on pseudovirus preparations. 10-fold concentrated pseudovirus preparations were incubated with 1 *μ*g/mL CV10 or BD55-1205, as optimized by titration on WT or ΔSpike pseudovirus preparations (**Figure S15A, B**), followed by Alexa Fluor 647 (AF647)-conjugated goat anti-human IgG secondary antibody (Invitrogen #A-21445; 1:2,000 dilution). Following singlet gating, spike staining was analyzed using SSC-H versus AF647-H plots to assess the distribution of AF647-positive events across the SSC range (**Figure S7A**). AF647-positive and AF647-negative populations were identified using a fixed AF647-H fluorescence gate established from the ΔSpike pseudovirus, secondary antibody-only, and isotype control samples (**Figure S15C**). Stain performance was quantified using the stain index described in Maltseva and Langlois (2022) (**Figure S15D**):

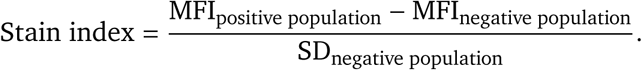

To verify single-particle detection, stained pseudovirus preparations were serially diluted prior to acquisition. Particle concentration decreased linearly with dilution, whereas the median fluorescence intensity (MFI) of the antibody-positive population remained constant, indicating that fluorescence measurements reflected individual particles rather than coincident events (**Figure S15E**).

Although antibody staining confirmed that the SSC-defined population was enriched for spike-bearing particles across all variants (**Figure S16**, **Figure S17**), fluorescence measurements were used to validate and characterize the particle population rather than for particle normalization. Antibody positivity reflects not only the presence of spike but also antibody accessibility, epitope recognition, and the amount and conformation of spike displayed on individual particles, all of which influence antibody binding. Consequently, antibody-based measurements may underestimate the number of infection-competent particles if functional spike-bearing particles exhibit reduced antibody-binding because of differences in epitope accessibility, antibody recognition, or spike presentation. Therefore, antibody-independent SSC-based particle counts (**Figure S8**) were used for particle normalization after establishing a common linear operating range across all variants.

#### Particle-normalized infectivity assay

To compare infectivity across viral variants independently of differences in pseudovirus production, infectivity was measured as relative luminescence units per SSC-defined physical particle (RLU/particle).

A pilot infectivity study was performed using WT and all mutant pseudoviruses to define the linear operating range of the luciferase assay for each variant. Crude pseudovirus preparations were subjected to a 10-point, 3-fold serial dilution, and 100 *μL* of each dilution was added per well to HEK293T/ACE2/ TMPRSS2 cells. Following infection, cells were incubated for 48 h at 37^◦^C. Luciferase activity was then measured by adding 100*μ*L of BriteLite reagent (prepared as a 1:1 mixture with DPBS) to each well, and luminescence was quantified as relative luminescence units (RLU) using a Tecan Spark Multimode Microplate Reader.

The dilution-response relationship was evaluated to identify the range over which RLU scaled proportionally with viral input. Particle input per well was calculated from the flow virometry-measured particle concentration (**Figure S8**) and the corresponding inoculation volume for each dilution (**Figure S9A**). A construct-specific linear operating range was then determined for each pseudovirus variant. For the wild-type XFG pseudovirus and all variants except 475insC, the first four dilution points (1:1 to 1:27) exhibited an approximately linear relationship between particle input and RLU. In contrast, 475insC generated quantifiable luminescence only at the first two dilution points and was therefore excluded from defining the common linear operating range used for subsequent assay optimization (**Figure S6**).

A common particle concentration range was subsequently selected for a particle-normalized infectivity assay such that the second and third points of a four-point, 3-fold serial dilution series fell within this shared operating window. When the target starting particle concentration exceeded that of the unconcentrated pseudovirus supernatant, 10× concentrated pseudovirus prepared in DMEM was used to achieve the desired starting concentration. All samples had a final FBS concentration of 10% throughout the dilution series. Following infection, cells were incubated for 48 h at 37^◦^C, and luciferase activity was measured as described in **Figure S9B** and **Figure S10**.

Infectivity was then calculated as:

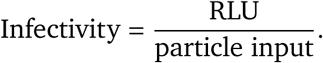

Because the second and third dilution points were predetermined to fall within the shared linear operating window across all constructs, these dilution points were designated *a priori* as the primary comparison range. Infectivity values from the two dilution points were averaged to generate a single infectivity estimate for each condition.

Relative infectivity was calculated by normalization to WT XFG measured in parallel within each experiment:

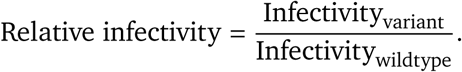

Final values represent the mean across independent experiments.

#### Ethics declaration and institutional review

Work with SARS-CoV-2 spike-pseudotyped lentiviral particles was approved by the Stanford University Institutional Biosafety Committee/Administrative Panel on Biosafety under APB protocol 6007-BH0126. The protocol covers production and use of lentiviral particles pseudotyped with SARS-CoV-2 spike proteins from variant spike proteins, including emerging variants of concern, and spike constructs containing point mutations within the receptor-binding domain. All pseudotyped lentiviral particles were replication-incompetent, used for single-cycle entry infectivity assays, and handled under approved institutional biosafety procedures at BSL-2 containment.

#### Statement on AI use

Claude Opus 4.6–4.8, 5, and 5.5 were used to assist in several areas of the codebase, including parsing input data files, evaluating existing models, templating the Transformer training loop, revising figure plotting code, and packaging antiGen as a Python library. While all code was edited and reviewed by AI tools, original implementations of the phylogenetic tree augmentation logic and the training loss function were written by the authors. All authors take full responsibility for the entire content of the manuscript, including any portions developed with the assistance of AI tools, and have reviewed and verified all text, code, figures, and results.

## Data Availability

Training data were assembled from public phylogenetic trees and sequences available via Nextstrain (Hadfield et al., 2018), LAPIS (Chen et al., 2023), and GISAID (Shu and McCauley, 2017; Wallau et al., 2023) as detailed in **Methods**.

Sequences of the RBD mutants generated for pseudovirus variant production in this study, primer sequences used for cloning, and template vector information used for pseudovirus preparation are provided as **Data S1**. Amino acid sequences of the antibody heavyand light-chain variable regions used in this study are provided as **Data S2**. Both data files are available to the public at https://doi.org/10.5281/zenodo.21481224.

## Code Availability

Model training, querying, and evaluation code, as well as the training data generation pipeline, are available to the public at https://github.com/evo-design/antiGen.antiGen is available as a Python package, installable from PyPI with `pip` `install` `antigen-model`.

## Acknowledgements

We thank Daniel Chang, Marc Lipsitch, Aditi T. Merchant, Pardis C. Sabeti, Talal Widatalla, and members of the Laboratory of Evolutionary Design for helpful discussions and assistance with manuscript preparation. We thank members of the Felix Horns lab for assistance with flow virometry. We thank members of the Peter S. Kim lab for contributing experimental reagents related to the pseudoviral infectivity assays.

I.S. acknowledges funding support from the Fannie and John Hertz Foundation and Knight-Hennessy Scholars. S.P. acknowledges funding support from the Stanford Graduate Fellowship and Sarafan ChEM-H Chemistry/Biology Interface Program. S.C. acknowledges funding support from the Masason Foundation fellowship. G.B. acknowledges funding support from the National Science Foundation Graduate Research Fellowship Program. J.A.P. acknowledges funding support from NIH R35 GM148338 and NSF career award 2143242. B.L.H. acknowledges funding support from the Gates Foundation, the Silicon Valley Community Foundation, the NIH Director’s New Innovator Award, the Advanced Research Projects Agency for Health (ARPA-H), Arc Institute, Schmidt Sciences AI2050, Stanford Center for Digital Health, and Stanford Human-Centered Artificial Intelligence (HAI) Hoffman-Yee Research Grants.

## Author Contributions

I.S., G.B., and B.L.H. conceived the study. B.L.H. supervised the study. I.S. curated the training data and performed phylogenetic tree construction. I.S., G.B., and J.A.P. developed the phylogeny-informed training objective. I.S., G.B., and A.V. conducted model training and fine-tuning. I.S. and S.C. conducted the development and evaluation of the gated models. I.S. and A.V. conducted the model evaluations. S.P. conducted pseudovirus production, flow virometry, infectivity assays, experimental optimization, quality control, and data analysis. C.L.D. provided assistance with and feedback on the experimental methodology. S.P. and C.L.D. performed molecular cloning. I.S., S.P., S.C., C.L.D., and B.L.H. wrote the initial draft of the manuscript. All authors wrote the final version of the manuscript.

## Competing interests

B.L.H. acknowledges outside interest in Arpelos Biosciences and Genyro, Inc. as a scientific co-founder. All other authors declare no competing interests.

## Supplementary Figures

**Figure S1.**
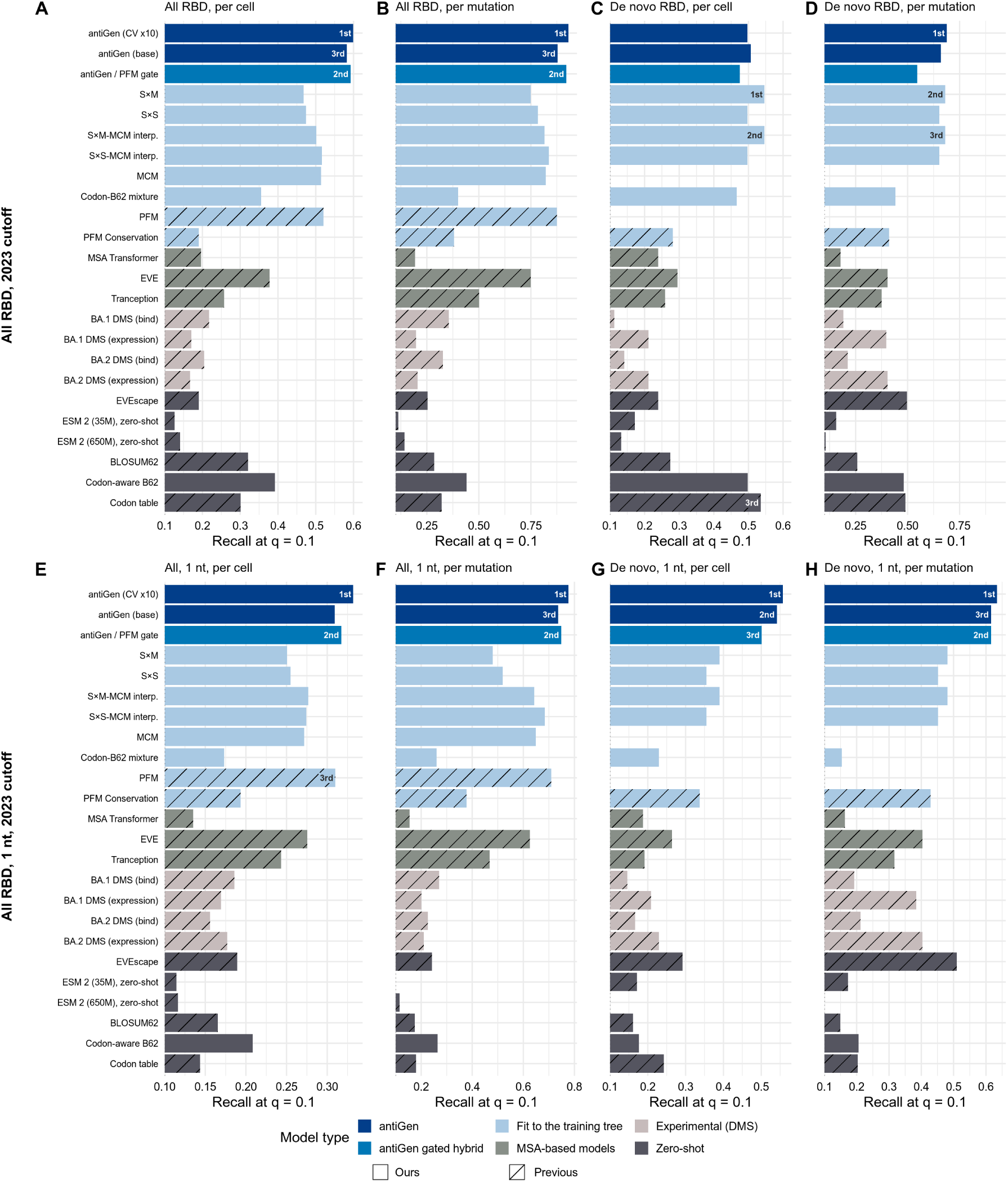
SARS-CoV-2 forecasting for a wider panel of models. Top-decile recall on SARS-CoV-2 spike mutations in the RBD with a train-test cutoff of January 1st, 2023, parallel to Figure 2C**–J**, expanded to a wider panel of models including the antiGen-PFM gate, antiGen trained at a fixed 32 epochs without cross-validation (“base”), and additional DMSs and statistical models.

**Figure S2.**
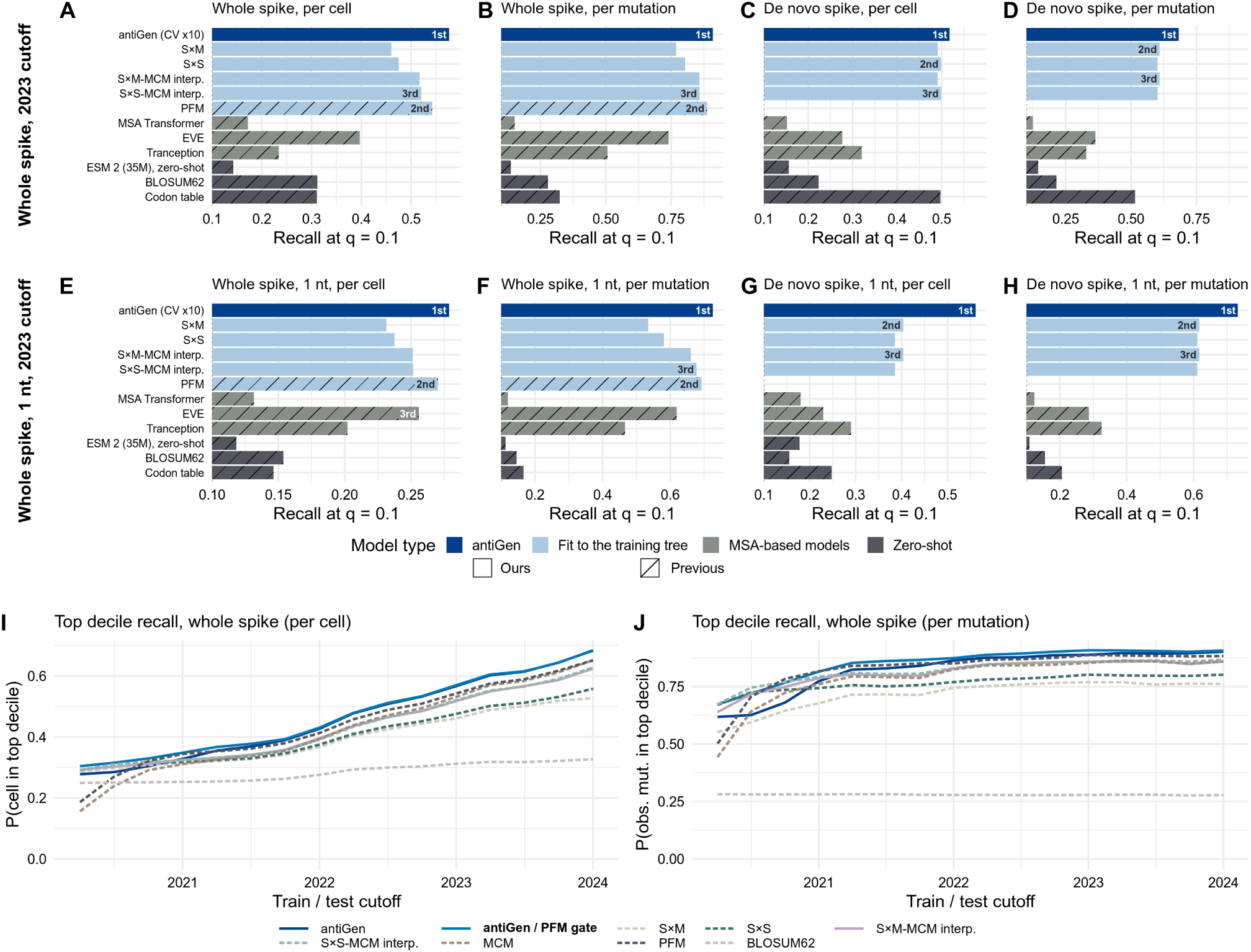
Forecasting evolution of the full SARS-CoV-2 spike protein. Top-decile recall on SARS-CoV-2 mutations with a train-test cutoff of January 1st, 2023, parallel to Figure 2C**–J****, O–P**, expanded to evaluation on the whole spike protein instead of the RBD alone.

**Figure S3.**
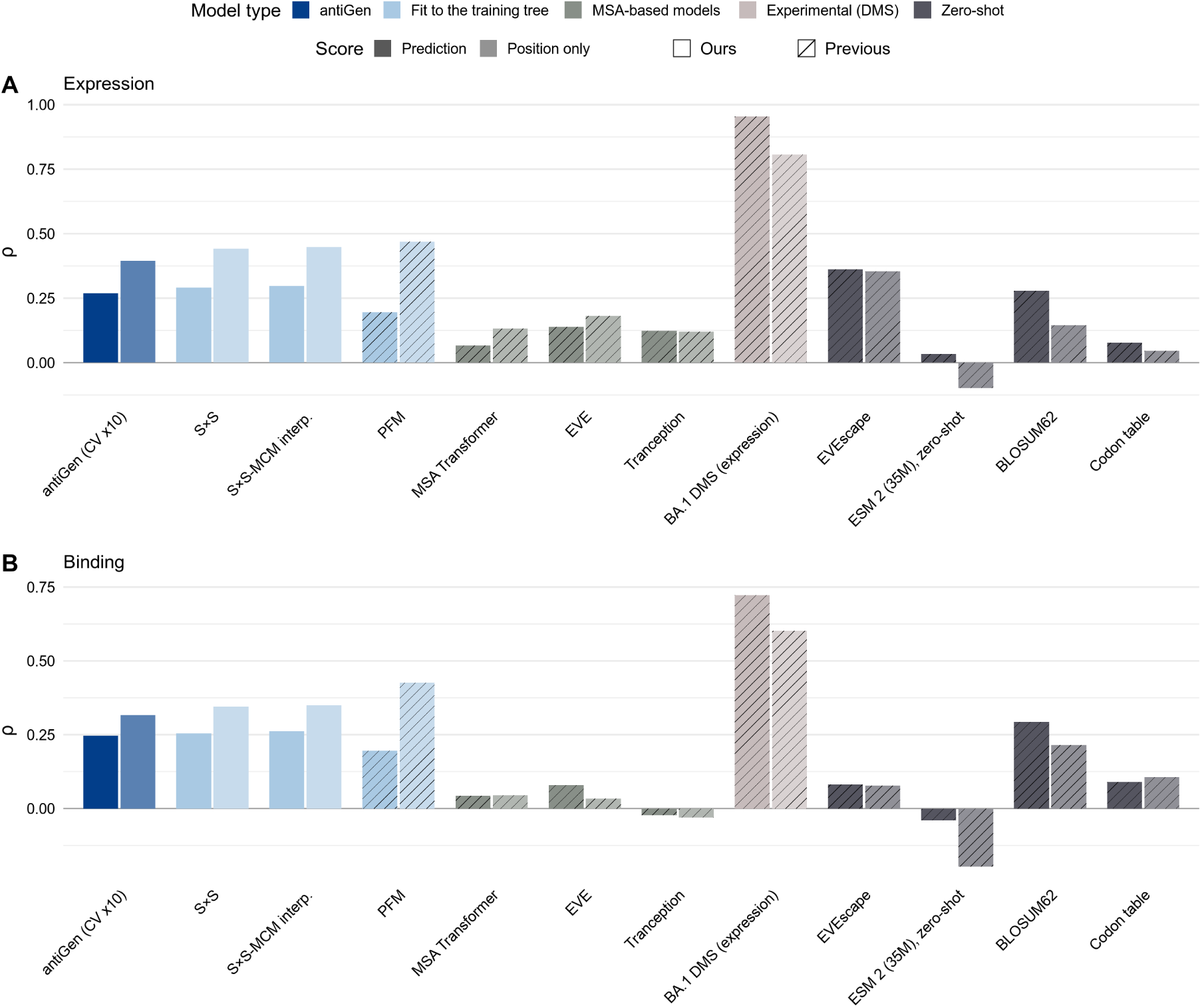
SARS-CoV-2 spike DMS correlation. Evaluation of antiGen and other models in terms of the Spearman correlation of their RBD mutation predictions with a BA.2 **(A)** expression and **(B)** binding DMS (Starr et al., 2022). Spearman correlations shown as-is (darker bars) and after replacing the rank of all possible amino acid changes at a given position with the average rank at that position, removing a model’s ability to discriminate between different possible substitutions at the same position (paired lighter bars).

**Figure S4.**
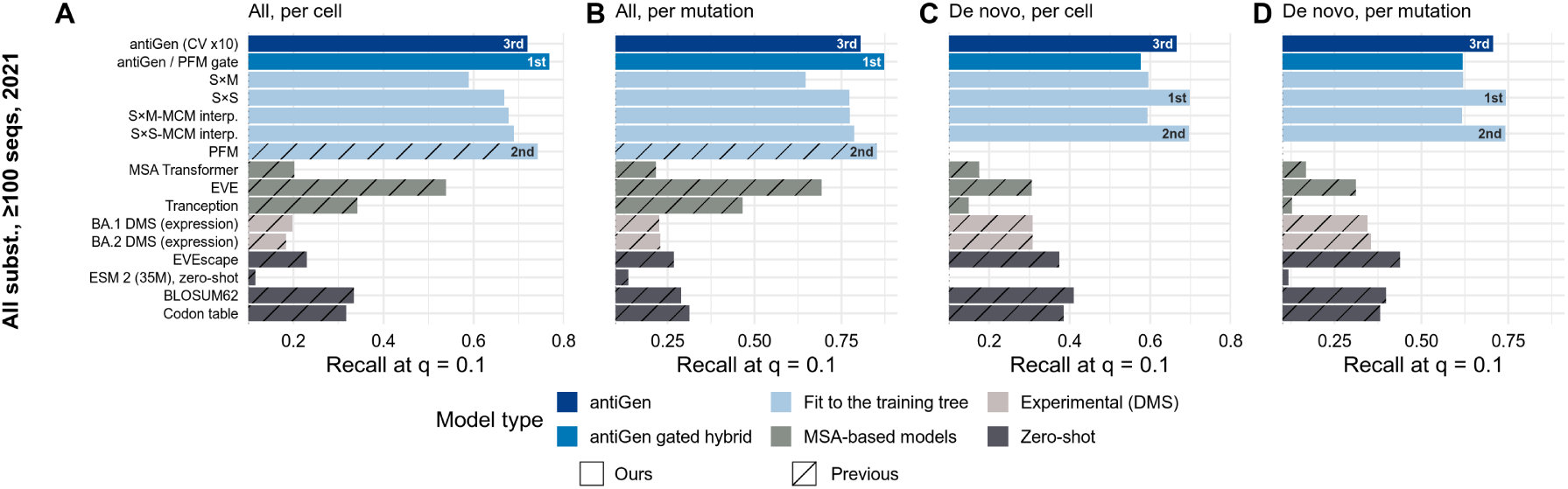
Forecasting SARS-CoV-2 spike RBD substitutions above a prevalence threshold. Top-decile recall on SARS-CoV-2 spike mutations in the RBD observed at frequency ≥ 100 after the train-test cutoff date of January 1st, 2021, parallel to Figure 2K**–N** except without the *de novo* restriction and including the antiGen-PFM gate.

**Table S1.**
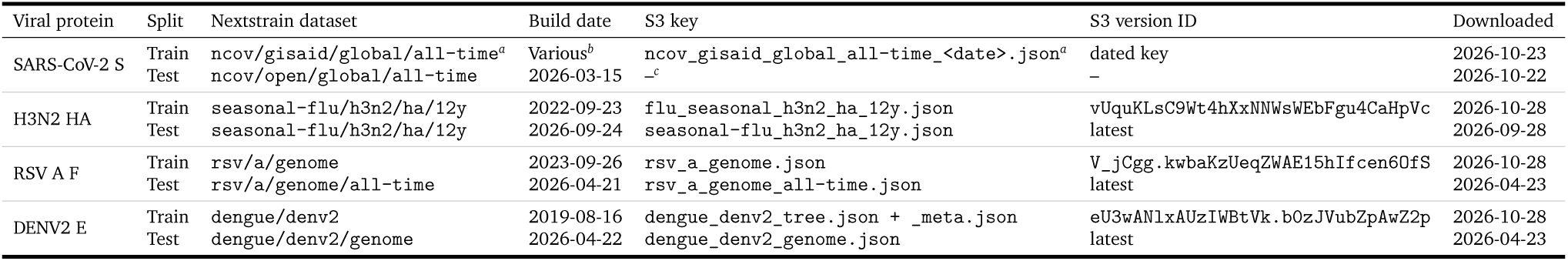
Nextstrain trees for SARS-CoV-2 and endemic viruses. Training trees are the latest build published before each cutoff, retrieved from Nextstrain’s public S3 bucket (`nextstrain-data`). For the endemic viruses (H3N2 2022-10-01, RSV A 2023-10-01, DENV2 2019-10-01), these are versions of a single undated object. SARS-CoV-2 builds are written under dated keys, one per build. Test trees for the endemic viruses are the current build at download. *^a^*The build has been renamed over time: `ncov`, `ncov/global` and `ncov/gisaid/global` before `ncov/gisaid/global/all-time`, with S3 keys `ncov_<date>`, `ncov_global_<date>`, `ncov_gisaid_global_<date>` and `ncov_gisaid_global_all-time_<date>`. Root-sequence sidecars without a dated key were taken as the latest object version at or before the build date. *^b^*One build per quarterly cutoff from 2020-04-01 to 2026-01-01, built 2020-03-31 to 2025-10-10. *^c^*Retrieved through Nextstrain’s API (getDataset?prefix=/ncov/open/global/all-time@2026-03-21), which returns the latest build at or before the requested date; the build is dated by its own `meta.updated`.

**Figure S5.**
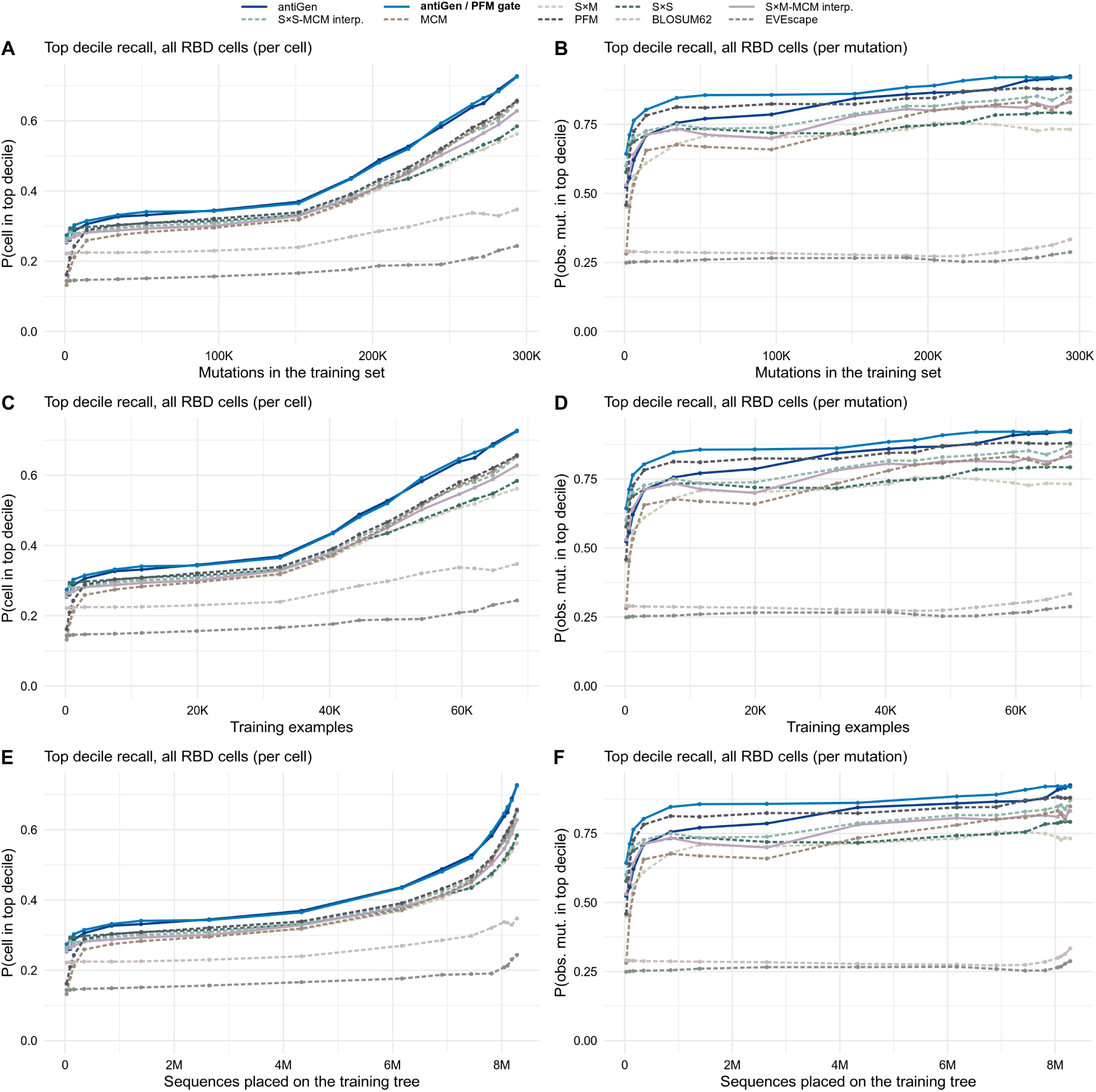
Model performance by training data statistics. The evaluation of Figure 2O**, P**, except with the *x* axis in units of **(A, B)** total number of mutations in the training set, **(C, D)** number of training examples, and **(E, F)** number of sequences used in the generation of the training examples.

**Figure S6.**
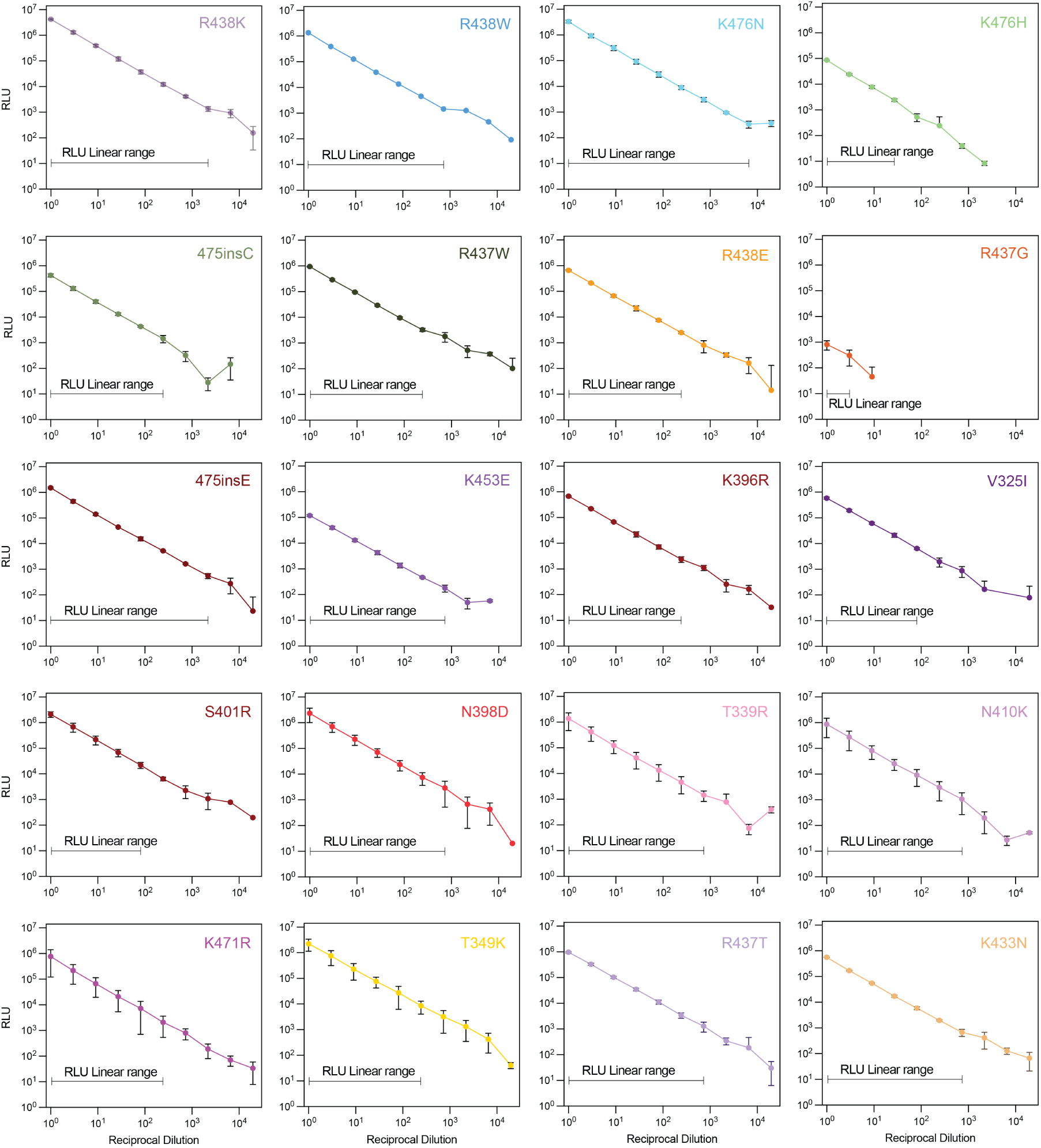
Serial-dilution infectivity profiling of antiGen-predicted SARS-CoV-2 XFG spike variants. 10-point serial-dilution infectivity assays were performed for lentiviral pseudoviruses bearing XFG spike variants containing antiGen-predicted mutations. Luciferase reporter signal is plotted as RLU against reciprocal dilution for each pseudovirus stock. The dilution points over which reporter signal scaled proportionally with viral input are indicated on each plot. Points show mean values and error bars indicate standard deviation (n = 3).

**Figure S7.**
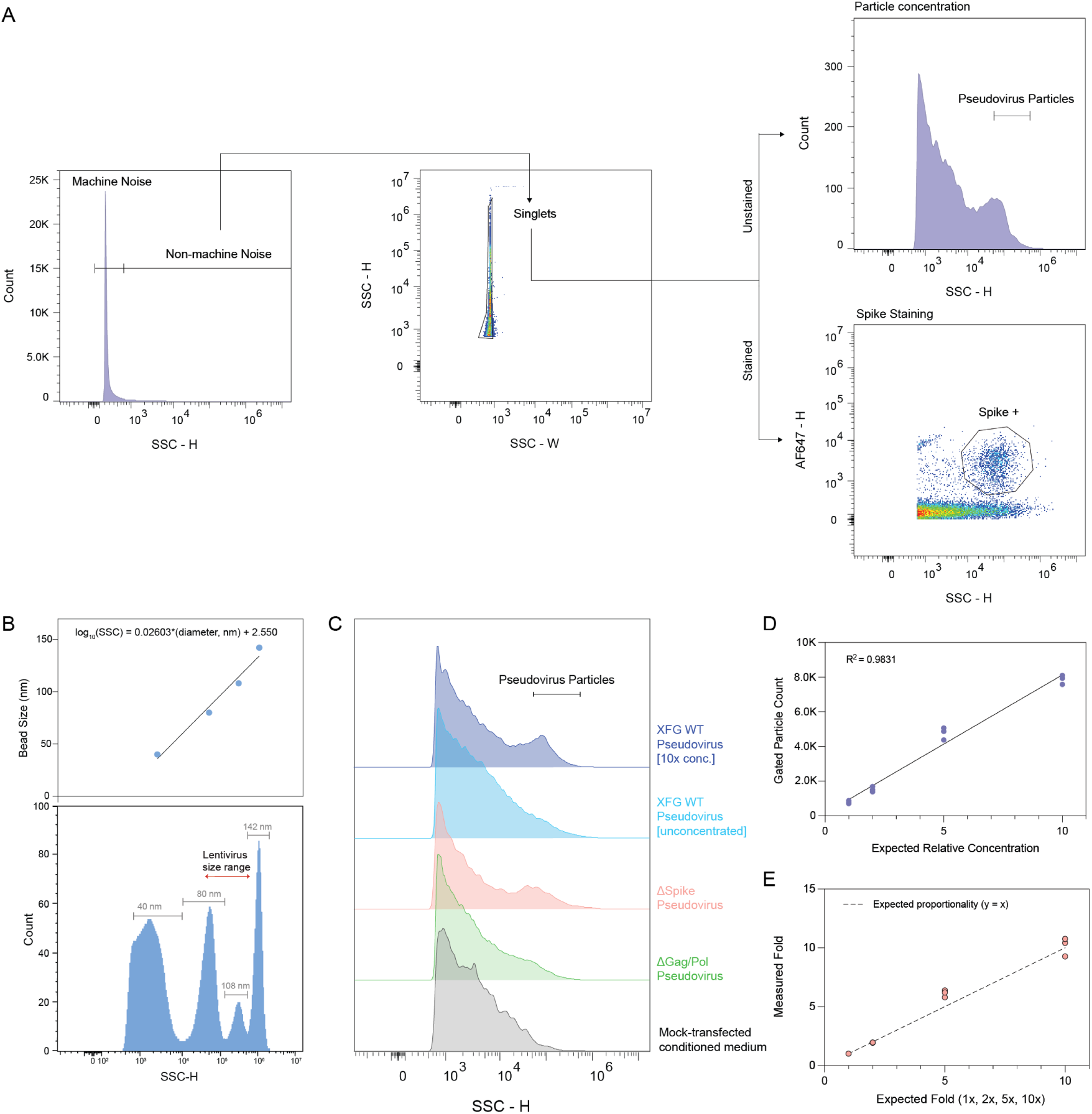
Establishment and validation of pseudovirus particle gating by flow virometry. **(A)** Sequential CytoFLEX Nano gating strategy for pseudovirus particle identification. Events were separated from machine noise by SSC-H, singlets were selected using SSC-H and SSC-W, and pseudovirus-associated particles were gated by SSC-H; spike-positive events were identified by AF647 fluorescence after spike staining. **(B)** Calibration of SSC-H signal using size-standard beads. The relationship between bead diameter and SSC-H was used to define the approximate lentivirus size range for analysis. **(C)** Representative SSC-H distributions of mock-conditioned medium and pseudovirus preparations. WT pseudovirus after 10× concentration or pseudovirus lacking spike showed enrichment of a distinct SSC-H population relative to mock and pseudovirus production controls lacking Gag/Pol. **(D)** SSC-gated particle counts increased linearly with expected relative concentration in a WT pseudovirus dilution series (*R*^2^ = 0.9831). **(E)** Measured fold changes approximately followed expected dilution factors, with underestimation at the highest concentration. These data validate the linear operating range of SSC-based particle enumeration used for subsequent particle quantification (n = 3).

**Figure S8.**
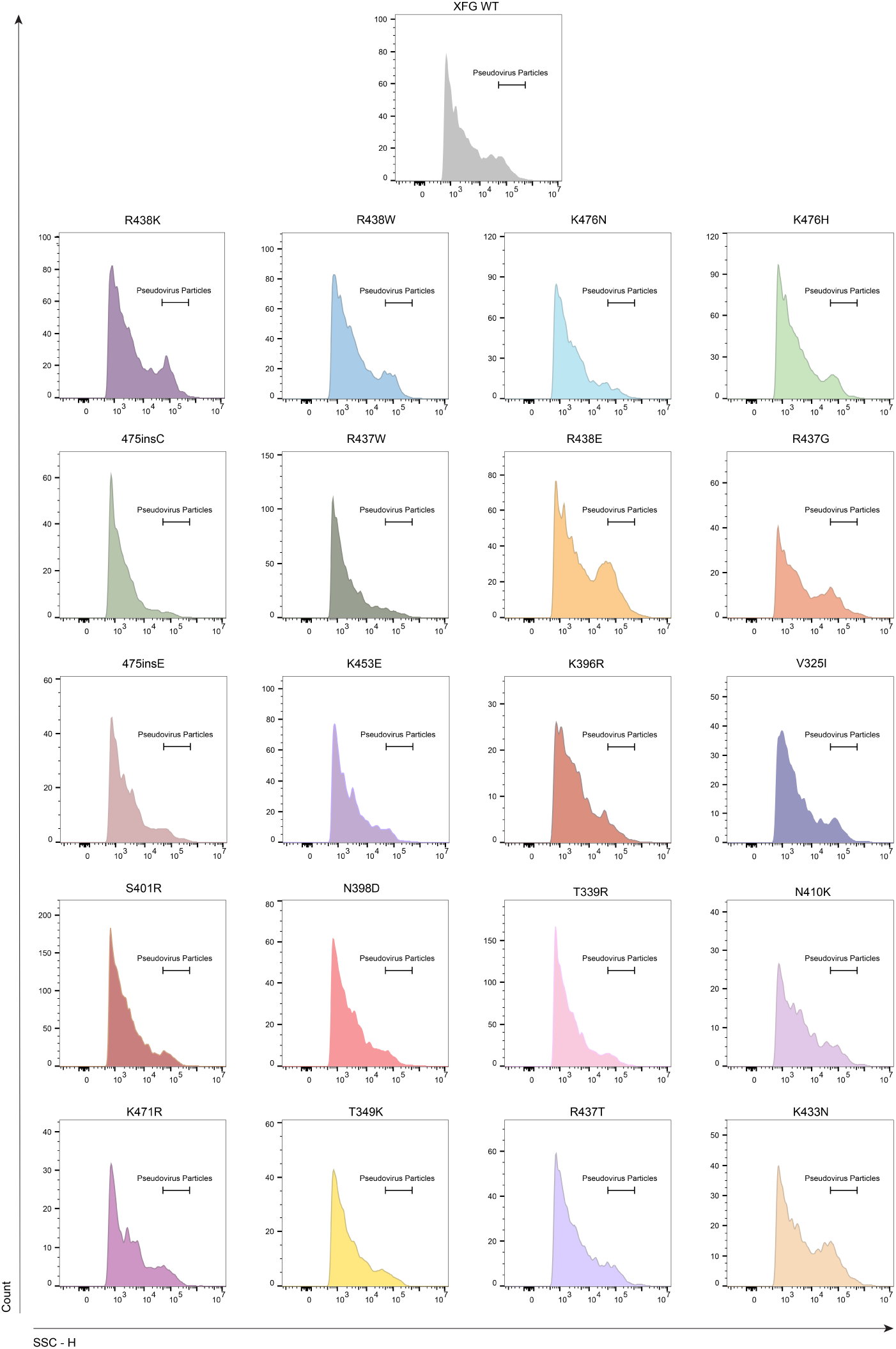
SSC-based particle gating for quantification of XFG spike-pseudotyped lentiviral preparations. Representative SSC-H histograms of XFG WT and antiGen-predicted XFG spike variant pseudovirus preparations analyzed by flow virometry. The indicated SSC-H gate was applied consistently across all preparations to quantify pseudovirus particle count for downstream particle normalization. Histograms show the distribution of events after exclusion of machine noise and singlet gating, with particle counts derived from events falling within the defined pseudovirus particle gate. Representative histograms from one of two replicate acquisitions are shown, and stock particle concentrations were calculated as the mean of the two measurements (*n* = 2).

**Figure S9.**
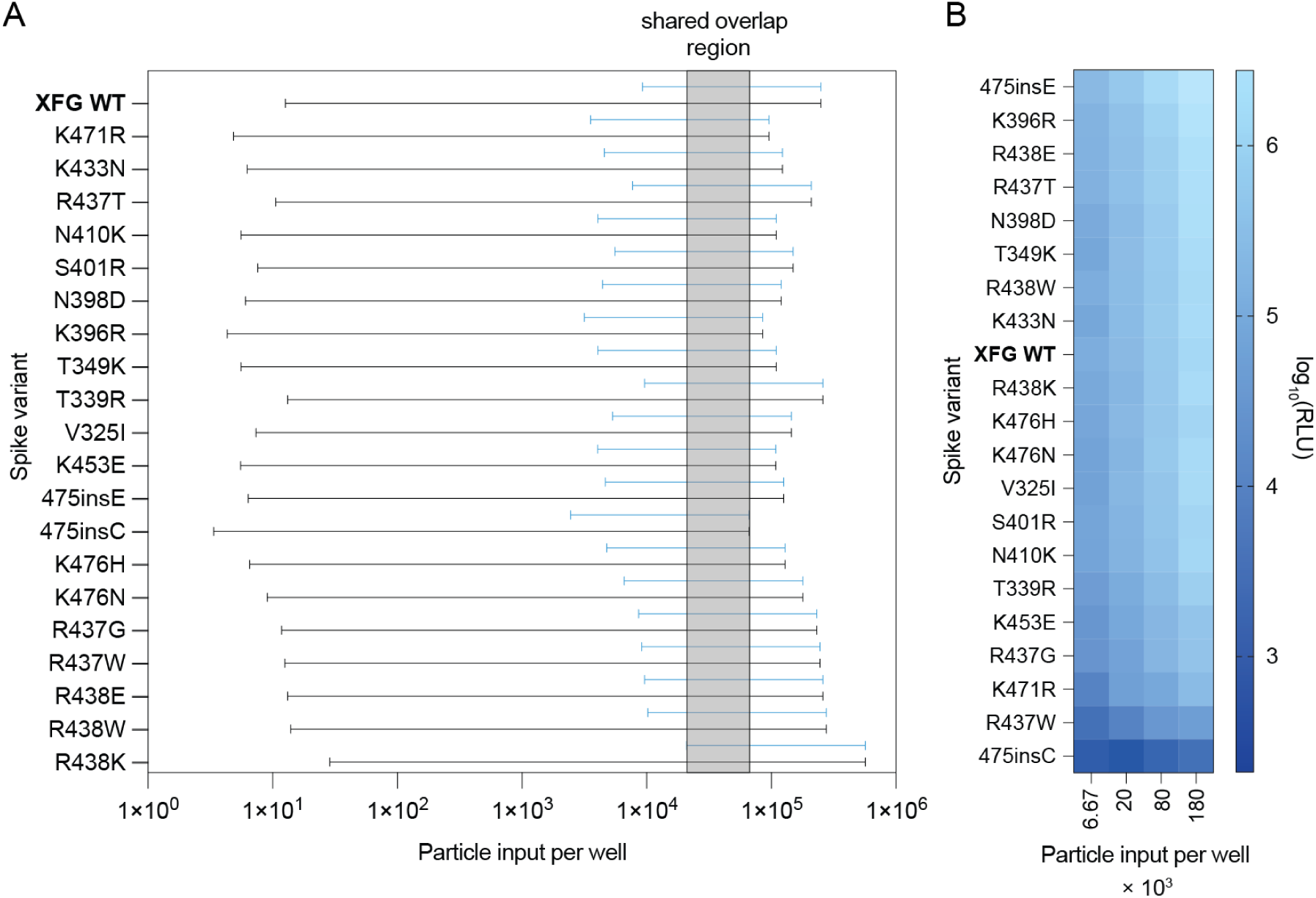
Defining a shared particle-input window for particle-normalized infectivity measurements. **(A)** Construct-specific infectivity assay dilution ranges were converted into particle input per well using flow virometry-derived particle concentrations and the infection volume. Black horizontal lines indicate the full particle-input range sampled in the initial 10-point serial-dilution infectivity assay for each spike variant. The shaded region denotes the overlap of the particle-input ranges over which each pseudovirus variant exhibited an approximately linear relationship between particle input and RLU. Blue horizontal lines indicate the subset of the initial 10-point dilution series that fell within this common linear window; the corresponding shared particle-input range was subsequently used to design the four-point particle-normalized infectivity assay. **(B)** Heat map of luciferase reporter signal from the four-point particle-normalized infectivity assay across predetermined particle inputs.

**Figure S10.**
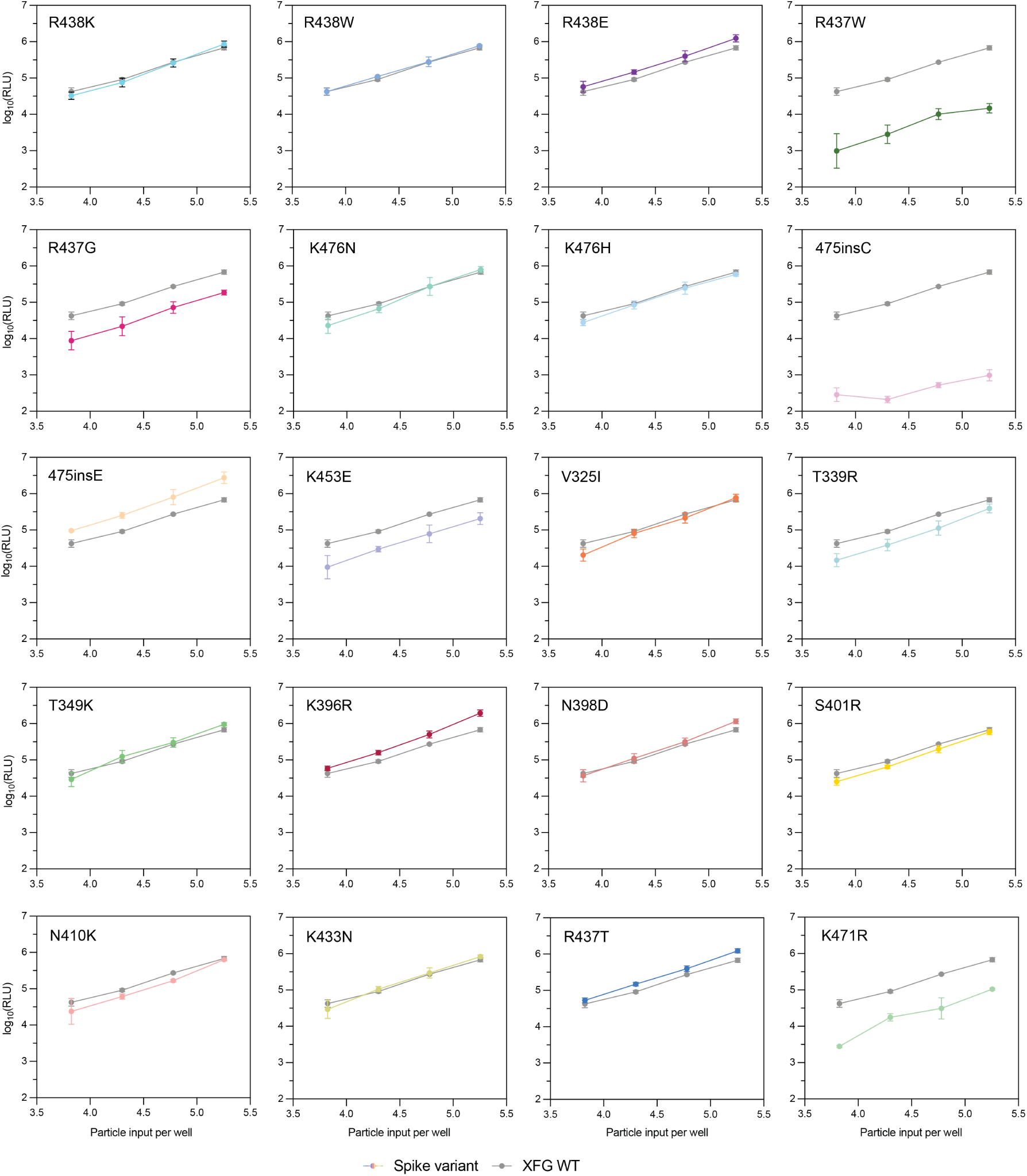
Particle-normalized infectivity assay of antiGen-predicted SARS-CoV-2 XFG spike variants. Four-point, 3-fold serial-dilution infectivity assays were performed using pseudovirus inputs normalized by flow virometry-derived particle concentrations. Colored lines indicate each XFG spike variant, while gray lines show the XFG WT control. Particle inputs were selected such that the central dilution points fell within the shared linear operating window defined across variants, enabling comparison of infectivity at equivalent particle input. Points show mean values and error bars indicate standard deviation (n = 3).

**Figure S11.**
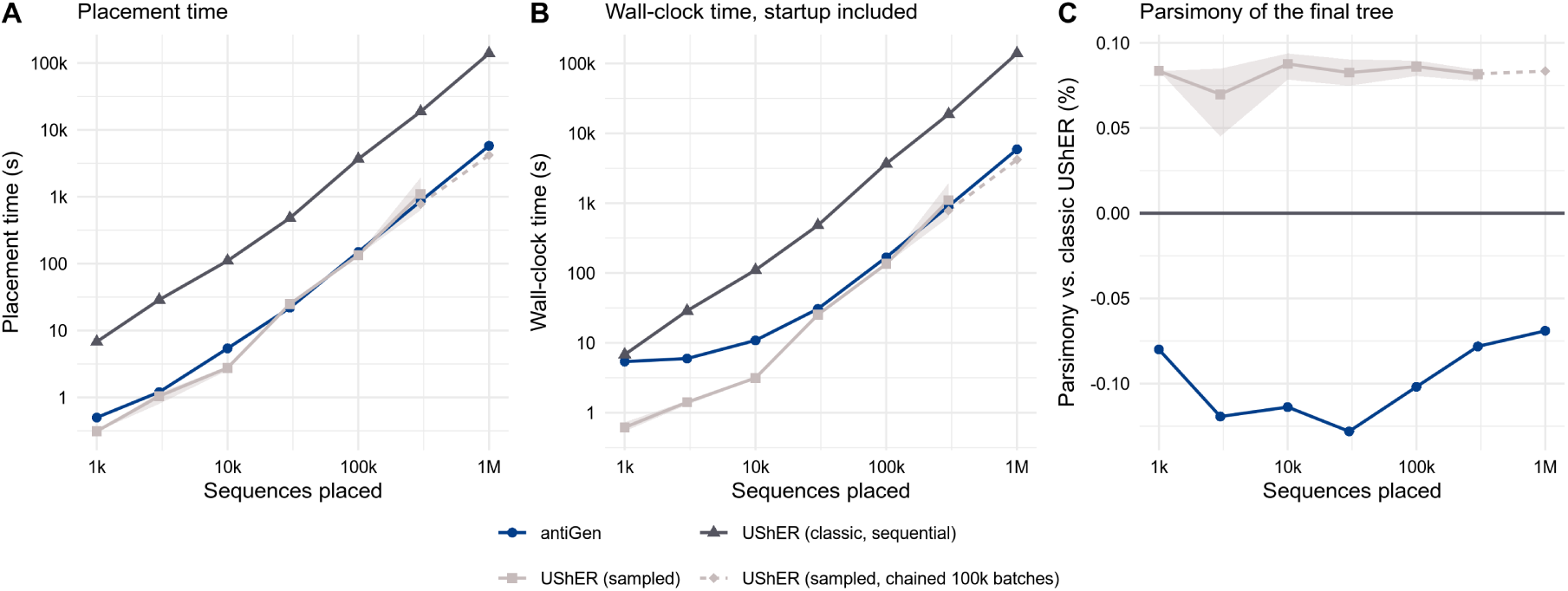
antiGen’s training data generator versus UShER. (A, **B)** Runtime and **(C)** parsimony score comparison between antiGen’s data generation pipeline and UShER (Turakhia et al., 2021) run two ways: classic mode, in which each sample is placed sequentially, and “sampled,” in which sequences can be batched and placed in parallel (among other runtime optimizations). The data used to benchmark both tools are the Nextstrain SARS-CoV-2 tree used throughout this paper, together with sequences obtained from the University of California, Santa Cruz SARS-CoV-2 genome browser (Casper et al., 2026). Both tools were run on whole-genome nucleotide sequences. All runtime data assumes 32 parallel CPU threads. For usher-sampled, the run with 1 million samples crashed upon launch (a “node id Mismatch” error, then a segfault), so we split the same chronological placement order into consecutive 100k batches and ran them in sequence (hence the single “sampled, chained 100k batches” data point for UShER).

**Figure S12.**
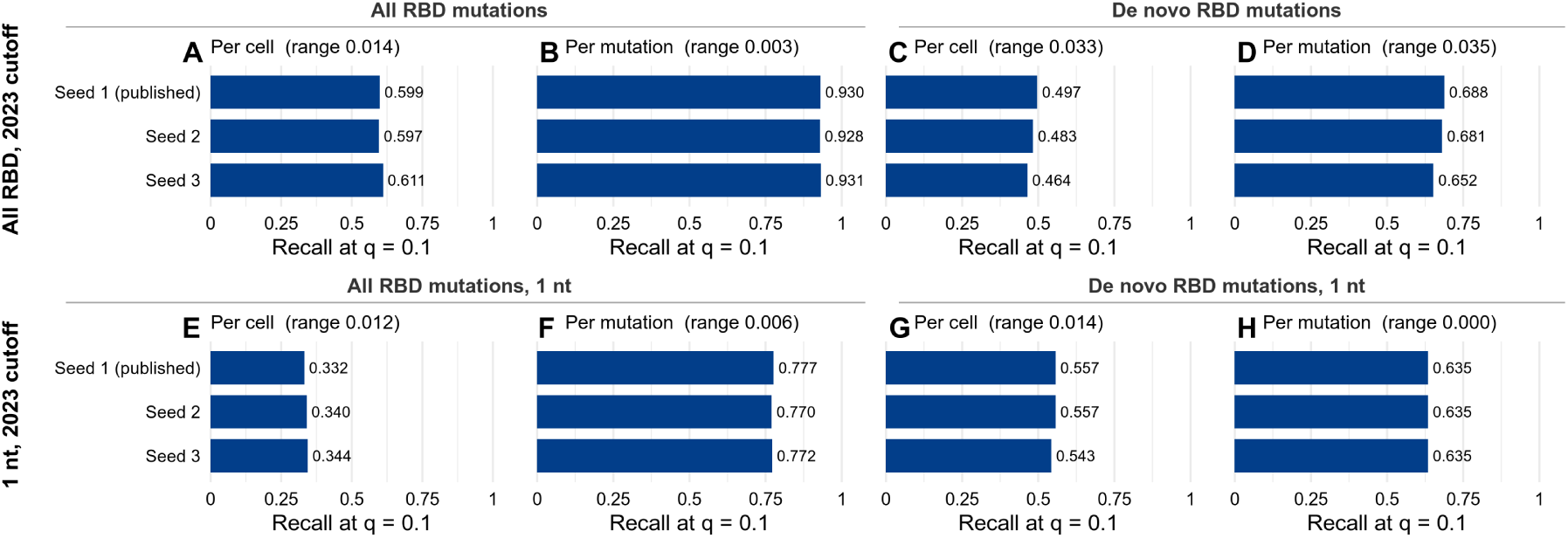
Robustness to random seeding in the training data generation pipeline. antiGen bars from Figure 2C**–J** regenerated under three different random seeds used for the training data generation pipeline. Testing examples remained constant to isolate the effect of variability in the training examples.

**Figure S13.**
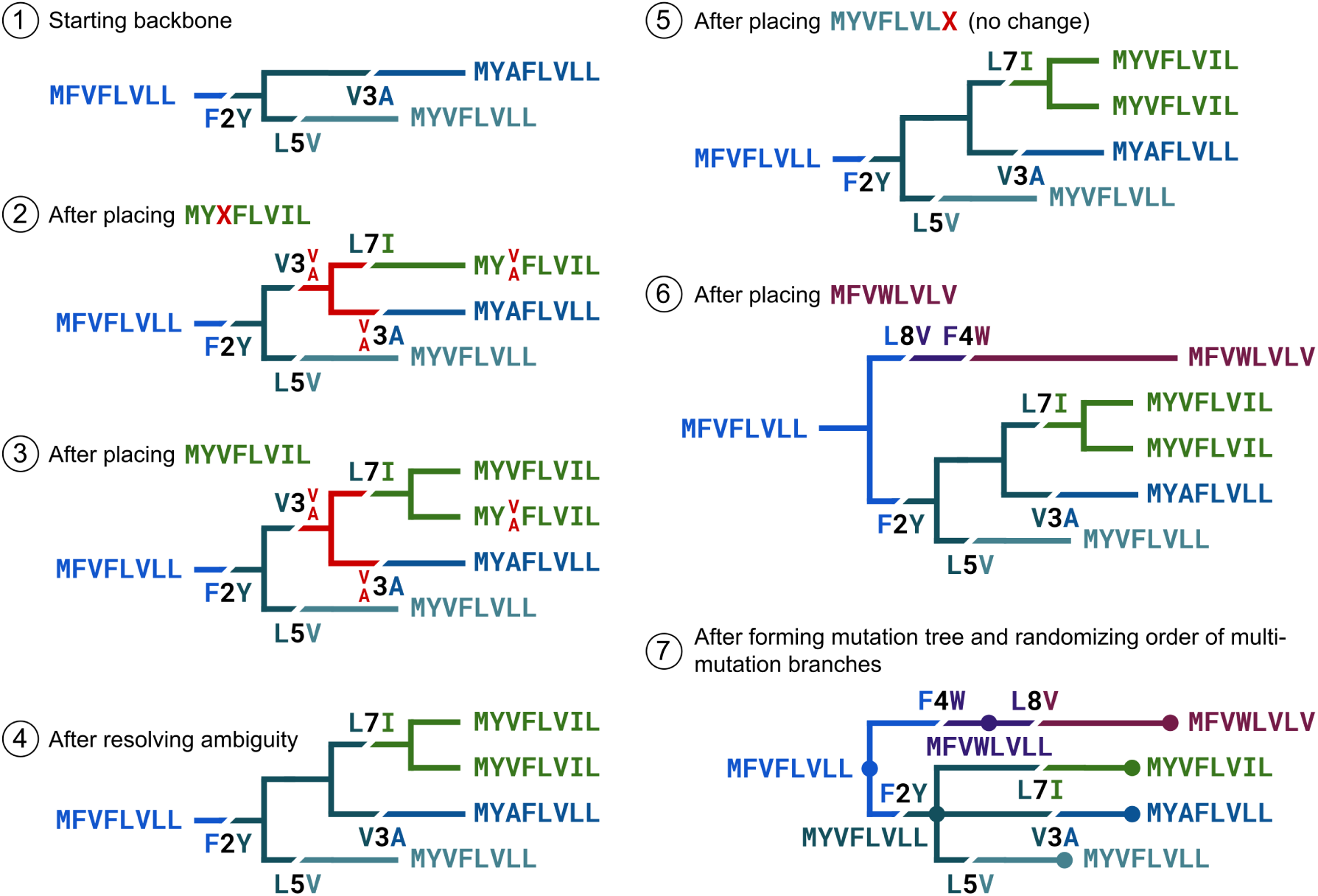
Worked example of the tree augmentation and training data generation algorithms. (1) We start with a backbone phylogenetic tree (e.g., obtained from Nextstrain) and run Hartigan’s parsimony to construct the set of possible parsimonious alleles at each site for internal node (in this simple case, there is only one possible parsimonious configuration, which is shown). (2) Suppose we first attach sequence `MYXFLVIL`, with `X` denoting an ambiguous amino acid. This sequence can be placed parsimoniously in any region of the tree corresponding to the `MYVFLVLL` or `MYAFLVLL` genotype, but whether it lies on one side or the other of the `V3A` mutation cannot be determined due to the ambiguity at site 3. Hence, the new node created upon attachment records that either `V` or `A` can exist at site 3. Moreover, the `X` in the newly-attached leaf is also updated to reflect these two possible choices. (3) Suppose we next attach sequence `MYVFLVIL`. The only parsimonious attachment location is as a sibling of `MYXFLVIL`, downstream of the `L7I` mutation. (4) Following this attachment, there is now only one possible parsimonious state for the `V`/`A` ambiguity at site 3, that being the `V` state. We update the tree to reflect this. (5) Next, suppose we attach `MYVFLVLX`. At all unambiguous positions, this sequence matches that of an existing node; hence, attachment is skipped (as it would not modify the mutation tree). (6) Finally, suppose we attach `MFVWLVLV`. This sequence lands downstream of the root on a branch with two mutations. (7) To derive training examples, we collapse branches with zero mutations and randomly reorder the mutations on branches containing several. We then explode branches with multiple mutations into multiple branches each containing one mutation. Each training example consists of a genotype ancestral to the sample together with the mutations inferred to have altered that genotype, corresponding to the nodes (circles) and their respective descendant edges (lines / elbows) in the resulting graph.

**Figure S14.**
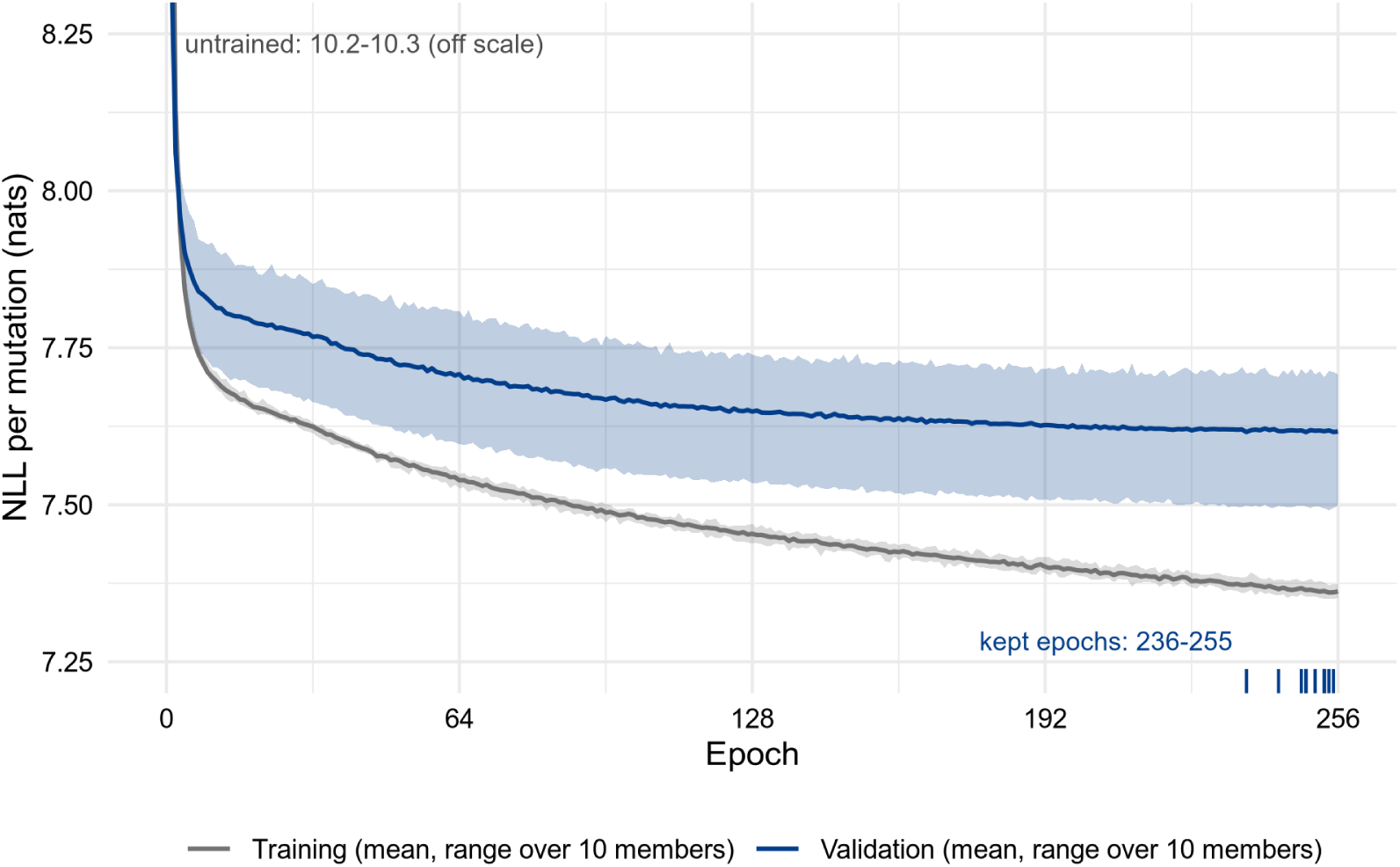
Loss curves for antiGen. antiGen’s training and validation loss on the SARS-CoV-2 dataset with January 1st, 2023 cutoff. Bands reflect the distribution over the ten separate training runs used for cross-validation.

**Figure S15.**
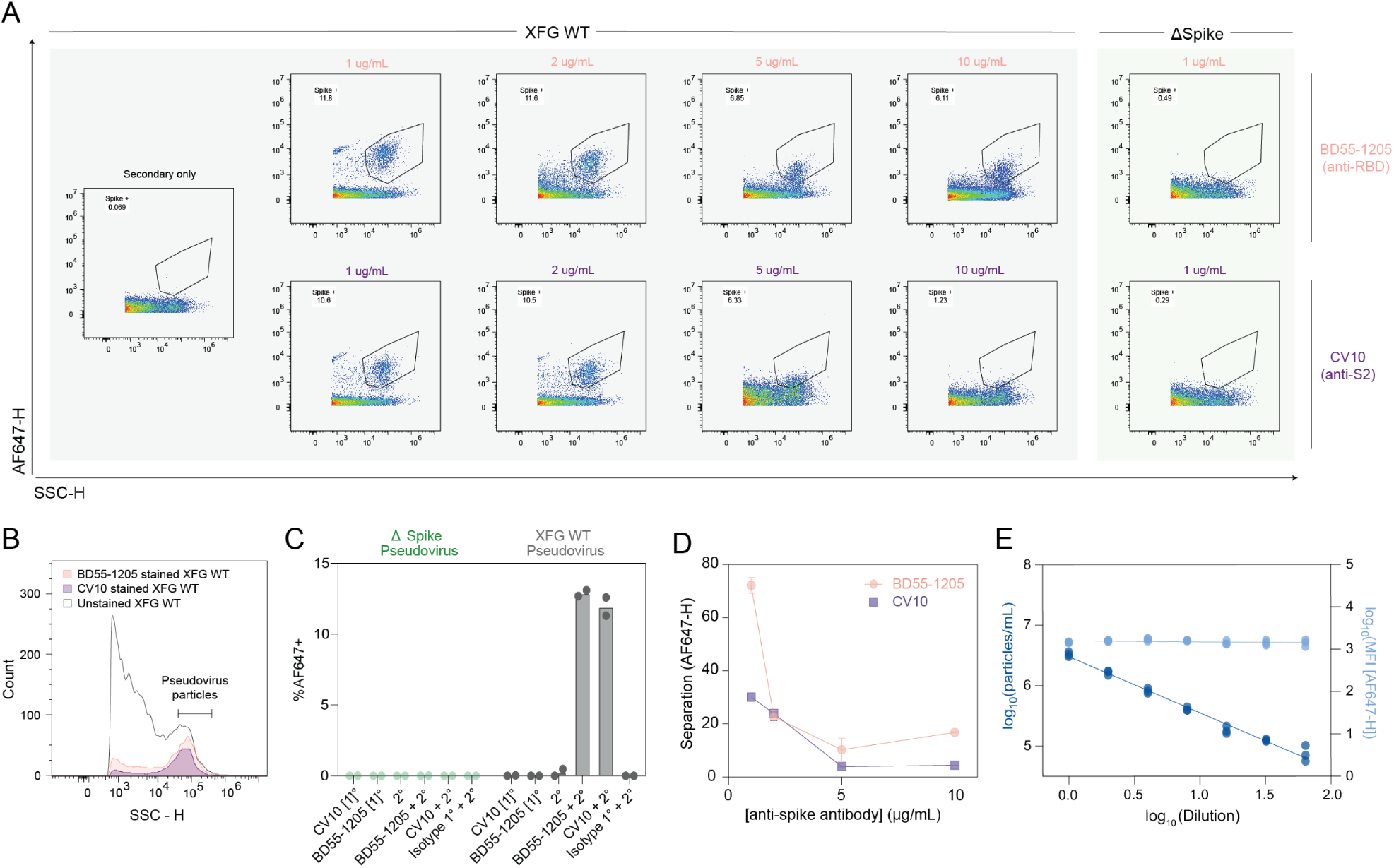
Optimization of flow virometric detection of intact SARS-CoV-2 spike on pseudotyped lentivirus particles. **(A)** Flow virometry detection of XFG WT spike-pseudotyped particles stained with titrated amounts of BD55-1205 anti-RBD or CV10 anti-S2 primary antibodies, followed by AF647-conjugated secondary antibody. Secondary-only and ΔSpike pseudovirus controls were used to define background staining and spike-positive gates. **(B)** SSC-H distributions of unstained, BD55-1205-stained, and CV10-stained XFG WT pseudovirus preparations. Spike-positive particles identified by S2 or RBD staining localized predominantly within the predefined pseudovirus particle gate, supporting enrichment of spike-bearing particles within the SSC-defined population. **(C)** Validation of the spike-positive gate using primary-only, secondary-only, and isotype control conditions. AF647+ events within the defined gate were observed only when both BD55-1205 or CV10 primary and AF647-conjugated secondary antibodies were applied to XFG WT particles, and not to ΔSpike particles, confirming that the gate specifically reports spike-dependent, antibody-specific staining. **(D)** Primary antibody titration for BD55-1205 and CV10 staining of XFG WT pseudovirus particles. Staining performance was evaluated using the stain index, a measure of separation between AF647-positive and AF647-negative particle populations, to identify the primary antibody concentration providing the greatest separation from background (n = 2). **(E)** Effect of pseudovirus dilution on particle recovery and spike-staining intensity. SSC-gated particle concentration decreased proportionally with dilution, whereas the AF647-H median fluorescence intensity of the antibody-positive population remained stable across the tested range. The stability of AF647 fluorescence intensity across serial dilutions indicates that fluorescence measurements were not substantially influenced by particle concentration within the validated operating range, consistent with single-particle detection. Together with the localization of antibody-positive particles within the predefined SSC gate and the specificity controls, these data support the use of SSC-based particle counts as a proxy for pseudovirus particle input in subsequent normalization experiments (n = 3).

**Figure S16.**
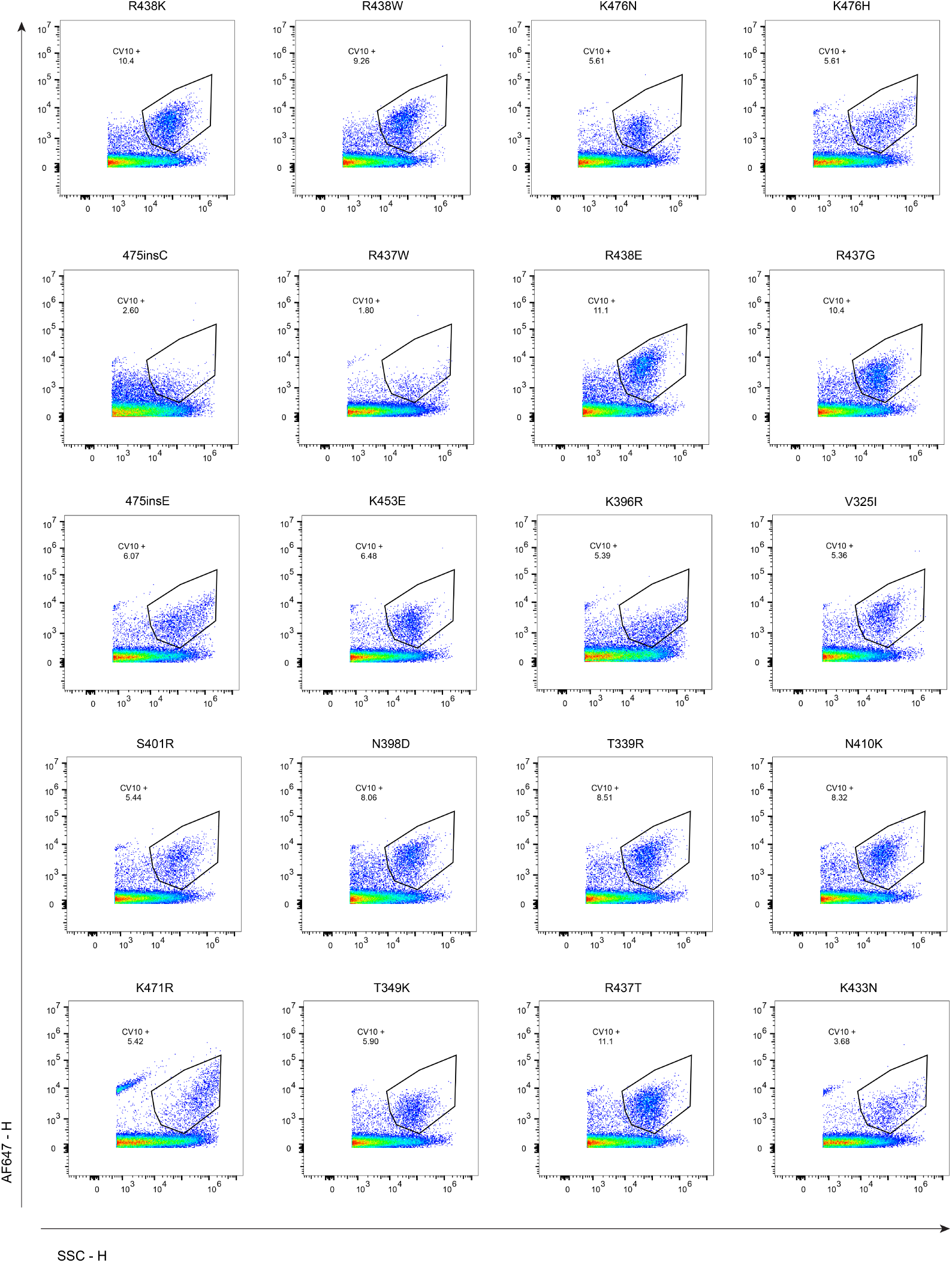
Anti-S2 CV10 staining of spike-bearing particles across XFG spike variants. Representative flow-virometry plots of 10× concentrated pseudovirus preparations stained with CV10, an S2-specific monoclonal antibody, followed by AF647-conjugated goat anti-human IgG secondary antibody detection. AF647-positive populations were identified using a common fluorescence gate established from secondary-only controls. The percentage of CV10-positive events is indicated for XFG WT and each antiGen-predicted spike variant (n = 2).

**Figure S17.**
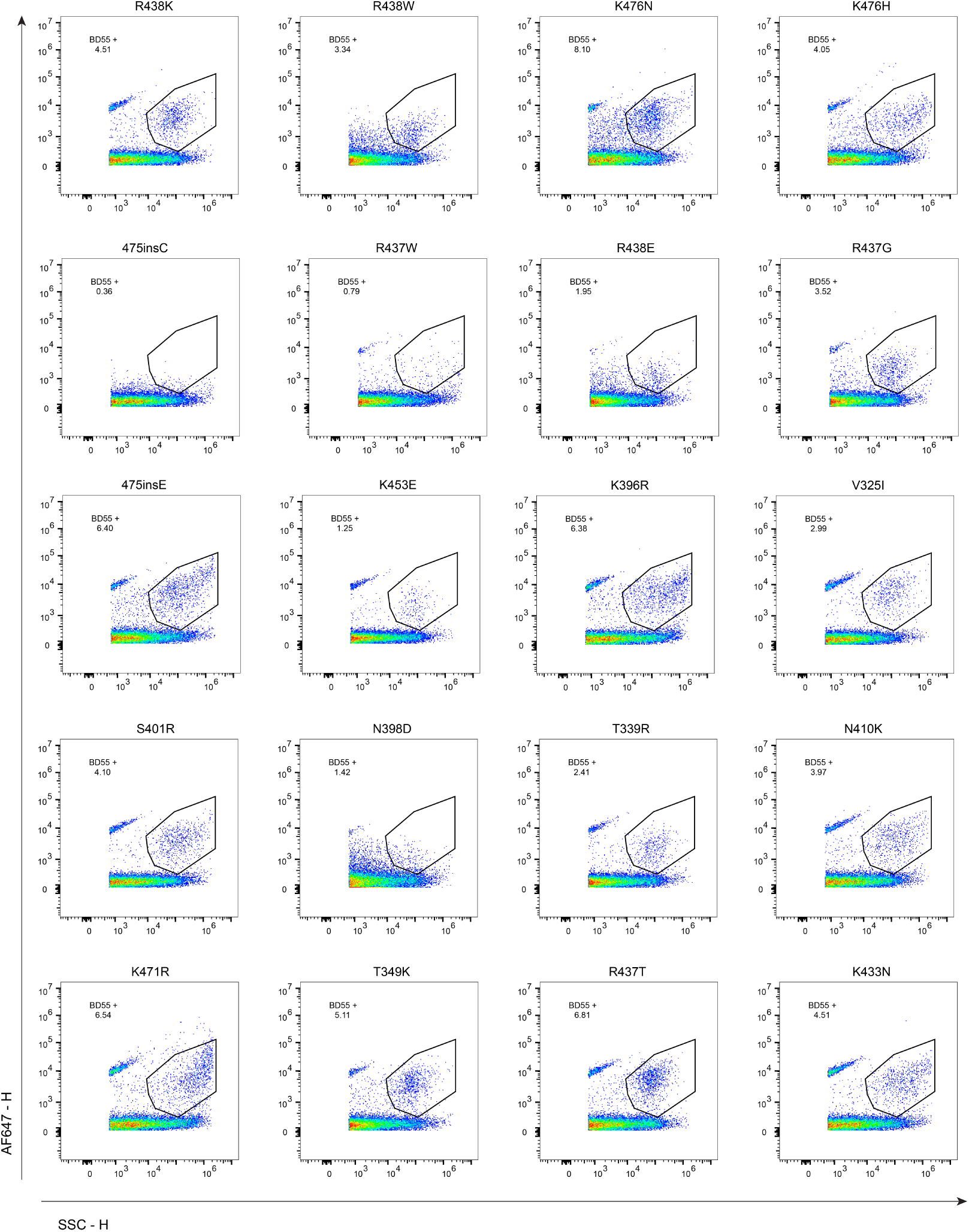
Anti-RBD BD55-1205 staining of spike-bearing particles across XFG spike variants. Representative flow-virometry plots of 10× concentrated pseudovirus preparations stained with BD55-1205, an anti-RBD monoclonal antibody, followed by AF647-conjugated goat anti-human IgG secondary antibody detection. AF647-positive populations were identified using a common fluorescence gate established from secondary-only controls. The percentage of BD55-1205-positive events is indicated for each spike variant (n = 2).

**Figure S18.**
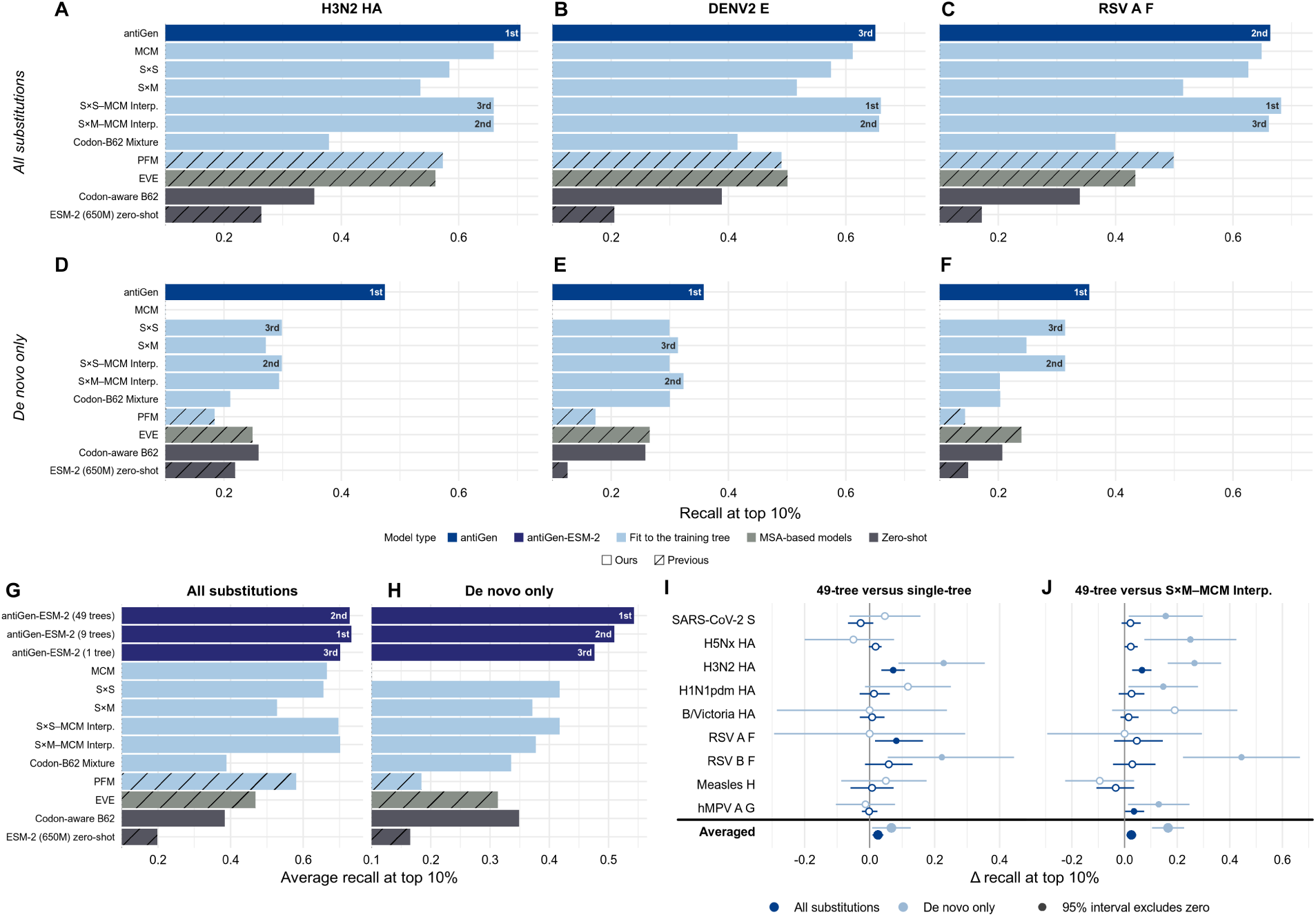
Mutation forecasting for endemic viruses and multi-tree training, restricted to single-nucleotide mutations. We reproduced Figure 4A**–F****, H–K** with top-decile recall restricted to amino acid changes accessible by a single nucleotide substitution.

### Algorithm S1

**Phylogenetic Tree Augmentation**

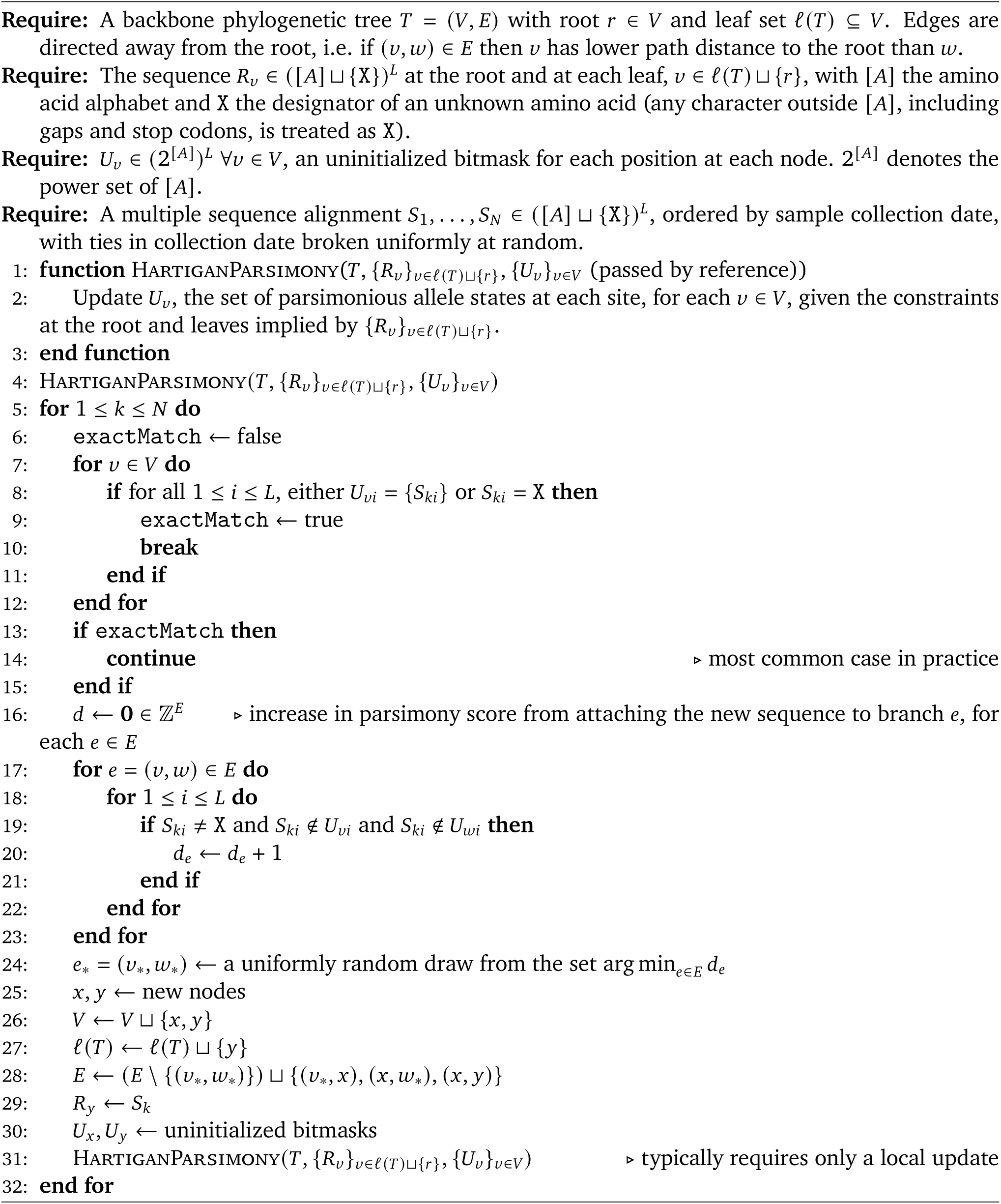

### Algorithm S2

**Example Construction given an Augmented Tree**

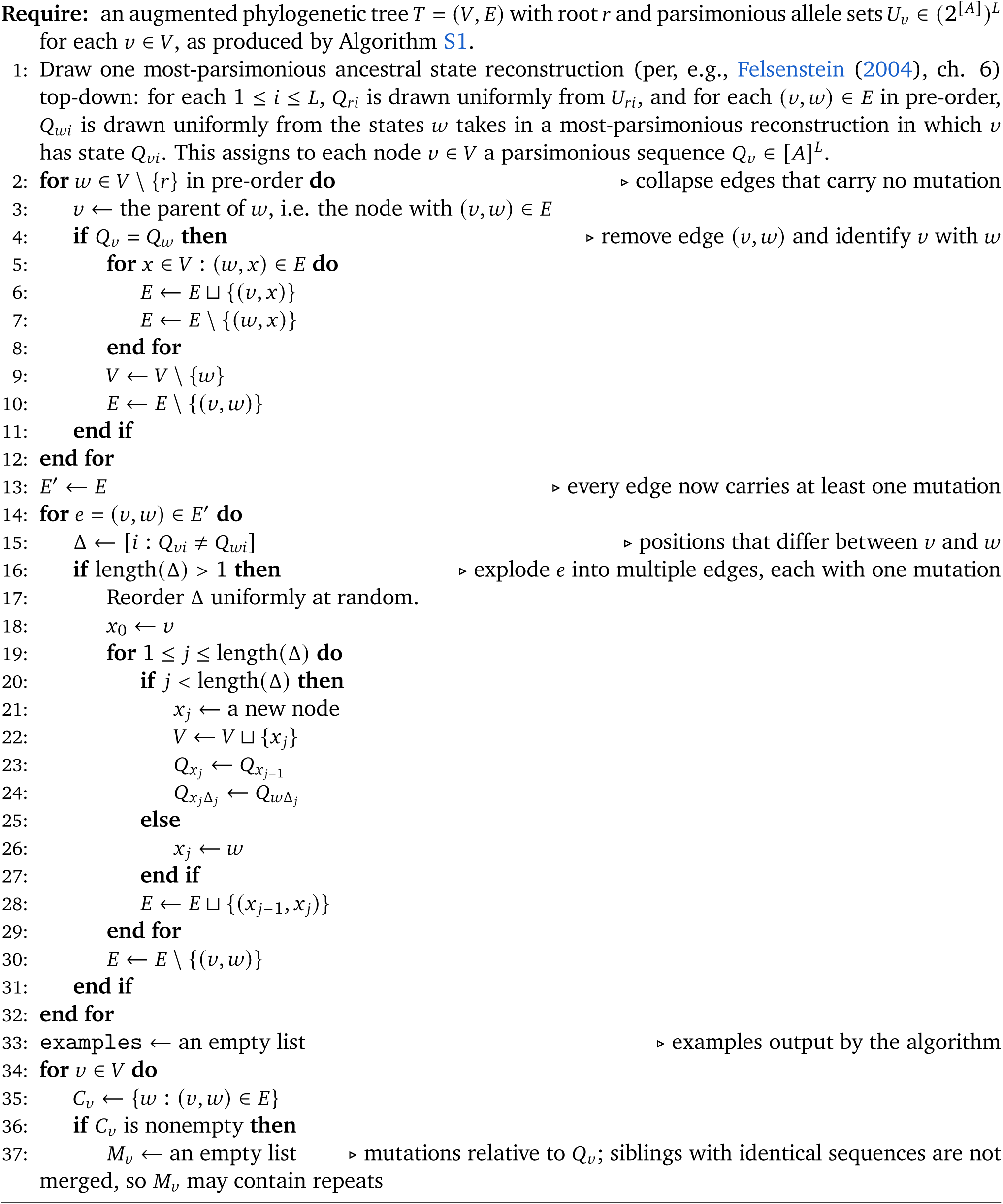
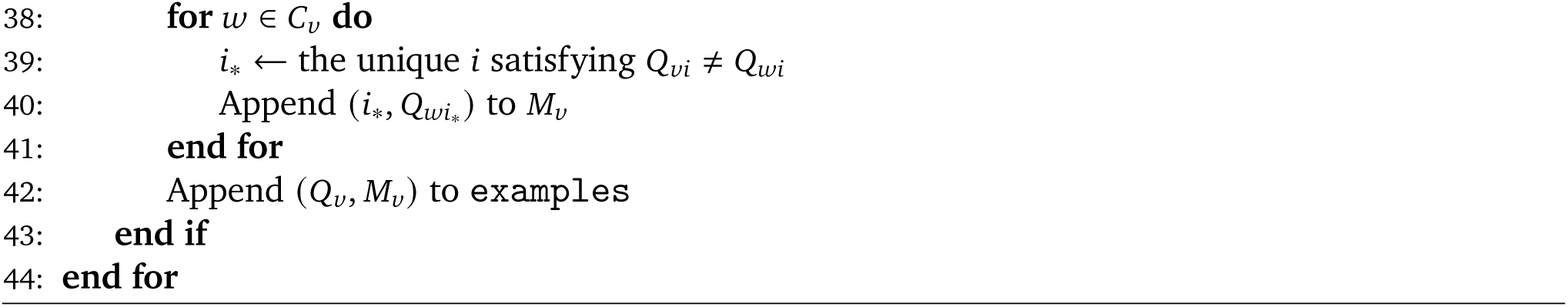

**Table S2.** Nextstrain trees for the nine-virus panel. Each training tree is the dataset’s latest build published on or before 2025-01-01 (*`dataset`* @2025-01-01); each test tree is the current build at download.

| Viral protein | Split | Nextstrain dataset | Build date | Downloaded |
|---|---|---|---|---|
| H5Nx HA | Train | avian-flu/h5nx/ha/all-time@2025-01-01 | 2024-12-06 | 2026-10-17 |
|  | Test | avian-flu/h5nx/ha/all-time | 2026-09-28 | 2026-10-16 |
| SARS-CoV-2 S | Train | ncov/open/global/all-time@2025-01-01 | 2024-12-27 | 2026-10-17 |
|  | Test | ncov/open/global/all-time | 2026-10-11 | 2026-10-16 |
| RSV A F | Train | rsv/a/F/all-time@2025-01-01 | 2024-12-24 | 2026-10-17 |
|  | Test | rsv/a/F/all-time | 2026-10-13 | 2026-10-16 |
| RSV B F | Train | rsv/b/F/all-time@2025-01-01 | 2024-12-24 | 2026-10-17 |
|  | Test | rsv/b/F/all-time | 2026-10-13 | 2026-10-16 |
| Measles H | Train | measles/genome/global@2025-01-01 | 2025-01-01 | 2026-10-17 |
|  | Test | measles/genome/global | 2026-10-13 | 2026-10-16 |
| H3N2 HA | Train | seasonal-flu/h3n2/ha/12y@2025-01-01 | 2024-12-29 | 2026-10-16 |
|  | Test | seasonal-flu/h3n2/ha/12y | 2026-10-08 | 2026-10-16 |
| H1N1pdm HA | Train | seasonal-flu/h1n1pdm/ha/12y@2025-01-01 | 2024-12-29 | 2026-10-16 |
|  | Test | seasonal-flu/h1n1pdm/ha/12y | 2026-10-08 | 2026-10-16 |
| B/Victoria HA | Train | seasonal-flu/vic/ha/12y@2025-01-01 | 2024-12-29 | 2026-10-16 |
|  | Test | seasonal-flu/vic/ha/12y | 2026-10-08 | 2026-10-16 |
| hMPV A G | Train | hmpv/a/G@2025-01-01 | 2024-12-23 | 2026-10-16 |
|  | Test | hmpv/a/G | 2026-10-14 | 2026-10-16 |

**Table S3.** The forty additional Nextstrain trees in the 49-tree training corpus (training only). Each is the dataset’s latest build published on or before 2025-01-01 (*`dataset`* @2025-01-01).

| Viral protein | Nextstrain dataset | Build date | Downloaded |
|---|---|---|---|
| H5N1 (cattle outbreak) HA | avian-flu/h5n1-cattle-outbreak/genome@2025-01-01 | 2024-12-31 | 2026-10-26 |
| H5N1 NA | avian-flu/h5n1/na/all-time@2025-01-01 | 2024-12-06 | 2026-10-16 |
| H5Nx NA | avian-flu/h5nx/na/all-time@2025-01-01 | 2024-12-06 | 2026-10-16 |
| H7N9 HA | avian-flu/h7n9/ha/all-time@2025-01-01 | 2024-12-06 | 2026-10-16 |
| H7N9 NA | avian-flu/h7n9/na/all-time@2025-01-01 | 2024-12-06 | 2026-10-16 |
| H9N2 HA | avian-flu/h9n2/ha/all-time@2025-01-01 | 2024-12-06 | 2026-10-16 |
| H9N2 NA | avian-flu/h9n2/na/all-time@2025-01-01 | 2024-12-06 | 2026-10-17 |
| DENV1 E | dengue/denv1/E@2025-01-01 | 2025-01-01 | 2026-10-17 |
| DENV1 pr | dengue/denv1/genome@2025-01-01 | 2025-01-01 | 2026-10-26 |
| DENV2 E | dengue/denv2/E@2025-01-01 | 2025-01-01 | 2026-10-17 |
| DENV2 pr | dengue/denv2/genome@2025-01-01 | 2025-01-01 | 2026-10-26 |
| DENV3 E | dengue/denv3/E@2025-01-01 | 2025-01-01 | 2026-10-17 |
| DENV4 E | dengue/denv4/E@2025-01-01 | 2025-01-01 | 2026-10-17 |
| DENV4 pr | dengue/denv4/genome@2025-01-01 | 2025-01-01 | 2026-10-26 |
| EV-D68 VP2 | enterovirus/d68/genome@2025-01-01 | 2024-07-23 | 2026-10-26 |
| EV-D68 VP3 | enterovirus/d68/genome@2025-01-01 | 2024-07-23 | 2026-10-26 |
| EV-D68 VP1 | enterovirus/d68/vp1@2025-01-01 | 2024-05-29 | 2026-10-16 |
| hMPV B F | hmpv/b/F@2025-01-01 | 2024-12-23 | 2026-10-16 |
| hMPV B SH | hmpv/b/genome@2025-01-01 | 2024-12-23 | 2026-10-26 |
| Lassa GPC | lassa/gpc@2025-01-01 | 2024-12-09 | 2026-10-16 |
| Measles F | measles/genome/global@2025-01-01 | 2025-01-01 | 2026-10-17 |
| Mumps F | mumps/global@2025-01-01 | 2024-03-21 | 2026-10-16 |
| Mumps HN | mumps/global@2025-01-01 | 2024-03-21 | 2026-10-16 |
| Mumps SH | mumps/global@2025-01-01 | 2024-03-21 | 2026-10-16 |
| Oropouche M polyprotein | oropouche/M@2025-01-01 | 2024-11-20 | 2026-10-16 |
| Rabies G | rabies@2025-01-01 | 2025-01-01 | 2026-10-16 |
| RSV A G | rsv/a/genome/all-time@2025-01-01 | 2024-12-24 | 2026-10-26 |
| RSV A SH | rsv/a/genome/all-time@2025-01-01 | 2024-12-24 | 2026-10-26 |
| RSV B G | rsv/b/G/all-time@2025-01-01 | 2024-12-24 | 2026-10-16 |
| RSV B SH | rsv/b/genome/all-time@2025-01-01 | 2024-12-24 | 2026-10-26 |
| HCoV-229E S | seasonal-cov/229e@2025-01-01 | 2025-01-01 | 2026-10-16 |
| HCoV-HKU1 S | seasonal-cov/hku1@2025-01-01 | 2025-01-01 | 2026-10-16 |
| HCoV-NL63 S | seasonal-cov/nl63@2025-01-01 | 2025-01-01 | 2026-10-16 |
| HCoV-OC43 S | seasonal-cov/oc43@2025-01-01 | 2025-01-01 | 2026-10-16 |
| H1N1pdm NA | seasonal-flu/h1n1pdm/na/3y@2025-01-01 | 2024-12-29 | 2026-10-26 |
| H3N2 NA | seasonal-flu/h3n2/na/12y@2025-01-01 | 2024-12-29 | 2026-10-16 |
| B/Victoria NA | seasonal-flu/vic/na/3y@2025-01-01 | 2024-12-29 | 2026-10-26 |
| B/Yamagata HA | seasonal-flu/yam/ha/12y@2025-01-01 | 2024-02-14 | 2026-10-16 |
| B/Yamagata NA | seasonal-flu/yam/na/6y@2025-01-01 | 2024-02-14 | 2026-10-26 |
| Yellow fever E | yellow-fever/genome@2025-01-01 | 2024-12-16 | 2026-10-26 |

## Notes

https://github.com/evo-design/antiGen

https://doi.org/10.5281/zenodo.21481224

### Summary of Updates

SARS-CoV-2 mutation forecasting section updated to include prediction of prevalent variants and control for codon distance Endemic viruses section updated to include multi-tree finetuning of ESM-2 Training and testing data generation pipeline updated to handle genotypic uncertainty and better guard against leakage antiGen training regime updated to support validation-based checkpoint selection and cross-validation New statistical baselines developed to compare against antiGen

## References

Abramson, J., Adler, J., Dunger, J., Evans, R., Green, T., Pritzel, A., Ronneberger, O., Willmore, L., Ballard, A., Bambrick, J., Bodenstein, S., Evans, D., Hung, C., O’Neill, M., Reiman, D., Tunyasuvunakool, K., Wu, Z., Žemgulytė, A., Arvaniti, E., Beattie, C., Bertolli, O., Bridgland, A., Cherepanov, A., Congreve, M., Cowen-Rivers, A., Cowie, A., Figurnov, M., Fuchs, F., Gladman, H., Jain, R., Khan, Y., Low, C., Perlin, K., Potapenko, A., Savy, P., Singh, S., Stecula, A., Thillaisundaram, A., Tong, C., Yakneen, S., Zhong, E., Zielinski, M., Žídek, A., Bapst, V., Kohli, P., Jaderberg, M., Hassabis, D., and Jumper, J. (2024). Accurate structure prediction of biomolecular interactions with alphafold 3. Nature, 630(8016):493–500.

Aksamentov, I., Roemer, C., Hodcroft, E. B., and Neher, R. A. (2021). Nextclade: clade assignment, mutation calling and quality control for viral genomes. Journal of Open Source Software, 6(67):3773.

Altschul, S. F., Gish, W., Miller, W., Myers, E. W., and Lipman, D. J. (1990). Basic local alignment search tool. Journal of Molecular Biology, 215(3):403–410.

Arakelyan, A., Fitzgerald, W., Margolis, L., and Grivel, J.-C. (2013). Nanoparticle-based flow virometry for the analysis of individual virions. Journal of Clinical Investigation, 123(9):3716–3727.

Berman, D. S., Howser, C., Mehoke, T., Ernlund, A. W., and Evans, J. D. (2023). Mutagan: A sequenceto-sequence gan framework to predict mutations of evolving protein populations. Virus Evolution, 9(1).

Bloom, J. D. and Neher, R. A. (2023). Fitness effects of mutations to sars-cov-2 proteins. Virus Evolution, 9(2):vead055.

Burnie, J., Ouano, C., Luo, V., Dzuvor, C. K., Miller, T., Ospina, G., Tanneti, N. S., Tan, L. H., Hamel, D. J., Hammond, C., Matthews, H., Evanson, L. R., Joseph, J., Moak, S. P., Kanki, P., Cohen, N. A., Weiss, S. R., and Corbett-Helaire, K. S. (2026). Evaluating spike antigenicity across endemic human coronavirus models using flow virometry.

Carabelli, A. M., Peacock, T. P., Thorne, L. G., Harvey, W. T., Hughes, J., COVID-19 Genomics UK Consortium, Peacock, S. J., Barclay, W. S., de Silva, T. I., Towers, G. J., and Robertson, D. L. (2023). SARS-CoV-2 variant biology: immune escape, transmission and fitness. Nature Reviews Microbiology, 21(3):162–177.

Casper, J., Speir, M. L., Raney, B. J., Perez, G., Nassar, L. R., Lee, C. M., Hinrichs, A. S., Gonzalez, J. N., Fischer, C., Diekhans, M., Clawson, H., Benet-Pages, A., Barber, G. P., Vaske, C. J., van Baren, M. J., Wang, K., Rodriguez, Y. J. P., Jenkins-Kiefer, J. A., Chalamala, M., Haussler, D., Kent, W. J., and Haeussler, M. (2026). The ucsc genome browser database: 2026 update. Nucleic Acids Research, 54(D1):D1331–D1335.

Chen, C., Taepper, A., Engelniederhammer, F., Kellerer, J., Roemer, C., and Stadler, T. (2023). Lapis is a fast web api for massive open virus sequencing data. BMC Bioinformatics, 24(232).

Cock, P. J. A., Antao, T., Chang, J. T., Chapman, B. A., Cox, C. J., Dalke, A., Friedberg, I., Hamelryck, T., Kauff, F., Wilczynski, B., and de Hoon, M. J. L. (2009). Biopython: freely available python tools for computational molecular biology and bioinformatics. Bioinformatics, 25(11):1422–1423.

Crawford, K. H. D., Eguia, R., Dingens, A. S., Loes, A. N., Malone, K. D., Wolf, C. R., Chu, H. Y., Tortorici, M. A., Veesler, D., Murphy, M., Pettie, D., King, N. P., Balazs, A. B., and Bloom, J. D. (2020). Protocol and reagents for pseudotyping lentiviral particles with sars-cov-2 spike protein for neutralization assays. Viruses, 12(5):513.

Dadonaite, B., Crawford, K. H. D., Radford, C. E., Farrell, A. G., Yu, T. C., Hannon, W. W., Zhou, P., Andrabi, R., Burton, D. R., Liu, L., Ho, D. D., Chu, H. Y., Neher, R. A., and Bloom, J. D. (2023). A pseudovirus system enables deep mutational scanning of the full SARS-CoV-2 spike. Cell, 186(6):1263– 1278.e20.

Dagum, L. and Menon, R. (1998). Openmp: An industry-standard api for shared-memory programming. IEEE Comput. Sci. Eng., 5(1):46–55.

Dalla Vecchia, E. (2024). Pathoplexus: towards fair and transparent sequence sharing. The Lancet Microbe, 5(12):100995.

Diaz-Uriarte, R. and Johnston, I. G. (2025). A picture guide to cancer progression and evolutionary accumulation models: Systematic critique, plausible interpretations, and alternative uses. IEEE Access, 13:62306–62340.

Ding, F. and Steinhardt, J. (2024). Protein language models are biased by unequal sequence sampling across the tree of life. In ICLR 2024 Workshop on Generative and Experimental Perspectives for Biomolecular Design. Exec. Order No. 14,292 (2025). Improving the safety and security of biological research. 90 Fed. Reg. 19,611.

Felsenstein, J. (2004). Inferring phylogenies, volume 2. Sinauer associates Sunderland, MA.

Fitch, W. M. (1971). Toward defining the course of evolution: Minimum change for a specific tree topology. Systematic Biology, 20(4):406–416.

Fraser, J., Notin, P., Dias, M., Gomez, A., Min, J. K., Brock, K., Gal, Y., and Marks, D. S. (2021). Disease variant prediction with deep generative models of evolutionary data. Nature, 599:91–95.

Geoghegan, J. L. and Holmes, E. C. (2018). Learning from prepandemic data to forecast viral escape. Nature Reviews Genetics, 19:756–769.

Gordon, C. W., Lu, A. X., and Abbeel, P. (2025). Protein language model fitness is a matter of preference. In The Thirteenth International Conference on Learning Representations.

Gretton, D., Wang, B., Edison, R., Foner, L., Berlips, J., Vogel, T., Kysel, M., Chen, W., Sage-Ling, F., Van Hauwe, L., Wooster, S., Cozzarini, H., Weinstein-Raun, B., DeBenedictis, E. A., Liu, A. B., Chory, E., Cui, H., Li, X., Dong, J., Fabrega, A., Dennison, C., Don, O., Ye, C. T., Uberoy, K., Rivest, R. L., Gao, M., Yu, Y., Baum, C., Damgard, I., Yao, A. C., and Esvelt, K. M. (2025). Exact-match search with functional variant prediction enables automated dna screening. bioRxiv.

Grubaugh, N. D., Ladner, J. T., Lemey, P., Pybus, O. G., Rambaut, A., Holmes, E. C., and Andersen, K. G. (2019). Tracking virus outbreaks in the twenty-first century. Nature Microbiology, 4:10–19.

Hadfield, J., Megill, C., Bell, S. M., Huddleston, J., Potter, B., Callender, C., Sagulenko, P., Bedford, T., and Neher, R. A. (2018). Nextstrain: real-time tracking of pathogen evolution. Bioinformatics, 34(23):4121–4123.

Harris, C. R., Millman, K. J., van der Walt, S. J., Gommers, R., Virtanen, P., Cournapeau, D., Wieser, E., Taylor, J., Berg, S., Smith, N. J., Kern, R., Picus, M., Hoyer, S., van Kerkwijk, M. H., Brett, M., Haldane, A., del Río, J. F., Wiebe, M., Peterson, P., Gérard-Marchant, P., Sheppard, K., Reddy, T., Weckesser, W., Abbasi, H., Gohlke, C., and Oliphant, T. E. (2020). Array programming with NumPy. Nature, 585(7825):357–362.

Hartigan, J. A. (1973). Minimum mutation fits to a given tree. Biometrics, 29(1):53–65.

Henikoff, S. and Henikoff, J. G. (1992). Amino acid substitution matrices from protein blocks. Proceedings of the National Academy of Sciences, 89(22):10915–10919.

Hie, B., Zhong, E. D., Berger, B., and Bryson, B. (2021). Learning the language of viral evolution and escape. Science, 371(6526):284–288.

Hopf, T. A., Ingraham, J. B., Poelwijk, F. J., Schärfe, C. P. I., Springer, M., Sander, C., and Marks, D. S. (2017). Mutation effects predicted from sequence co-variation. Nature Biotechnology, 35:128–135.

Huddleston, J., Barnes, J. R., Rowe, T., Xu, X., Kondor, R., Wentworth, D. E., Whittaker, L., Ermetal, B., Daniels, R. S., McCauley, J. W., Fujisaki, S., Nakamura, K., Kishida, N., Watanabe, S., Hasegawa, H., Barr, I., Subbarao, K., Barrat-Charlaix, P., Neher, R. A., and Bedford, T. (2020). Integrating genotypes and phenotypes improves long-term forecasts of seasonal influenza a/h3n2 evolution. eLife, 9:e60067.

Hunter, J. D. (2007). Matplotlib: A 2d graphics environment. Computing in Science & Engineering, 9(3):90–95.

Ito, J., Strange, A., Liu, W., Joas, G., Lytras, S., to Phenotype Japan (G2P-Japan) Consortium, T. G., and Sato, K. (2025). A protein language model for exploring viral fitness landscapes. Nature Communications, 16.

Jahn, K., Kuipers, J., and Beerenwinkel, N. (2016). Tree inference for single-cell data. Genome Biology, 17.

Jian, F., Wec, A. Z., Feng, L., Yu, Y., Wang, L., Wang, P., Yu, L., Wang, J., Hou, J., Berrueta, D. M., Lee, D., Speidel, T., Ma, L., Kim, T., Yisimayi, A., Song, W., Wang, J., Liu, L., Yang, S., Niu, X., Xiao, T., An, R., Wang, Y., Shao, F., Wang, Y., Pecetta, S., Wang, X., Walker, L. M., and Cao, Y. (2025). Viral evolution prediction identifies broadly neutralizing antibodies to existing and prospective sars-cov-2 variants. Nature Microbiology, 10(8):2003–2017.

Jungbauer-Groznica, M., Commere, P.-H., Planchais, C., Cottignies-Calamarte, A., De Cruz, A., Fantin Rengifo, A., Guivel-Benhassine, F., Staropoli, I., Schmutz, S., Novault, S., Veyer, D., Péré, H., Mouquet, H., Schwartz, O., and Bruel, T. (2026). Detection and characterization of single sars-cov-2 viral particles by flow virometry.

Kingma, D. P. and Ba, J. (2015). Adam: A method for stochastic optimization. In Bengio, Y. and LeCun, Y., editors, 3rd International Conference on Learning Representations, ICLR 2015, San Diego, CA, USA, May 7-9, 2015, Conference Track Proceedings.

Koehl, A., Prillo, S., Liu, M., Xiong, J., Weng, L., Savage, D. F., and Song, Y. S. (2026). Deep models of protein evolution in time generate realistic evolutionary trajectories and functional proteins. bioRxiv.

Levinstein Hallak, K. and Rosset, S. (2022). Statistical modeling of sars-cov-2 substitution processes: predicting the next variant. Communications Biology, 5(1).

Lin, Z., Akin, H., Rao, R., Hie, B., Zhu, Z., Lu, W., Smetanin, N., Verkuil, R., Kabeli, O., Shmueli, Y., dos Santos Costa, A., Fazel-Zarandi, M., Sercu, T., Candido, S., and Rives, A. (2023). Evolutionary-scale prediction of atomic-level protein structure with a language model. Science, 379(6637):1123–1130.

Lu, S. Z., Vermani, A., Sanno, K., Lu, J., Matsen, F. A., Jagota, M., and Song, Y. S. (2026). Conditionally site-independent neural evolution of antibody sequences.

Łuksza, M. and Lässig, M. (2014). A predictive fitness model for influenza. Nature, 507:57–61.

Luo, X. G., Kuipers, J., and Beerenwinkel, N. (2023). Joint inference of exclusivity patterns and recurrent trajectories from tumor mutation trees. Nature Communications, 14.

Maltseva, M. and Langlois, M.-A. (2022). Flow virometry for characterizing the size, concentration, and surface antigens of viruses. Current Protocols, 2(2):e368.

Matsen, F. A., Dumm, W., Sung, K., Johnson, M. M., Rich, D., Starr, T., Song, Y. S., Fukuyama, J., and Haddox, H. K. (2026). Separating selection from mutation in antibody language models. elife.

McKinney, W. (2010). Data Structures for Statistical Computing in Python. In Stéfan van der Walt and Jarrod Millman, editors, Proceedings of the 9th Python in Science Conference, pages 56–61.

Mille-Fragoso, L. S., Driscoll, C. L., Wang, J. N., Dai, H., Widatalla, T., Zhang, J. L., Zhang, X., Rao, B., Feng, L., Hie, B. L., et al. (2026). Efficient generation of epitope-targeted antibodies with germinal. Nature biotechnology, pages 1–10.

National Science and Technology Council (2024). Framework for nucleic acid synthesis screening. Technical report, Office of Science and Technology Policy. Revised September 2024. Prepared by the Fast Track Action Committee on Synthetic Nucleic Acid Procurement Screening.

Neher, R. A., Russell, C. A., and Shraiman, B. I. (2014). Predicting evolution from the shape of genealogical trees. eLife, 3.

Notin, P., Dias, M., Frazer, J., Marchena-Hurtado, J., Gomez, A., Marks, D. S., and Gal, Y. (2022). Tranception: Protein fitness prediction with autoregressive transformers and inference-time retrieval. In Proceedings of the 39th International Conference on Machine Learning. PMLR.

Notin, P., Kollasch, A., Ritter, D., van Niekerk, L., Paul, S., Spinner, H., Rollins, N., Shaw, A., Orenbuch, R., Weitzman, R., Frazer, J., Dias, M., Franceschi, D., Gal, Y., and Marks, D. (2023). Proteingym: Large-scale benchmarks for protein fitness prediction and design. In Oh, A., Naumann, T., Globerson, A., Saenko, K., Hardt, M., and Levine, S., editors, Advances in Neural Information Processing Systems, volume 36, pages 64331–64379. Curran Associates, Inc.

Obermeyer, F., Jankowiak, M., Barkas, N., Schaffner, S. F., Pyle, J. D., Yurkovetskiy, L., Bosso, M., Park, D. J., Babadi, M., MacInnis, B. L., Luban, J., Sabeti, P. C., and Lemieux, J. E. (2022). Analysis of 6.4 million sars-cov-2 genomes identifies mutations associated with fitness. Science, 376(6599):1327– 1332.

Paszke, A., Gross, S., Massa, F., Lerer, A., Bradbury, J., Chanan, G., Killeen, T., Lin, Z., Gimelshein, N., Antiga, L., Desmaison, A., Köpf, A., Yang, E., DeVito, Z., Raison, M., Tejani, A., Chilamkurthy, S., Steiner, B., Fang, L., Bai, J., and Chintala, S. (2019). PyTorch: an imperative style, high-performance deep learning library. Curran Associates Inc., Red Hook, NY, USA.

Powell, A. E., Caruso, H., Park, S., Chen, J.-L., O’Rear, J., Ferrer, B. J., Stieh, D. J., Weiss, A. M., Belnap, D. M., Walker, A., Bruening, A., Hartwig, A., Sprouse, K. R., Addetia, A., Alshukairi, A. N., Ahyong, V., Dougherty, C. S., Veesler, D., Bowen, R., Ledgerwood, J. E., Kay, M. S., Weidenbacher, P. A.-B., and Palanski, B. A. (2026). A stabilized mers-cov spike ferritin nanoparticle vaccine elicits robust and protective neutralizing antibody responses. Nature Communications, 17(1).

Rao, R. M., Liu, J., Verkuil, R., Meier, J., Canny, J., Abbeel, P., Sercu, T., and Rives, A. (2021). Msa transformer. In Proceedings of the 38th International Conference on Machine Learning. PMLR.

Responsible AI for Protein Design Community (2023). Community values, guiding principles, and commitments for the responsible development of AI for protein design. https://responsiblebiodesign.ai/. Accessed: 2026-07-04.

Sanjuán, R. (2010). Mutational fitness effects in rna and single-stranded dna viruses: common patterns revealed by site-directed mutagenesis studies. Philosophical Transactions of the Royal Society B: Biological Sciences, 365(1548):1975–1982.

Sankoff, D. (1975). Minimal mutation trees of sequences. SIAM Journal on Applied Mathematics, 28(1):35–42.

Shu, Y. and McCauley, J. (2017). Gisaid: Global initiative on sharing all influenza data—from vision to reality. Eurosurveillance, 22(13).

Simonich, C. A., McMahon, T. E., Kampman, L., Chu, H. Y., and Bloom, J. D. (2026). Complete definition of how mutations affect antibodies used to prevent rsv. bioRxiv.

Starr, T. N., Greaney, A. J., Stewart, C. M., Walls, A. C., Hannon, W. W., Veesler, D., and Bloom, J. D. (2022). Deep mutational scans for ace2 binding, rbd expression, and antibody escape in the sars-cov-2 omicron ba.1 and ba.2 receptor-binding domains. PLOS Pathogens, 18(11):1–20.

Tai, L., Zhu, G., Yang, M., Cao, L., Xing, X., Yin, G., Chan, C., Qin, C., Rao, Z., Wang, X., Sun, F., and Zhu, Y. (2021). Nanometer-resolution in situ structure of the sars-cov-2 postfusion spike protein. Proceedings of the National Academy of Sciences, 118(48).

Taylor, A. L. and Starr, T. N. (2026). Deep mutational scanning of recent sars-cov-2 variants highlights changing amino acid preferences within epistatic hotspot residues. bioRxiv.

Thadani, N. N., Gurev, S., Notin, P., Youssef, N., Rollins, N. J., Ritter, D., Sander, C., Gal, Y., and Marks, D. S. (2023). Learning from prepandemic data to forecast viral escape. Nature, 622:818–825.

Transfiguracion, J., Tran, M. Y., Lanthier, S., Tremblay, S., Coulombe, N., Acchione, M., and Kamen, A. A. (2020). Rapid in-process monitoring of lentiviral vector particles by high-performance liquid chromatography. Molecular Therapy Methods & Clinical Development, 18:803–810.

Turakhia, Y., Thornlow, B., Hinrichs, A. S., De Maio, N., Gozashti, L., Lanfear, R., Haussler, D., and Corbett-Detig, R. (2021). Ultrafast sample placement on existing trees (usher) enables real-time phylogenetics for the sars-cov-2 pandemic. Nature Genetics, 53:809–816.

Vaswani, A., Shazeer, N., Parmar, N., Uszkoreit, J., Jones, L., Gomez, A. N., Kaiser, L. u., and Polosukhin, I. (2017). Attention is all you need. In Guyon, I., Luxburg, U. V., Bengio, S., Wallach, H., Fergus, R., Vishwanathan, S., and Garnett, R., editors, Advances in Neural Information Processing Systems, volume 30. Curran Associates, Inc.

Virtanen, P., Gommers, R., Oliphant, T. E., Haberland, M., Reddy, T., Cournapeau, D., Burovski, E., Peterson, P., Weckesser, W., Bright, J., van der Walt, S. J., Brett, M., Wilson, J., Millman, K. J., Mayorov, N., Nelson, A. R. J., Jones, E., Kern, R., Larson, E., Carey, C. J., Polat, İ., Feng, Y., Moore, E. W., VanderPlas, J., Laxalde, D., Perktold, J., Cimrman, R., Henriksen, I., Quintero, E. A., Harris, C. R., Archibald, A. M., Ribeiro, A. H., Pedregosa, F., van Mulbregt, P., and SciPy 1.0 Contributors (2020). SciPy 1.0: Fundamental Algorithms for Scientific Computing in Python. Nature Methods, 17:261–272.

Wallau, G. L., Abanda, N. N., Abbud, A., Abdello, S., Abera, A., Ahuka-Mundeke, S., Falconi-Agapito, F., Alagarasu, K., Ariën, K. K., Ayres, C. F. J., et al. (2023). Arbovirus researchers unite: expanding genomic surveillance for an urgent global need. The Lancet Global Health, 11:e1501–e1502.

Wasserman, W. W. and Sandelin, A. (2004). Applied bioinformatics for the identification of regulatory elements. Nature Reviews Genetics, 5(4):276–287.

Weidenbacher, P. A.-B., Waltari, E., de los Rios Kobara, I., Bell, B. N., Morris, M. K., Cheng, Y.-C., Hanson, C., Pak, J. E., and Kim, P. S. (2022). Converting non-neutralizing sars-cov-2 antibodies into broad-spectrum inhibitors. Nature Chemical Biology, 18(11):1270–1276.

Wolf, T., Debut, L., Sanh, V., Chaumond, J., Delangue, C., Moi, A., Cistac, P., Rault, T., Louf, R., Funtowicz, M., Davison, J., Shleifer, S., von Platen, P., Ma, C., Jernite, Y., Plu, J., Xu, C., Scao, T. L., Gugger, S., Drame, M., Lhoest, Q., and Rush, A. M. (2020). Huggingface’s transformers: State-of-the-art natural language processing.

World Health Organization (2025). WHO TAG-VE risk evaluation for SARS-CoV-2 variant under monitoring: XFG. TAG-VE risk evaluation, World Health Organization, Geneva, Switzerland. Technical Advisory Group on SARS-CoV-2 Virus Evolution (TAG-VE). Accessed 2026-07-27.

Youssef, N., Gurev, S., Ghantous, F., Brock, K. P., Jaimes, J. A., Thadani, N. N., Dauphin, A., Sherman, A. C., Yurkovetskiy, L., Soto, D., Estanboulieh, R., Kotzen, B., Notin, P., Kollasch, A. W., Cohen, A. A., Dross, S. E., Erasmus, J., Fuller, D. H., Bjorkman, P. J., Lemieux, J. E., Luban, J., Seaman, M. S., and Marks, D. S. (2025). Computationally designed proteins mimic antibody immune evasion in viral evolution. Immunity, 58(6):1411–1421.e6.

Yu, T. C., Kikawa, C., Dadonaite, B., Loes, A. N., Englund, J. A., and Bloom, J. D. (2026). Pleiotropic mutational effects on function and stability constrain the antigenic evolution of influenza haemagglutinin. Nature Ecology and Evolution, 10:452–466.

